# A complete reference genome improves analysis of human genetic variation

**DOI:** 10.1101/2021.07.12.452063

**Authors:** Sergey Aganezov, Stephanie M. Yan, Daniela C. Soto, Melanie Kirsche, Samantha Zarate, Pavel Avdeyev, Dylan J. Taylor, Kishwar Shafin, Alaina Shumate, Chunlin Xiao, Justin Wagner, Jennifer McDaniel, Nathan D. Olson, Michael E.G. Sauria, Mitchell R. Vollger, Arang Rhie, Melissa Meredith, Skylar Martin, Joyce Lee, Sergey Koren, Jeffrey A. Rosenfeld, Benedict Paten, Ryan Layer, Chen-Shan Chin, Fritz J. Sedlazeck, Nancy F. Hansen, Danny E. Miller, Adam M. Phillippy, Karen H. Miga, Rajiv C. McCoy, Megan Y. Dennis, Justin M. Zook, Michael C. Schatz

## Abstract

Compared to its predecessors, the Telomere-to-Telomere CHM13 genome adds nearly 200 Mbp of sequence, corrects thousands of structural errors, and unlocks the most complex regions of the human genome to clinical and functional study. Here we demonstrate how the new reference universally improves read mapping and variant calling for 3,202 and 17 globally diverse samples sequenced with short and long reads, respectively. We identify hundreds of thousands of novel variants per sample—a new frontier for evolutionary and biomedical discovery. Simultaneously, the new reference eliminates tens of thousands of spurious variants per sample, including up to 12-fold reduction of false positives in 269 medically relevant genes. The vast improvement in variant discovery coupled with population and functional genomic resources position T2T-CHM13 to replace GRCh38 as the prevailing reference for human genetics.

**One Sentence Summary:** The T2T-CHM13 reference genome universally improves the analysis of human genetic variation.

## Introduction

For the past twenty years, the human reference genome has served as the bedrock of human genetics and genomics [1–3]. One of the central applications of the human reference genome, and reference genomes in general, has been to serve as a substrate for clinical genomics, comparative analyses, and population genomics. More than one million human genomes have been sequenced to study genetic diversity and clinical relationships, and nearly all of them have been analyzed by aligning the sequencing reads from the donors to the reference genome, e.g. [4–6]. Even when the donor genomes are assembled *de novo*, independent of any reference genome, the assembled sequences will nearly always be compared to a reference genome to characterize variation by leveraging the deep catalog of available annotations [7,8]. Consequently, human genetics and genomics benefit tremendously from the availability of a high-quality reference genome, ideally without gaps or errors that may obscure important variation and regulatory relationships.

The current human reference genome, GRCh38, is used for countless applications, with rich resources available to visualize and annotate the sequence across cell types and disease states [3,9–12]. However, despite decades of effort to construct and refine its sequence, the human reference genome suffers from several major limitations that hinder comprehensive analysis. Most immediately, GRCh38 contains more than 100 million nucleotides that either remain entirely unresolved (‘N’ characters), such as the p-arms of the acrocentric chromosomes, or are represented with artificial models, such as the centromeric satellite arrays [13]. Furthermore, GRCh38 possesses 11.5 Mbp of unplaced and unlocalized sequences that are represented outside of the primary chromosomes [3,14]. These sequences are difficult to study, and many genomic analyses exclude them to avoid identifying false variants and false regulatory relationships [6]. Relatedly, reports have persisted of major artifacts that arise when identifying variants relative to GRCh38, such as an apparent imbalance between insertions and deletions (indels) arising from systematic mis-assemblies in GRCh38 [15–17]. Overall, these errors and omissions in GRCh38 introduce biases in genomic analyses, particularly in centromeres, satellites, and other complex regions.

Another major concern is how the reference genome may influence the analysis of variation across large cohorts for population and clinical genomics. Several large studies, such as the 1000 Genomes Project (1KGP) [18] and gnomAD [6], have provided valuable information about the extent of genetic diversity within and between human populations. For example, many analyses of Mendelian and complex diseases use these catalogs of single nucleotide variants (SNVs), small indels, and structural variants (SVs) to rank and prioritize potential causal variants based on allele frequencies (AFs) and other evidence [19–21]. When evaluating these resources, the overall quality and representation of the human reference genome are important, if often overlooked, factors. Any gaps or errors in the sequence could obscure variation and its contribution to human phenotypes and disease. In addition to major omissions such as centromeric sequences or acrocentric chromosome arms, the current reference genome possesses other errors and biases distributed throughout, including within genes of known medical relevance [22,23]. Furthermore, GRCh38 was assembled from multiple donors using clone-based sequencing, which creates an excess of artificial haplotype structures that can subtly bias analyses [1,24]. Over the years, there have been some attempts to replace rare alleles from these haplotypes with more common alleles, but hundreds of thousands of artificial haplotypes and rare alleles remain to this day [25,26]. Increasing the continuity, quality, and representativeness of the reference genome is therefore crucial for improving genetic diagnosis, as well as for understanding the complex relationship between genetic and phenotypic variation.

The recently assembled Telomere-to-Telomere (T2T) CHM13 genome addresses many of the limitations of the current reference [27]. Specifically, the T2T-CHM13v1.0 assembly adds nearly 200 megabases of novel sequence and corrects many of the errors present in GRCh38. Here we demonstrate the impact of the T2T-CHM13 reference on variant discovery and genotyping in a globally diverse cohort. This includes all 3,202 samples from the recently expanded 1000 Genomes Project (1KGP) sequenced with short reads [28] along with 17 samples from diverse populations sequenced with long reads [8,27,29]. From these data, we document universal improvements in read mapping and variant calling with major implications for population and clinical genomics including more than 2 million additional variants found in newly accessible and corrected regions of the genome, 9.2 megabases of corrections found within syntenic regions, and 622 medically-relevant genes showing improved variant calls and interpretation. When combined with the other cellular and genomic resources available for T2T-CHM13, this genome assembly is poised to replace GRCh38 as the predominant reference for human genetics.

## Results

### Structural comparisons of GRCh38 and T2T-CHM13

#### Introducing the T2T-CHM13 genome

The T2T-CHM13 reference genome was primarily assembled from Pacific Biosciences (PacBio) High Fidelity (HiFi) reads augmented with Oxford Nanopore Technology (ONT) reads to close gaps and resolve complex repeats [27]. The resulting T2T-CHM13v1.0 assembly was subsequently validated and polished to exceedingly high quality, with a consensus accuracy estimated to be between Phred Q67 and Q73 [27,30] and with zero known structural defects. The assembly is highly contiguous, with only five unresolved regions from the most highly repetitive ribosomal DNA (rDNA) arrays, representing only 9.9 Mbp of sequence out of >3.0 Gbp of fully resolved sequence. The version 1.0 assembly adds or revises 229 Mbp of sequence compared to GRCh38, defined as regions of the T2T-CHM13 assembly that do not linearly align to GRCh38 over a 1 Mbp interval (i.e., are non-syntenic). Furthermore, 189 Mbp of sequence are not covered by any primary alignments from GRCh38 and are therefore completely novel to the T2T-CHM13 assembly. A summary diagram of the syntenic/non-syntenic regions and their associated annotations are presented in **Fig. 1A** for chromosomes 1 and 21, and a detailed report of all chromosomes is presented in **Fig. S1.1**. Note that the subsequent T2T-CHM13v1.1 assembly [27] further resolves the rDNA regions using model sequences for some array elements, although for this study we analyze the prior v1.0 assembly, which does not contain these representations.

**Figure 1.**
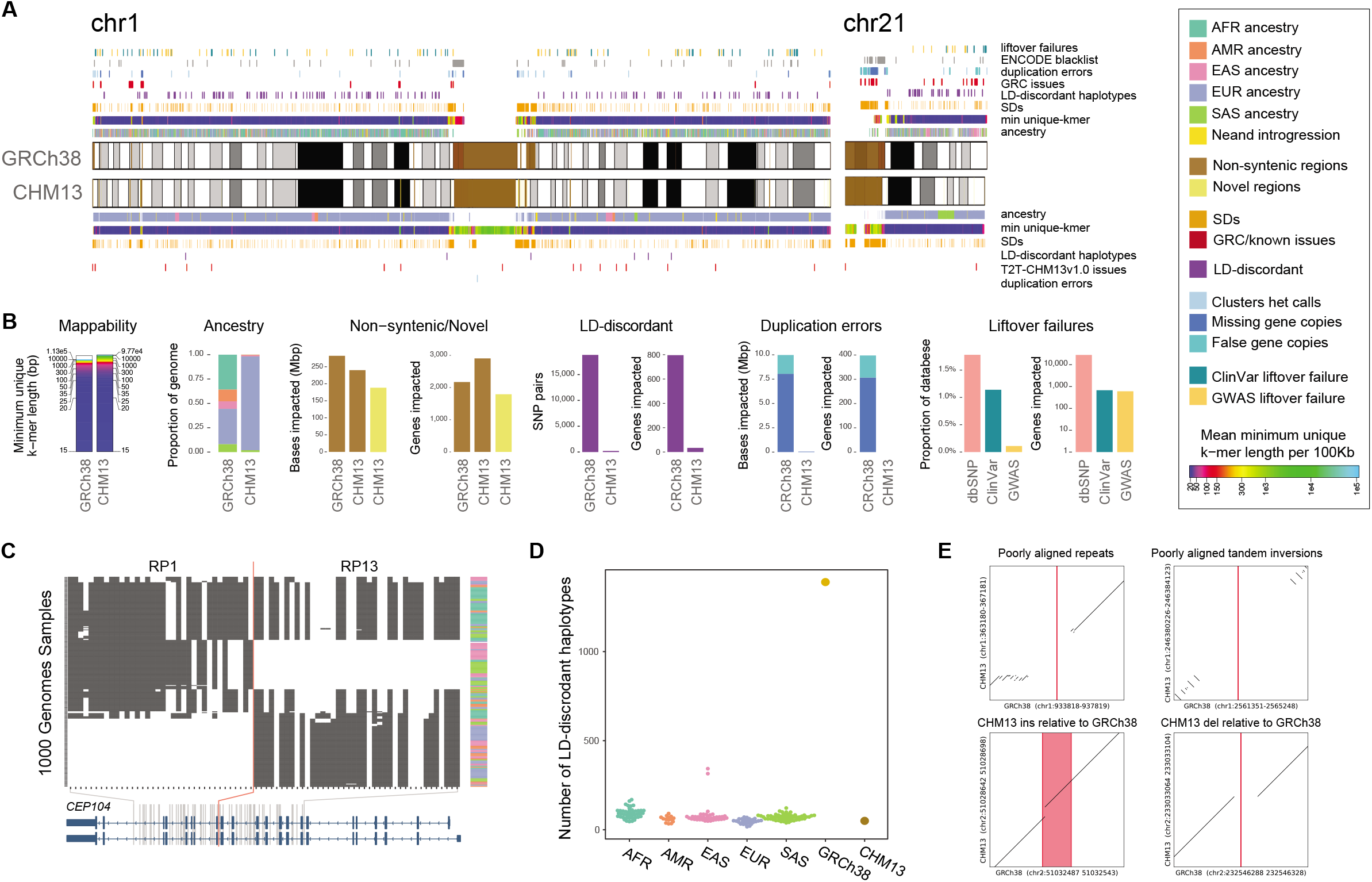
Genomic comparisons of human assemblies GRCh38 and T2T-CHM13. **(A)** Overview of annotations available for GRCh38 and T2T-CHM13 chromosomes 1 and 21 with colors indicated in legend. Cytobands are pictured as gray bands with red bands representing centromeric regions within ideograms. Complete annotations of all chromosomes can be found in Figure S1.1. **(B)** Summary of the number of bases and/or genes annotated by different features for the assemblies with colors indicated in the legend. **(C)** Example of a clone boundary (red line) where GRCh38 possesses a combination of alleles that segregate in negative LD within the 1KGP sample (which we term as an “LD-discordant haplotype”). SNPs are depicted in columns, while phased 1KGP samples are depicted in rows. White indicates reference allele genotypes, while black indicates alternative allele genotypes. Superpopulation ancestry of each sample is indicated in the rightmost column. **(D)** Tally of such LD-discordant haplotypes in a selection of 1000 Genomes samples, as well as GRCh38 and CHM13. **(E)** Examples of variants that cannot be lifted over to T2T-CHM13 because of structural differences between the genomes. The position of the reference allele on GRCh38 is shown in red.

The bulk of the non-syntenic sequence within T2T-CHM13 comprises centromeric satellites (190 Mbp) [31] and new copies of segmental duplications (218 Mbp) [32]. These novel sequences could prove challenging for variant analysis, especially for variants identified using short-read sequencing. However, compared to GRCh38, we report an overall increase of unique sequence as defined by k-length strings (k-mers) found only a single time in the genome (e.g., 14.9 Mbp of newly added unique sequence when considering 50-mers, 23.5 Mbp for 100-mers, and 39.5 Mbp for 300-mers). These unique sequences enable more confident mapping for short-read paired-end sequences or longer reads in some newly uncovered regions of the genome (**Figs. 1B, S1.2, and S1.3**). Regarding highly repetitive regions, more than 106 Mbp of additional sequence was identified in T2T-CHM13 that requires reads of more than 300 bp to uniquely map compared to GRCh38. Concomitantly, there was a decrease in long, exactly duplicated sequences (?5 kbp) shared across chromosomes (excluding sequence pairs within centromeres) from GRCh38 to T2T-CHM13 (**Figs. S1.4 and S1.5**). The former possessed 28 large shared interchromosomal sequences, primarily consisting of pairs of sub-telomeric sequences, with an additional 42 pairs involving at least one unplaced contig. All of these exactly matching sequence pairs, save one between two subtelomeres, are non-identical in T2T-CHM13, as small but important differences between repetitive elements are resolved for the first time.

#### T2T-CHM13 accurately represents the haplotype structure of human genomes

The human reference genome serves as the standard to which other genomes are compared, and is typically perceived as a haploid representation of an arbitrary genome from the population [25]. In contrast with T2T-CHM13, which derives from a single complete hydatidiform mole, the Human Genome Project constructed the current reference genome via the tiling of sequences obtained from bacterial artificial chromosomes (BACs) and other clones with lengths ranging from ~50–150 kbp [24], which derive from multiple donor individuals. GRCh38 and its predecessors thus comprise mosaics of many haplotypes, albeit with a single library (RP11) contributing the majority [24].

To further characterize this aspect of GRCh38 and its implications for population studies, we performed local ancestry inference for both GRCh38 and T2T-CHM13 through comparison to haplotypes from the 1KGP (see **Supplemental Methods**; **Figs. 1A and S1.6**). Continental superpopulation-level ancestry was inferred for 72.9% of GRCh38 clones based on majority votes of nearest-neighbor haplotypes. For the remaining 27.1% of clones, no single superpopulation achieved a majority of nearest neighbors, and ancestry thus remained ambiguous. This ambiguity occurs primarily for short clones with few informative SNPs (**Fig. S1.7**), but also for some longer clones with potential admixed ancestry. In accordance with Green *et al*. [24], we inferred that library RP11, which comprises 72.6% of the genome, is derived from an individual of admixed African-American ancestry, with 56.0% and 28.1% of its component clones assigned to African and European local ancestries, respectively. The second most abundant library, CTD (5.5% of the genome), consists of clones of predominantly (86.3%) East Asian local ancestries, while the remaining libraries are derived from individuals of predominantly European ancestries. In contrast, CHM13 exhibits European ancestries nearly genome wide (**Fig. S1.8**). In addition, GRCh38 and T2T-CHM13 harbor 26.7 Mbp and 51.0 Mbp, respectively, of putative Neanderthal-introgressed sequences that originate from ancient interbreeding between the hominin groups approximately 60 thousand years ago [24]. The excess of introgressed sequence in CHM13, even when restricting to the genomic intervals of GRCh38 clones with confident ancestry assignments, is consistent with its greater proportion of non-African ancestry.

We hypothesized that the mosaic nature of GRCh38 would generate abnormal haplotype structures at clone boundaries, producing many combinations of alleles that are rare or absent from the human population. Indeed, some previous patches of the reference genome sought to correct abnormal haplotype structures in cases where they were noticed due to their impacts on genes of clinical importance (e.g., *ABO* and *SLC39A4*) [3]. Such artificial haplotypes would mimic rare recombinant haplotypes that are private to any given sample, but at an abundance and genomic scale that is unrepresentative of any living human. To test this hypothesis, we identified pairs of common (minor-allele frequency > 10%) autosomal SNP alleles that are always observed on the same haplotype (i.e., segregate in perfect |/C = 1] linkage disequilibrium [LD]) in the 2,504 unrelated individuals of the 1KGP and queried the allelic states of these SNPs in both GRCh38 and T2T-CHM13 (see **Supplemental Methods**).

In accordance with our expectations, we identified numerous haplotype transitions in GRCh38 that are absent from the 1KGP samples, with 18,813 pairs of LD-discordant SNP alleles (i.e., in perfect negative LD) distributed in 1,390 narrow non-overlapping clusters (median length = 3,703 bp) throughout the genome (**Fig. 1C**). Such rare haplotype transitions are comparatively scarce in T2T-CHM13, with only 209 pairs of common high-LD SNPs (50 non-overlapping clusters) possessing allelic combinations absent from the 1KGP sample (**Fig. 1D**). Based on a leave-one-out analysis, we confirmed that T2T-CHM13 possesses a similar number of LD-discordant haplotypes as phased “haploid” samples from 1KGP, whereas GRCh38 vastly exceeds this range (**Fig. 1E**). By intersecting the GRCh38 results with the tiling path of BAC clones, we found that 88.9% (16,733 of 18,813) of discordant SNP pairs straddle the documented boundaries of adjacent clones (**Fig. S1.9**). Of these, 45.9% (7,686 of 16,733) of the clone pairs derived from different BAC libraries, whereas the remainder likely largely reflects random sampling of distinct homologous chromosomes from the same donor individual. Thus, our analysis suggests that T2T-CHM13 accurately reflects haplotype patterns observed in contemporary human populations, whereas GRCh38 does not.

#### T2T-CHM13 corrects systematic errors in GRCh38, including collapsed duplications and falsely-duplicated regions

Genome assemblies often suffer from errors at complex genomic regions such as SDs. In the case of GRCh38, targeted sequencing of BAC clones has been performed to fix many such loci [3,16,33–36], but problems persist. To systematically identify errors in GRCh38 that could produce spurious variant calls, we leveraged the fact that T2T-CHM13 is a haploid-derived cell line that should produce only homozygous variants when its sequence is aligned to GRCh38. Thus, any apparent heterozygous variant can be attributed to (1) mutations accrued in the cell line; (2) sequencing errors; or (3) read mapping errors. In the last case, assembly errors or copy number polymorphism of SDs produce a contiguous stretch of heterozygous variants, which would confound the accurate detection of paralog-specific variants (PSVs). Mapping PacBio HiFi reads from the CHM13 cell line [27] as well as Illumina-like simulated reads (150 bp) obtained from the T2T-CHM13 reference to GRCh38, we identified 368,574 heterozygous SNVs within the autosomes and chromosome X, of which 56,413 (15.3%) were shared among both datasets, evidencing that each technology is distinctively informative due to mappability differences (**Fig. S1.10A and Table S1.1**).

To home in on variants likely deriving from collapsed duplications, we delineated ‘clusters’ of heterozygous calls (see **Methods**) and identified 908 putative problematic regions (541 supported by both technologies) comprising 20.8 Mbp (**Figs. 1 and S1.10B**) with a majority intersecting SD-(668/908; 73.6%) and centromere-associated regions (542/908; 59.7%) [31]. Variants flagged as excessively heterozygous in the population by gnomAD [6] were significantly enriched in these regions (10,000 permutations, empirical *p*-value = 1×10^-4^) representing 26.7% (78,034/292,514) of our CHM13 heterozygous variants—suggesting that these variants arise in genome screens and represent false positives—whilst 341/908 regions (37.55%) overlap with known GRCh38 issues (**Figs. 1A and S1.1**). We next ‘lifted over’ (translated the coordinates of) 821 of these 908 putative problematic regions to the T2T-CHM13 assembly and used human copy number estimates (n=268 individuals from the Simons Genome Diversity Project (SGDP)) [32,37] to conservatively identify 203 loci (8.04 Mbp) evidencing erroneous missing copies in GRCh38 (**Fig. S1.10**). These regions impact 308 gene features with 14 of the total 48 protein-coding genes fully contained within a problematic region, indicating that complete gene homologs are hidden from GRCh38-based population variation analyses as is the case of *DUSP22*, a gene involved in immune regulation [38], and *KMT2C*, a gene implicated in Kleefstra syndrome 2 (OMIM #617768) (**Fig. S1.12**). Additionally, we identified 30 SNPs with known phenotype association (GWAS Catalog) within problematic regions. Finally, we evaluated the status of these regions in the T2T-CHM13 reference by following a similar approach as for GRCh38, obtaining 9,193 heterozygous variants clustered in 11 regions (**Table S1.2**). This revealed that all problematic regions in GRCh38 no longer exhibit clusters of heterozygous variants in T2T-CHM13, enabling accurate variant calling in 48 newly accessible protein-coding genes. We did identify one putative collapsed duplication in T2T-CHM13, based on the presence of a heterozygous variant cluster and reduced copy number in T2T-CHM13, localized to an rDNA array corrected in the most recent version of T2T-CHM13v1.1 [27].

Conversely, we found that the T2T-CHM13 reference also fixed regions falsely duplicated in GRCh38. We identified 12 regions affecting 1.2 Mbp and 74 genes (including 22 protein-coding genes) with duplications private to GRCh38 and not found in T2T-CHM13 or the 268 genomes from SGDP [37] (**Fig. S1.11 and Table S1.3**). In contrast, only five regions affecting 160 kbp have duplications in T2T-CHM13 that are not in GRCh38 or the SGDP. Inspecting the CHM13 data, we deemed that these five loci are true duplications in T2T-CHM13 with strong support from mapped HiFi reads [30]. The five largest duplications in GRCh38, affecting 15 protein-coding genes on the q-arm of chromosome 21, have the falsely duplicated sequence in BAC clones misplaced between many gaps on the heterochromatic p-arm of chromosome 21. The Genome Reference Consortium (GRC) determined that these five clones were incorrectly localized to the acrocentric short arm using admixture mapping and should not have been added to GRCh38 (**Supplemental Section 1**). Of the seven false duplications outside chromosome 21, two are in short contigs between gaps, two are adjacent to a gap, two are on unlocalized “random” contigs, and one is a tandem duplication (**Table S1.4**). A list of falsely duplicated gene pairs in GRCh38 that are corrected in T2T-CHM13 is provided in **Table S1.5.** Using T2T-CHM13 as a reference authoritatively corrects these false duplications, improving variant calling for short- and long-read technologies, including in medically relevant genes, as demonstrated below.

#### Liftover of clinically relevant and trait-associated variation from GRCh38 to T2T-CHM13

In transitioning to a new reference genome, it is imperative to document the locations of known genetic variation of biological and clinical relevance with respect to the updated coordinate system. To this end, we sought to lift over 802,674 unique variants in the ClinVar database and 736,178,420 variants from the NCBI dbSNP database (including 151,876 NHGRI-EBI GWAS Catalog variants) from the GRCh38 reference to the T2T-CHM13 reference. Unambiguous liftover was successful for 789,909 (98.4%) ClinVar variants, 718,683,909 (97.6%) NCBI dbSNP variants, and 103,879 (68.4%) GWAS Catalog SNPs (**Fig. 1A and Table S1.6**), which we provide as a resource for the scientific community within the UCSC Genome Browser and the NHGRI AnVIL, along with lists of all variants that failed liftover and the associated reasons. Critically, this resource includes 138,300 of 138,927 (99.5%) ClinVar variants that were annotated as “pathogenic” or “likely pathogenic.”

Of the 12,765 ClinVar variants that failed to lift over, 9,452 (74.0%) are cases in which the reference and alternative allele are exchanged in T2T-CHM13 versus GRCh38 and are easily recovered. An additional 1,581 liftover failures are caused by scenarios where T2T-CHM13 possessed a third allele that matched neither the reference nor alternative allele of the respective variant. Of the remaining 1,390 ClinVar variants that failed to lift over, 1,186 overlap documented insertions or deletions that distinguish the GRCh38 and T2T-CHM13 assemblies. 342 of these variants are a specialized case of a ref/alt swap (not recoverable with standard liftover tools), in which the ClinVar alternative allele on the GRCh38 reference is an indel and the T2T-CHM13 assembly carries this indel. The remaining 546 variants (< 0.1% of all variants) lie within regions of poor alignment between the GRCh38 and T2T-CHM13 assemblies (**Fig. 1F**). It should be noted that 98.5% of the GWAS Catalog variants that failed to lift over were cases in which the position of the variant exists in the T2T-CHM13 assembly, but the reference allele in T2T-CHM13 does not match the reference allele in GRCh38. The modes of liftover failure for variants in dbSNP and the GWAS Catalog otherwise follow similar distributions and are presented in **Table S1.6.** In all, these annotated variants represent a critical resource enabling researchers to effectively interpret future genetic results using the T2T-CHM13 assembly.

### T2T-CHM13 improves analysis of global genetic diversity based on 3,202 short-read samples from the 1KGP dataset

#### T2T-CHM13 improves mapping of3,202 short-read samples from the 1KGP dataset

To investigate how the T2T-CHM13 assembly impacts short-read variant calling, we realigned and reprocessed all 3,202 samples from the recently expanded 1KGP cohort [28] using the NHGRI AnVIL Platform [39]. In this collection, each sample is sequenced to at least 30x coverage using paired-end Illumina sequencing, with samples from 26 diverse populations across 5 major continental superpopulations (**Fig. S2.1**). Most samples are unrelated, although the expanded collection includes 602 complete trios that we use to estimate the rate of inaccurate (non-Mendelian) variants below. We matched the previously used analysis pipeline for GRCh38 [28] as closely as possible so that any major differences would be attributable to the reference genome rather than technical differences in the analysis software (**Supplemental Methods**).

On average, an additional 7.4 × 10^6^(0.97%) of properly paired reads map to T2T-CHM13 rather than GRCh38 using BWA-MEM [40], even when considering the alt and decoy sequences used by the original analysis project (**Fig. S2.4**). Interestingly, even though more reads align to T2T-CHM13, the observed per-read mismatch rate after alignment to T2T-CHM13 is 20% to 25% smaller for all continental populations, although African samples continue to present the highest mismatch rate (**Fig. 2A**). This is expected because the observed mismatch rate counts both genuine sequencing errors, which are largely consistent across all samples, and any true biological differences between the read and the reference genome, which vary substantially based on the ancestry of the sample. Relatedly, T2T-CHM13 improved several other mapping characteristics, such as a decline in the number of mis-oriented read pairs (**Fig. 2A**). Finally, by considering the alignment coverage across 500 bp bins across the respective genomes, we observed improved coverage uniformity within every sample’s genome when using T2T-CHM13 rather than GRCh38. For example, within gene regions, we noted a 4-fold decrease in the standard deviation of the coverage (**Fig. 2A**) and observed similar improvements in other types of genomic regions among all population groups (**Fig. S2.5**). Overall, these improvements in error rates, mapping characteristics, and coverage uniformity demonstrate the superiority of T2T-CHM13 as a reference genome for short-read alignment across all populations.

**Figure 2.**
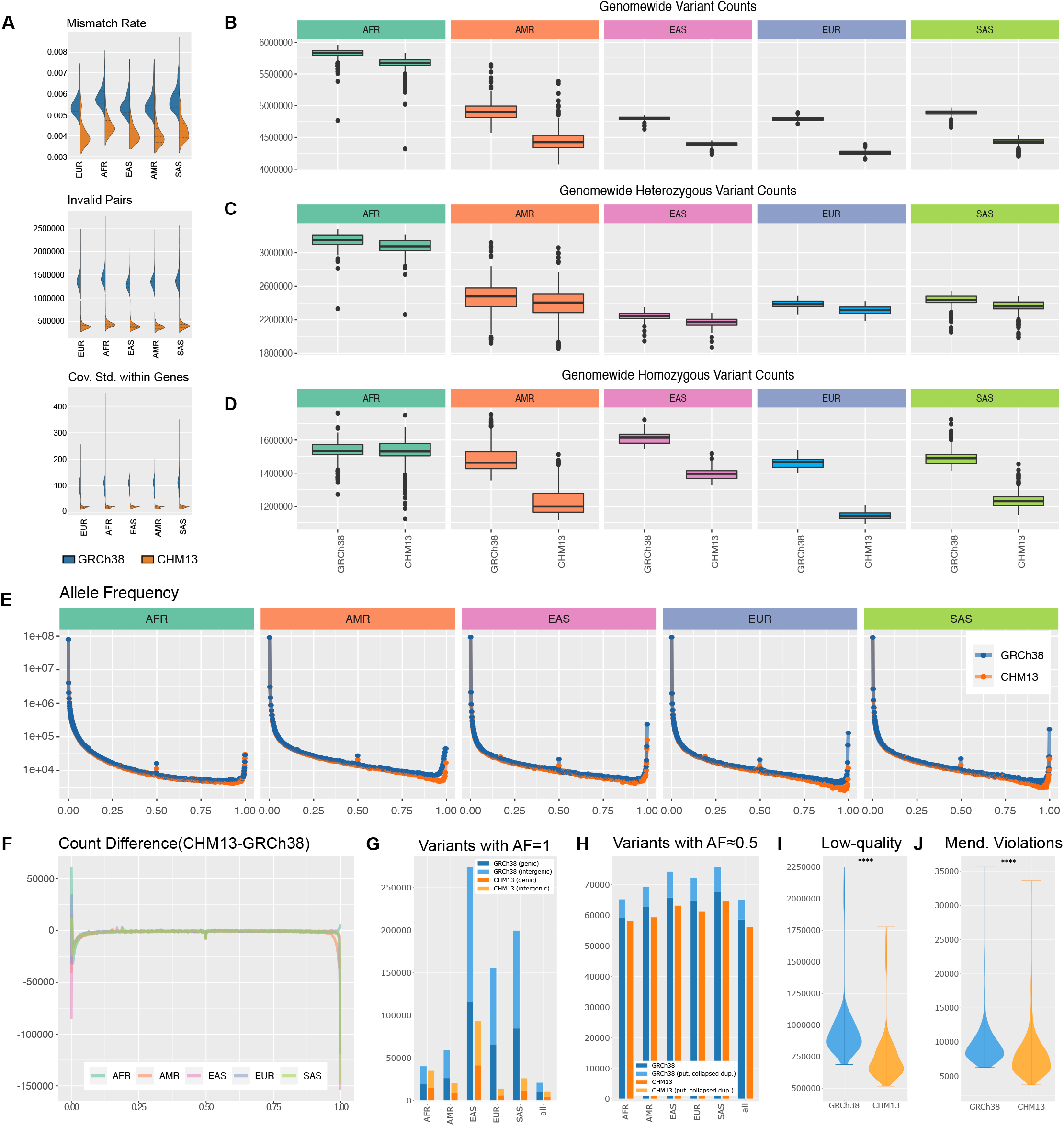
Improvements to Short-Read Mapping and Variant Calling. **(A)** Summary of alignment characteristics aligning to CHM13 instead of GRCh38. **(B)** Boxplot of overall number of variants found in each person across superpopulations. **(C)** Boxplot of the number of heterozygous variants found in each person across superpopulations. **(D)** Boxplot of the number of homozygous variants found in each person across superpopulations. **(E)** Allele frequency distribution of each superpopulation relative to CHM13 and GRCh38. **(F)** Change in allele frequency distribution. **(G)** Number of variants with allele frequency equal to 100%, both within protein-coding genes and without. **(H)** Number of variants with allele frequency equal to 50%, both within putative collapsed duplications and without. **(I)** Violin plot of the number of low-quality variants found when aligning to GRCh38 and CHM13. **(J)** Violin plot of the number of Mendelian violations found when aligning to GRCh38 and CHM13.

#### T2T-CHM13 improves variant calling across populations

From these alignments, we next evaluated SNV and small indel variant calls with the widely-used GATK Haplotype Caller, which uses a joint genotyping approach to optimize accuracy across large populations [41]. Again, we matched the pipeline used in the prior 1KGP study, albeit with newer versions of some analysis tools, to minimize software discrepancies and attribute differences to changes in the reference genome. Across all samples, we identified 57,315,786 high-quality (“PASS”) variants relative to T2T-CHM13 (per-sample mean: 4,717,525; median: 4,419,802) compared to 59,051,733 relative to GRCh38 (per-sample mean: 5,101,897; median: 4,867,871), additionally noting a decrease in the number of called variants per individual genome (**Fig. 2B**). We performed all subsequent analyses using these high-quality variants, as the PASS filter successfully removed spurious variants (**Fig. S2.6**), particularly in complex regions (**Fig S2.7**).

As with the improvement to the per-read mismatch rate, we attribute the reduction in the number of variant calls due to improvements in the number of rare alleles, consensus errors, and structural errors in T2T-CHM13 versus GRCh38. This conclusion is supported by the observation that the number of heterozygous variants per sample is more similar (**Fig. 2C**) across reference genomes in contrast to homozygous variants (**Fig. 2D**). This discrepancy is especially pronounced in non-African samples, which have substantially more homozygous variants relative to GRCh38 than T2T-CHM13. This is likely because ~70% of the GRCh38 sequence comes from an individual with African-American ancestry, and African populations are enriched for rare and private variants [18].

Further investigating this relationship, we computed the AFs of variants from unrelated samples from each of the five continental superpopulations (**Fig. 2E**). The distributions were nearly equivalent over most of the AF spectrum, but substantial differences were observed for rare alleles (AF < 0.05), intermediate-frequency alleles, including errors where nearly all individuals are heterozygous (AF ≅ 0.5), and fixed/nearly-fixed alleles (AF > 0.95). The most prominent difference in AF distributions affected fixed or nearly-fixed alleles in each assembly, for which all non-African superpopulations showed an excess of ~150,000 variants in GRCh38, while the African superpopulation showed an excess of 2,364 variants in T2T-CHM13 (**Fig. 2F**). This observation is driven by a striking decrease in the number of completely fixed variants (100% AF) relative to GRCh38 (**Fig. 2G**). Such variants represent positions where the reference genome itself is the only sample observed to possess the corresponding allele. These alleles arise either because of genuine private variants in one of the GRCh38 donors, or from sequencing errors in the reference genome itself, and result in 100% of other individuals possessing two copies of the alternative allele. As a result, these ‘variants’ will not be reported at all if the same reads are mapped to a different genome that does not have these private alleles. In addition to this lower rate of private variation, T2T-CHM13 possesses fewer ultra-rare variants, effectively reducing the number of “nearly fixed” alleles in population data such as 1KGP. Finally, the reduction in AF ≅ 0.5 variants is largely explained by the corrections to collapsed SDs (**Table S1.1**), as these regions are highly enriched for heterozygous PSVs in nearly all individuals caused by the false pileup of reads from the duplicated regions to a single location (**Fig. 2H**). Collectively, the decrease in variants with AF = 1 and AF ≅ 0.5 largely explains the decrease in the overall number of variants observed per sample and across the entire population for T2T-CHM13.

Informed by these results, we considered the feasibility of using the T2T-CHM13 reference to call variants and then lifting over the results to GRCh38 for further analyses. Using a liftover tool to transform a variant call set for a single sample into a call set with respect to GRCh38 requires special handling to account for variants for which the two references have different alleles. Specifically, if one of the reference alleles is not present in the sample, it will be necessary to genotype the site against the new reference. Although this issue is less of a concern for large datasets like the 1KGP, even these large samples will contain a small number of variants that become invisible when switching reference genomes (**Fig. S2.10**). In addition, differences in variant representation, especially in regions of low complexity, may cause lifted variant sets to differ from those called against the target reference.

#### Significant reduction of genome-wide Mendelian violations

As further quality control for the variant calls, we performed a Mendelian concordance analysis using the 602 trios represented in the 1KGP cohort. We observed a statistically significant decrease in both the number of low-quality variants (median: 890,701 (GRCh38) vs. 682,609 (T2T-CHM13), *p*-value = 4.943 × 10^-96^, Wilcoxon signed-rank test) (**Fig. 2I**) and the number of Mendelian violations (median: 8,879 (GRCh38) vs. 7,484 (T2T-CHM13), *p*-value = 7.346 × 10^-96^, Wilcoxon signed-rank test) (**Fig. 2J**) when aligned to T2T-CHM13 as compared to GRCh38. In addition to providing an estimate of the error rate for variant calls in this callset, this improvement has broad implications for clinical genetics analyses of *de novo* or somatic mutations, which have been implicated as causes of autism spectrum disorders [42] and many forms of cancer [43].

### T2T-CHM13 improves structural variant analysis of 17 diverse long-read samples

#### T2T-CHM13 improves mapping of 17 long-read samples

Next, we investigated the effects of using T2T-CHM13 as a reference genome for alignment and large SV calling from both PacBio HiFi and ONT long reads. To this end, we aligned reads and called SVs in seventeen samples of diverse ancestry from the Human Pangenome Reference Consortium (HPRC+) [27] and the Genome in a Bottle Consortium (GIAB) [29], including two trios (**Table 3.1**). All of these samples had HiFi data available, and fourteen had also been sequenced with ONT (**Fig. 3A**), with mean read lengths of 18.1 kbp and 21.9 kbp and read N50 values of 18.3 kbp and 44.9 kbp, respectively (**Fig. S3.1**).

**Figure 3.**
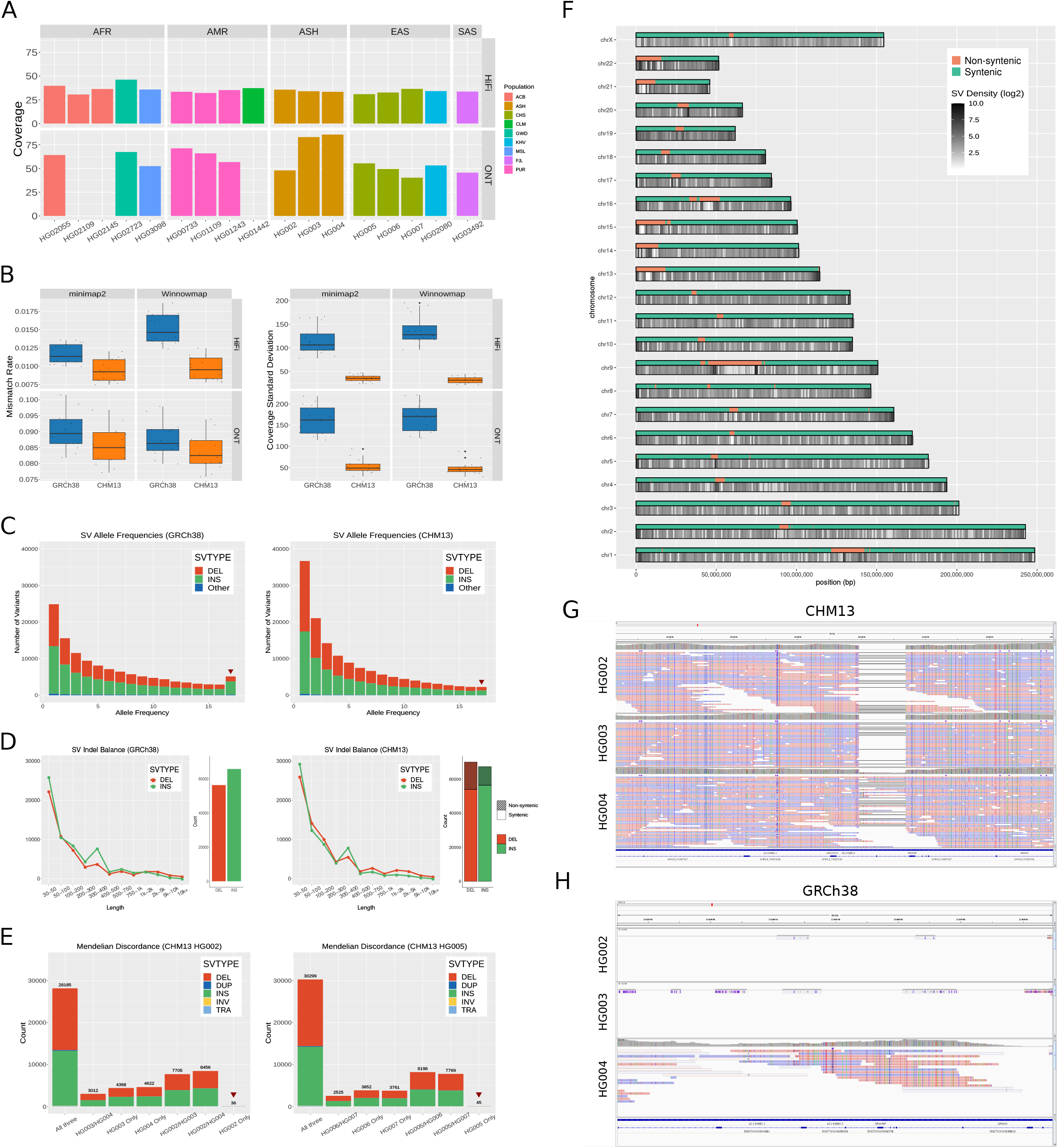
Improvements to Long-Read Alignment and SV Calling in CHM13. **(A)** The coverage, ancestry, and sequencing platforms available for the 17 samples sequenced with long reads. **(B)** The genome-wide mapping error rate and the standard deviation of the coverage for CHM13 and GRCh38. The standard deviation was computed across each 500bp bin of the genome. **(C)** The allele frequency of SVs derived from HiFi data in CHM13 and GRCh38 among the 17-sample cohort. The red arrows indicate uniform (100% frequency) variants. **(D)** The balance of insertions vs. deletion calls in the 17-sample cohort in CHM13 and GRCh38. Variants in CHM13 are stratified by whether or not they intersect regions which are non-syntenic with GRCh38. **(E)** The SV calls in CHM13 for two trios: a trio of Ashkenazi ancestry (child HG002, and parents HG003 (46XY), and HG004 (46XX), and a trio of Han Chinese ancestry (child HG005, and parents HG006 (46XY) and HG007 (46XX)). The red arrows indicate child-only, or candidate de novo, variants. **(F)** The density of SVs called from HiFi data in the 17-sample cohort across CHM13. **(G)** Alignments of HiFi reads in the HG002 trio to CHM13 showing a deletion spanning an exon of the transcript AC134980.2. **(H)** Alignments of HiFi reads in the HG002 trio to the same region of GRCh38 as shown in (g), showing much poorer mapping to GRCh38 than to CHM13.

In line with our short-read results, aligning long reads to T2T-CHM13 compared to GRCh38 did not substantially change the number of reads mapped using either Winnowmap [44] or minimap2 [45] because most of the novel sequence in T2T-CHM13 represents additional copies of SDs or satellite repeats already partially represented in GRCh38 (**Fig. S3.2**). However, aligning to T2T-CHM13 reduced the observed per-read mapping mismatch rates by 5% to 40% across the four combinations of sequencing technologies and aligners because GRCh38 has more rare alleles. T2T-CHM13 also corrects errors in GRCh38 and is a complete assembly of the genome, which helps map all reads to the correct loci, similar to what we observed for short reads (**Fig. 3B**). To study coverage uniformity, we next measured the average coverage across each 500-bp bin on a per-sample basis and computed the standard deviation of the coverage among all bins. Across all aligners and technologies, the median standard deviation of the per-bin coverage was reduced by more than a factor of three, indicating more stable mapping to T2T-CHM13 (**Fig. 3B**). This difference in coverage uniformity was particularly pronounced in satellite repeats and other regions of GRCh38 non-syntenic with T2T-CHM13 (**Fig. S3.3**). The more uniform coverage will broadly improve variant calling and other long-read based analysis.

#### T2T-CHM13 fixes SV imbalances on GRCh38

We next used our optimized SV-calling pipeline, including Sniffles [46], Iris, and Jasmine [47], to call SVs in all 17 samples and consolidate them into a cohort-level callset in each reference from HiFi data. From these results, we observe a reduction from 5,147 to 2,260 SVs that are homozygous in all 17 individuals when calling variants relative to T2T-CHM13 instead of GRCh38 (**Fig. 3C**). Previous studies [16,17] have noted the excess of such SV calls when using GRCh38 as a reference and attributed them to structural errors; the use of a complete and accurate reference genome predictably reduces the number of such variants. In addition, the number of indels is more balanced when calling against T2T-CHM13, whereas GRCh38 exhibited a bias towards insertions caused by missing or incomplete sequence (**Fig. 3D**), such as incorrectly collapsed tandem repeats [16]. With respect to T2T-CHM13, we observe a small bias towards deletions, which likely results from the challenges in calling insertions with mapping-based methods and in representing SVs within repeats, as this difference is especially prominent in highly repetitive regions such as centromeres and satellite repeats (**Fig. S3.8**). The variants we observe relative to T2T-CHM13 are enriched in the centromeres and sub-telomeric sequences (**Fig. 3F**), likely because of a combination of repetitive sequence and greater recombination rates [17]. We observe similar trends among SVs unique to single samples (**Fig. S3.11**).

We also observe similar improvements in the insertion/deletion balance for large SVs >500 bp detected by Bionano optical mapping data in HG002 against the T2T-CHM13 reference, with an increase in deletions from 1,199 to 1,379 and a decrease in insertions from 2,771 to 1,431 on GRCh38 and T2T-CHM13, respectively (**Fig. S3.14**). Using the T2T-CHM13 reference for Bionano also enables improved SV calling around gaps in GRCh38 that are closed in T2T-CHM13 (**Fig. S3.15**), suggesting that T2T-CHM13 offers improved indel balance compared to GRCh38 across multiple SV-calling methods.

#### De novo SV analysis within trios

To investigate the impacts of the T2T-CHM13 reference on our ability to accurately detect *de novo* variants, we called SVs in both of our trio datasets using a combination of HiFi and ONT data and identified SVs only present in the child of the trio and supported by both technologies—about 40 variants per trio (**Fig. 3E**). Manual inspection revealed a few strongly-supported variants in each trio with consistent coverage and alignment breakpoints, while the other candidates had less reliable alignments as seen in previous reports [47]. In HG002, we detected six strongly-supported candidate *de novo* SVs that had been previously reported [29,47]. In HG005, we detected a novel 1,571 bp deletion at chr17:49401990 in T2T-CHM13 that is strongly supported as a candidate *de novo* SV relative to both T2T-CHM13 and GRCh38 (**Fig. S3.9**). This demonstrates the ability of T2T-CHM13 to be used as a reference genome for *de novo* SV analysis.

#### T2T-CHM13 enables the identification of novel SVs

The improved accuracy and completeness of the T2T-CHM13 genome helps resolve complex genomic regions where reads fail to align to GRCh38. Within non-syntenic regions, we identified a total of 27,055 SVs (**Fig. 3D**), the majority of which were deletions (15,998) and insertions (10,912). 22,362 of these SVs (82.7%; 8,903 insertions; 13,334 deletions) overlap novel sequences in T2T-CHM13, while the remaining novel SVs are newly accessible because of the improved accuracy of the T2T-CHM13 reference. The AF and size distributions for these variants mirror the characteristics of the syntenic regions, with rare variants (**Fig. S3.10**) and smaller 30-50 bp indels (**Fig. S3.7**) being the most abundant. However, we also note some non-syntenic regions with relatively few or zero SVs identified. While many of these regions lie at the interiors of p-arms of acrocentric centromeres, which are gaps in T2T-CHM13v1.0 that have been resolved in later versions of the assembly (**Fig. 3F**), we also noticed depletions of SVs in a few other highly repetitive regions, such as the newly resolved human satellite array on chromosome 9. We largely attribute the reduction in variant density to the low mappability of these complex and repetitive regions. Future improvements in read lengths and alignment algorithms are needed to further resolve such loci.

Furthermore, within syntenic regions, we also note improvements to alignment and variant calling accuracy, including the identification of strongly-supported variant calls not previously observed within homologous regions of GRCh38. For example, in T2T-CHM13 we observe a deletion in all of the samples of the HG002 trio in an exon of the olfactory receptor gene *AC134980.2* (**Fig. 3G**), while the reads from those samples largely fail to align to the corresponding region of GRCh38 (**Fig. 3H**). Meanwhile, reads from African samples (**Fig. S3.13**) align to both references at this locus. The difference in alignment among different samples is likely due to the region being highly polymorphic for copy number variation; GRCh38 contains a fair representation of that region for the tested African samples, while the homologous region in T2T-CHM13 more closely resembles European samples (**Fig. S3.12**). This highlights the need for T2T reference genomes for as many diverse individuals as possible to capture these major haplotype differences.

### Novel variants and evolutionary signatures within newly accessible regions of the genome

#### T2T-CHM13 newly enables variant calling in novel and corrected regions of the genome

The T2T-CHM13 genome contains 229 Mbp of sequence that is non-syntenic to GRCh38, which intersects 207 protein-coding genes (**Figs. 4A and 4B**). Within these regions, we report a total of 2,050,129 PASS variants across all 1KGP samples based on short reads (**Figs. 4A and 4C**). Comparing variants called in a subset of 14 HPRC+ samples with Illumina, HiFi, and ONT data, we found that 73–78% of the Illumina-discovered SNVs are concordant with variants identified with PacBio HiFi long-read data using the PEPPER-Margin-DeepVariant algorithm (51,306–74,122 matching SNVs and genotypes per sample) [48]. Long reads discover over ten times more SNVs per sample than short reads in these regions, with 447,742–615,085 (41–43%) of SNVs matching between HiFi and ONT with PEPPER-Margin-DeepVariant.

**Figure 4.**
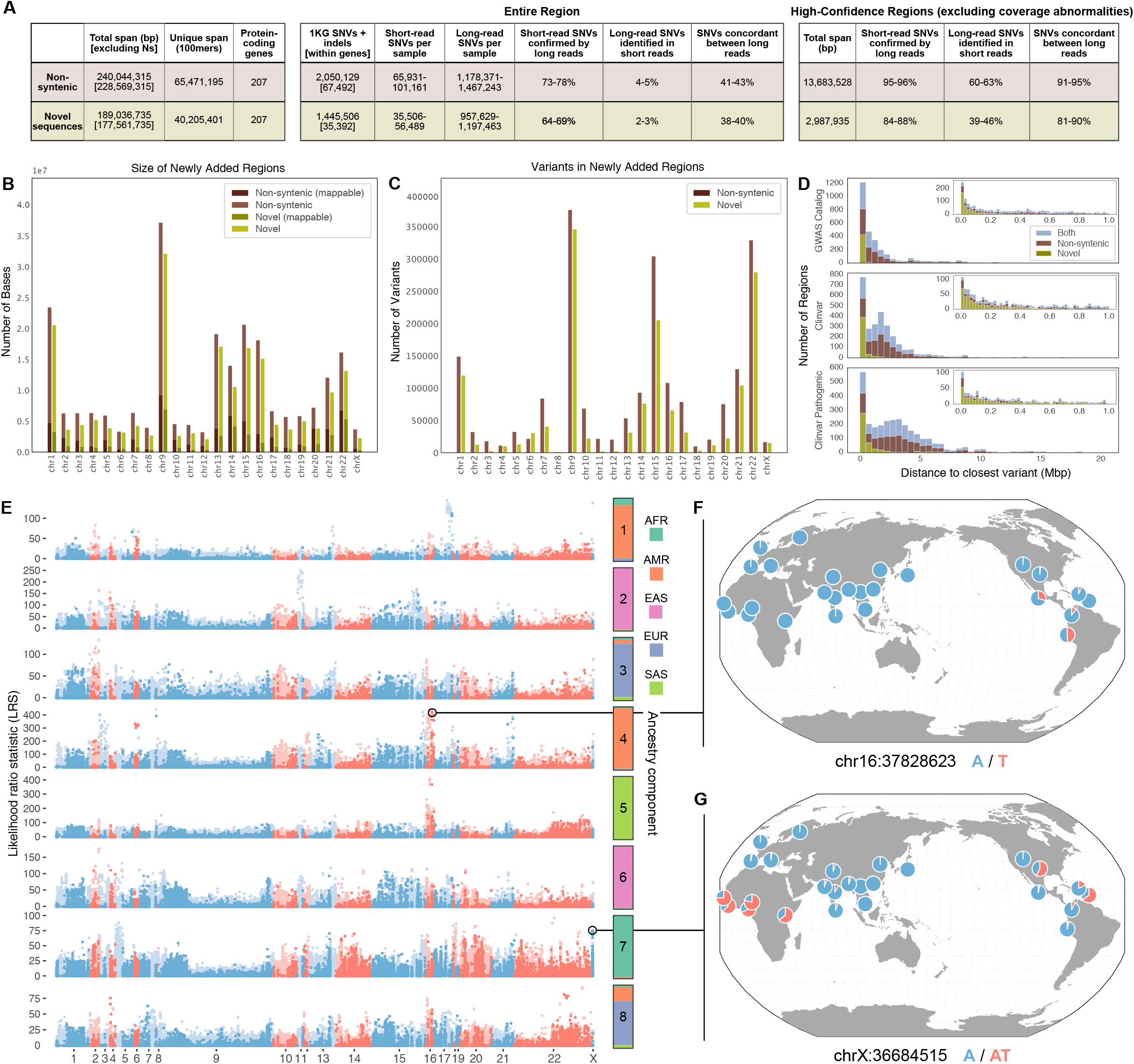
Characterization of variants within newly accessible regions of the genome. **(A)** Overview of non-syntenic and novel regions and their respective variant counts. **(B)** Number of bases added in non-syntenic and novel regions by chromosome, along with how many variants for each respective region are mappable (have contiguous unique 100mers). **(C)** Number of variants in these newly added regions by chromosome. **(D)** Distance from each novel-only, non-syntenic-only, or overlapping region to the closest Clinvar or GWAS Catalog variant. Insets are zoomed to 1 Mbp. **(E)** Scan for variants in non-syntenic (light blue and red) and novel (dark blue and red) regions that exhibit extreme patterns of allele frequency differentiation. Allele frequency outliers were identified for each of eight ancestry components, colored by the superpopulation membership of the corresponding 1000 Genomes samples. Large values of the likelihood ratio statistic (LRS) denote variants for which AF differences in the corresponding ancestry component exceeds that of a null model based on genome-wide covariances in allele frequencies. **(F, G)** Population-specific allele frequencies of two highly differentiated variants in novel regions.

We further define conservative higher-confidence regions by excluding regions with abnormal coverage in any long-read sample (i.e., coverage falling outside of 1.5x the interquartile range). This effectively excludes difficult-to-map regions with excessively repetitive alignments as well as copy number variable regions. After excluding abnormal coverage from non-syntenic regions, 14 Mbp remain, and SNVs from HiFi and ONT long reads are 91–95% concordant (21,835–28,237 variants). 95-96% (14,575–18,949) of short-read SNVs are found in HiFi long-read calls, though 37–40% of HiFi SNVs are still missing from the short-read calls due to poorer mappability of the short-reads (**Table S4.1**). While many non-syntenic regions will require further method development to achieve accurate variant calls, the concordance of long- and short-read calls for tens of thousands of variants highlights novel sequences that are immediately accessible to both technologies.

As these broadly-defined non-syntenic regions include inversions and other large structural changes between GRCh38 and T2T-CHM13 that do not necessarily alter many of the variants contained within, we also considered a narrower class of novel sequences, defined as segments of the T2T-CHM13 genome that do not align to GRCh38 using Winnowmap [44]. Within these novel sequences, which span a total of 189 Mbp (**Figs. 4A, 4B, S4.3**), we report a total of 1,445,506 PASS variants in 1KGP samples based on short reads, intersecting 207 protein-coding genes (**Figs. 4A, 4C, S4.1**). Note that this set of 207 genes is distinct from the 207 genes that intersected with the non-syntenic regions, and the two sets together comprise 329 unique genes. Because these novel sequences are enriched for highly repetitive sequences, concordance is slightly lower, such that 64–69% of the SNVs in each sample match variants found in PacBio HiFi long-read data from the same samples (24,371–36,501 matching SNVs and genotypes per sample), and 339,783–473,074 (38–40%) of SNVs match between HiFi and ONT. When removing difficult-to-map and copy-number-variable regions as above, 3 Mbp of high-confidence regions remain. Within high-confidence regions, 84–88% of short-read SNVs in each sample match variants found in each sample’s PacBio HiFi long-read data (2,938–3,811 matching SNVs and genotypes per sample), and 5,544–8,298 (81–90%) of SNVs match between HiFi and ONT (**Table S4.1**). While these novel regions are more challenging than non-syntenic regions, thousands of new variants can be called concordantly using short and long reads.

We noted homology between GRCh38 collapsed duplications and many T2T-CHM13 non-syntenic and/or novel regions (137 regions comprising 6.8 Mbp), indicating that these sequences are potentially corrected in the new assembly through the deconvolution of nearly identical repeats. Comparing total variants identified in the 1KGP dataset, we observed a significant decrease in variant densities of 41 protein-coding genes intersecting with GRCh38 collapsed duplications in T2T-CHM13 (mean: 27 variants per kbp) compared with GRCh38 (mean: 46 variants per kbp; *p-value* = 6.906 x 10^-8^, Wilcoxon signed-rank test) (**Fig. S4.2**). Besides differences in local ancestries between the references, these higher variant densities in GRCh38 in part represent PSVs or misassigned alleles from missing paralogs. Conversely, 1KGP variants were significantly increased in 32 protein-coding genes contained within GRCh38 false duplications using the new corrected reference genome (mean values of 48 variants per kbp in T2T-CHM13 vs. 12 variants per kbp in GRCh38; *p-value* = 4.657 x 10^-10^). To assess if these corrected complex regions in T2T-CHM13 correctly identify variation, we evaluated the concordance of variants generated from short-read Illumina and PacBio HiFi sequencing datasets of two trios from the GIAB consortium and the Personal Genome Project [49] and observed similar recall for Illumina data in T2T-CHM13 (20.1–28.3%) and GRCh38 (21.5–25.4%), but with significantly improved precision in the variants identified (98.1–99.7% in T2T-CHM13 vs. 64.3–67.3% in GRCh38) in a subset of the GRCh38 collapsed duplications (copy number < 10; ~910 kbp) (**Table S4.2**). Corrected false duplications (1.2 Mbp) exhibit dramatically improved recall for Illumina data compared with HiFi in T2T-CHM13 (57.4–68.3%) vs. GRCh38 (1.1–1.8%), as well as improved precision in T2T-CHM13 (98.5–99.3%) vs. GRCh38 (76.5–95.8%) (**Table S4.2**). These improvements show that variants can now be more reliably discovered and genotyped in regions and genes corrected by this new assembly.

#### Phenotypic associations and evolutionary signatures within newly accessible regions

Newly accessible sequences in the T2T-CHM13 assembly that are non-syntenic with GRCh38 offer a new frontier for genetic studies. Many such loci lie in close proximity to variation that has been implicated in complex phenotypes or disease, supporting their potential biomedical importance. These include 8 loci occurring within 10 kbp of GWAS hits and 19 loci within 10 kbp of ClinVar pathogenic variants (**Fig. 4D**). In addition, 113 unique **GWAS** hits (representing 0.5% of all variants in studies we tested) segregate in LD (*R*^2^ ≥ 0.5) with variants in non-syntenic regions, thereby expanding the catalog of potential causal variants for these studies [50] (**Fig. S4.4**).

Using short-read-based genotypes generated from the 1KGP cohort, we also searched for variants with large differences in AF between populations, a signature that can reflect historical positive selection or demographic forces. To study these signatures, we applied Ohana [51], a method that models individuals as possessing ancestry from *k* components and tests for ancestry component-specific frequency outliers. Focusing on continental-scale patterns (*k* = 8; **Fig. S4.5**), we identified 5,154 unique SNVs and indels across all ancestry components that exhibited strong deviation from genome-wide patterns of AF (99.9th percentile of distribution for each ancestry component; **Fig. 4E**). These included 814 variants overlapping with annotated genes, and 195 variants that intersected annotated exons.

We first focused on the 3,038 highly differentiated variants that lift over from T2T-CHM13 to GRCh38. These successful lift overs allowed us to make direct comparisons to selection results, generated with identical methods, based on 1KGP Phase 3 data aligned to GRCh38 (see **Supplemental Methods**; **Fig. S4.6**) [52]. For 41.3% of the lifted over variants, we found GRCh38 variants within a 2 kbp window that possessed similar or higher likelihood ratio statistics for the same ancestry component, indicating that these loci were possible to identify in previous scans based on GRCh38 (**Fig. S4.7**). Conversely, our results suggest that the remaining 58.7% of lifted over variants, while not “novel” with respect to GRCh38, may represent regions of the genome where T2T-CHM13 improves variant calling enough to resolve previously unknown signatures of AF differentiation (**Fig. S4.8**).

We then investigated the 943 variants that could not be lifted over from T2T-CHM13 to GRCh38 and were located in T2T-CHM13 novel sequences, as well as regions deemed mappable based on unique 100-mer analysis. Some of these variants overlap with genes, including several that are newly annotated on T2T-CHM13 based on RNA transcripts (**Fig. 4E and Table S4.3**). Here, we highlight two loci that exhibit some of the strongest allele frequency differentiation that we observed across ancestry components. The first locus, located in a centromeric alpha satellite on chromosome 16, contains variants that reach intermediate allele frequency in the ancestry component corresponding to the Peruvian in Lima, Peru (PEL) and other Admixed American populations of 1KGP (AFs: 0.49 in PEL; 0.20 in CLM [Colombian in Medellin, Colombia] and MXL [Mexican Ancestry in Los Angeles, California]; absent or nearly absent elsewhere; **Figs. 4F, S4.9, S4.10**). Variants at the second locus, which is located in a novel sequence on the X chromosome that contains a multi-kbp imperfect AT tandem repeat, exhibit high AFs in the ancestry component corresponding to African populations of 1KGP and low AFs in other populations (AFs: 0.67 in African populations and 0.014 in European populations; **Figs. 4G, S4.11, S4.12**). The variant at this locus with the strongest signature of frequency differentiation also lies within 10 kbp of two pseudogenes, *MOB1AP2-201* (MOB kinase activator 1A pseudogene 2) and *BX842568.1-201* (ferritin, heavy polypeptide-like 17 pseudogene).

We note that due to the repetitive nature of the sequences in which they reside, many of the loci that we highlight here remain challenging to genotype with short reads, and individual variant calls remain uncertain. Nevertheless, patterns of AF differentiation across populations are relatively robust to such challenges, and can still serve as proxies for more complex SVs whose sequences cannot be resolved by short reads alone. The presence of population-specific signatures at these loci highlight the potential for T2T-CHM13 to reveal new evolutionary signals in previously unresolved regions of the genome.

### Impact of T2T-CHM13 on Clinical Genomics

#### Potentially clinically relevant variants in T2T-CHM13

A deleterious variant in a reference genome can mislead the interpretation of a clinical variant identified in a patient because it may not be flagged as such using standard analysis tools. The GRCh38 reference genome is known to contain such variants that likely affect gene expression, protein structure, or protein function [25], though systematic efforts have sought to identify and remove these alleles [3]. To determine the existence and location of loss-of-function variants in T2T-CHM13, we aligned the assembly to GRCh38 using dipcall [53] to identify and functionally annotate nucleotide differences [54] (**Fig. 5A**). This identified 210 putative loss-of-function variants (defined as variants impacting protein-coding regions and predicted splice sites) impacting 189 genes, 31 of which are clinically relevant [23]. These results are in line with previous work showing that the average diploid human genome contains ~450 putative loss-of-function variants impacting ~200–300 genes using low-coverage Illumina sequencing (before stringent filtering) [55].

**Figure 5.**
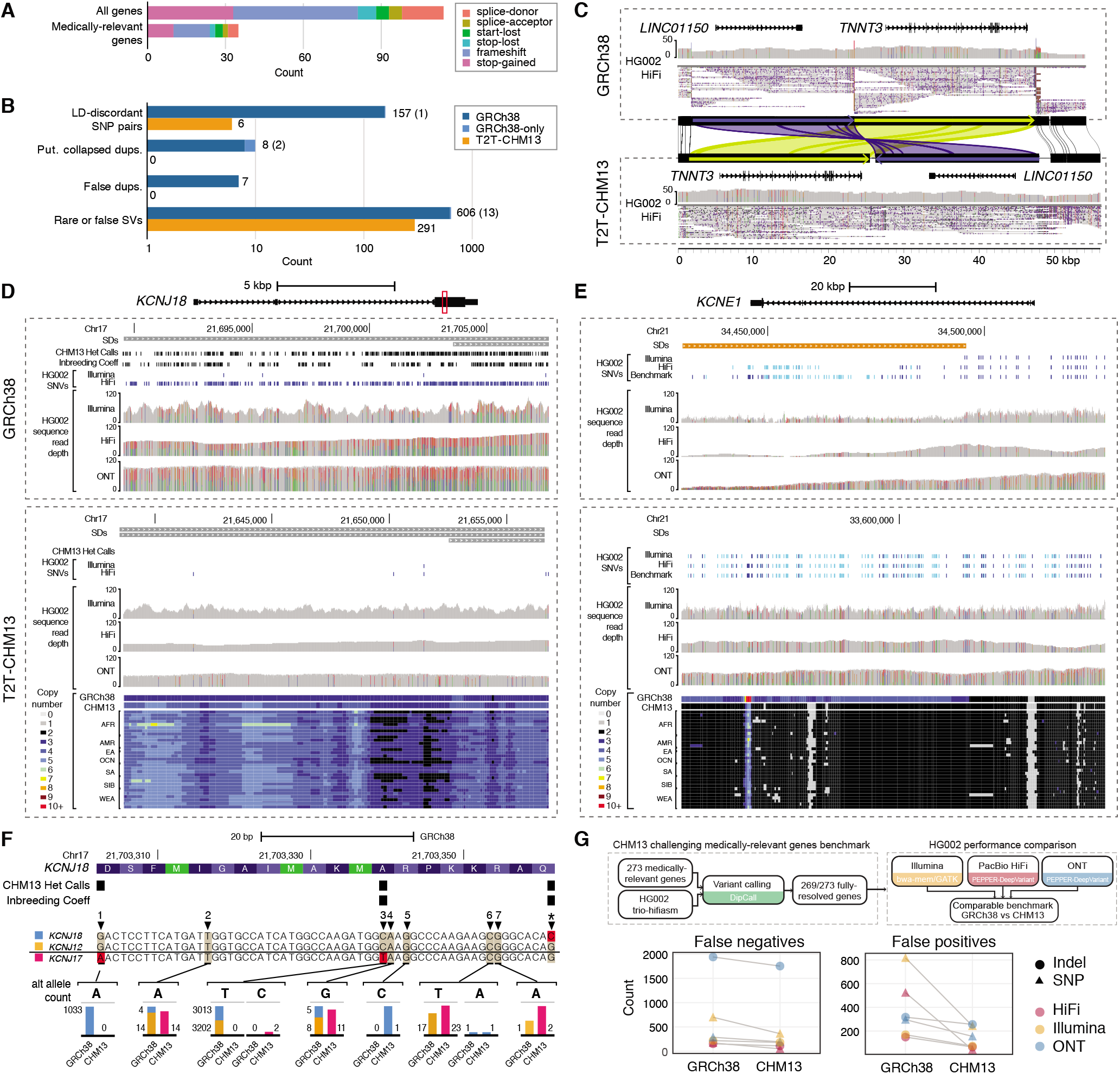
T2T-CHM13 Improves Clinical Genomics Variant Calling. **(A)** Bar plot of potential loss-of-function mutations in the T2T-CHM13 reference. **(B)** The counts of medically-relevant genes impacted by genomic features and variation in GRCh38 (blue) and CHM13 (orange) are depicted as bar plots on logarithmic scale. Light blue indicates genes impacted in GRCh38 where homologous genes were not identified in T2T-CHM13 due to inability to lift over, with counts included in parentheses. **(C)** An example erroneous GRCh38 complex SV corrected in T2T-CHM13 impacting *TNNT3* and *LINC01150*, displayed by sequence comparison using miropeats with homologous regions colored in green and blue, respectively. HG002 PacBio HiFi data is displayed showing read coverages and mappings. Snapshots of regions using Integrative Genomic Viewer and UCSC Genome Browser representing **(D)** a collapsed duplication in GRCh38 corrected in T2T-CHM13 impacting *KCNJ18* and **(E)** a false duplication in GRCh38 impacting most of *KCNE1*. CHM13 heterozygous (het) calls and gnomAD variants flagged for out-ofrange inbreeding coefficient (coeff) are depicted for *KCNJ18*. Read mappings and variants from HG002 Illumina, PacBio HiFi, and ONT (mappings only), with homozygous (light blue) and heterozygous (dark blue) variants depicted as dashes. For *KCNE1*, HG002 benchmark variants are also depicted using the same color scheme. Copy-number estimates, displayed as colors, across k-merized version of the GRCh38 and T2T-CHM13 references as well as representative examples of the SGDP individuals. **(F)** An example CDS region of *KCNJ18* (highlighted as a red box in **D**), with alignments of *KCNJ18* (blue), *KCNJ12* (orange), and *KCNJ17* (pink) along with allele counts of variants in each gene identified on GRCh38 and T2T-CHM13 are shown as bar plots (to approximate scale per variant), with examples 1–7 explained in the Results. **(G)** Schematic depicts a new benchmark for 269 challenging medically relevant genes for HG002. The number of variant-calling errors from three sequencing technologies on each reference is plotted.

Of these 210 variants, 158 have been identified in at least one individual from 1KGP, with most variants relatively common in human populations (median AF of 0.47), suggesting they are functionally tolerated. The remaining variants not found in 1KGP individuals comprise larger indels, which are more difficult to identify using 1KGP Illumina data, as well as alleles that are rare or unique to CHM13. We curated the ten variants impacting medically relevant genes and found seven that likely derived from duplicate paralogs: a 100-bp insertion also found in long reads of HG002, a stop gain in a final exon in one gnomAD sample, and an insertion in a homopolymer in a variable-number tandem repeat in *CEL*, which may be an error in the assembly. Understanding that the published T2T-CHM13 assembly represents a human genome harboring potentially functional or rare variants that in turn would affect the ability to call variants at those sites, we have made available the full list of putative loss-of-function variants to aid in clinical interpretation of sequencing results **(Table S5.1)**.

#### T2T-CHM13 improves variant calling for medically relevant genes

We sought to understand how the transition from GRCh38 to the T2T-CHM13 reference might impact variants identified in a previously compiled [23] set of 4,964 medically relevant genes residing on human autosomes and chromosome X (representing 4,924 genes in T2T-CHM13 via liftover; **Table S5.2**). Of these genes, 28 map to newly accessible regions in T2T-CHM13 (novel and/or non-syntenic). We found over twice as many medically relevant genes impacted by rare or erroneous structural alleles on GRCh38 (n=756 including 14 with no T2T-CHM13 liftover) compared to T2T-CHM13 (n=306) (**Fig. 5B**), of which 622 genes appear corrected in T2T-CHM13. This includes 116 genes falling in regions previously flagged as erroneous in GRCh38 by the GRC. The majority (82%) of impacted clinically relevant genes in GRCh38 overlap SVs that exist in all 13 HiFi-sequenced individuals, likely representing rare alleles or errors in the reference (see above), including 13 of the 14 genes with no T2T-CHM13 liftover.

One example of such a structural error involves *TNNT3*, which encodes Troponin T3, fast skeletal type and is implicated in forms of arthrogryposis [56]. When calling SVs with respect to GRCh38, *TNNT3* is supposedly impacted by a complex structural rearrangement in all individuals, consisting of a 24-kbp inversion and 22-kbp upstream deletion, which also ablates *LINC01150* (**Fig. 5C**). The GRC determined that a problem existed with the GRCh38 reference in this region (GRC issue HG-28). Analysis of this region in T2T-CHM13 instead shows a complex rearrangement with the 22 kbp region upstream of *TNNT3* inversely transposed in the new assembly to the proximal side of the gene. Besides potentially affecting interpretations of gene regulation, this structural correction of the reference places *TNNT3* >20 kbp closer to its genetically-linked partner *TNNI2* [57]. Other genes have VNTRs that are collapsed in GRCh38, such as one expanded by 17 kbp in most individuals in the medically relevant gene *GPI. MUC3A* was also flagged with a whole-gene amplification in all individuals, which we identified as residing within a falsely-collapsed SD in GRCh38, further evidencing that finding (**Fig. 1A**).

Seventeen medically relevant genes reside within erroneous duplicated and putative collapsed regions in GRCh38 **(Tables S1.1 and S1.3**), including *KCNE1* (false duplication) and *KCNJ18* (collapsed duplication) (**Figs. 5D and E**). For these genes, we show that a significant skew in total variant density occurs in GRCh38 (58 variants per kbp for eight genes in collapsed duplications and 21 variants per kbp for seven genes in false duplications;*p-values* = 5.684 x 10^-3^ and 6.195 x 10^-4^, respectively, Mann-Whitney U test) versus the rest of the 4,909 medically-relevant gene set (40 variants per kbp) that largely disappears in T2T-CHM13 (40 variants per kbp in collapsed duplications and 47 variants per kbp in false duplications versus 41 variants per kbp for the remaining gene set; *p-values* = 0.8778 and 0.0219, respectively) (**Fig. S4.2**). Examining the falsely-duplicated *KCNE1*, we find that coverage is much lower than normal on GRCh38 for short and long reads and that most variants are missed because many reads incorrectly map to the false duplication *(KCNE1B* on the p-arm of chromosome 21). The kmer-based copy number of this region in all 266 SGDP genomes supports the T2T-CHM13 copy number, and this region was not duplicated in GRCh37 [23]. As for *KCNJ18*, which resides within a GRCh38 collapsed duplication at chromosome 17p11.2 [58], we find increased coverage and variants within HG002 using short- and long-read sequences in GRCh38 relative to T2T-CHM13.

To verify if the additional variants identified using GRCh38 are false heterozygous calls from PSVs derived from missing duplicate paralogs, we compared the distributions of minor-allele frequencies across the 49-kbp SD and observed a shift in SNV proportions, with a relative decrease in intermediate-frequency alleles and a relative increase in rare alleles for *KCNJ18* and *KCNJ12* (another collapsed duplication residing distally at chromosome 17p11.2) in T2T-CHM13 compared with GRCh38 (*p*-value = 8.885 × 10^-2^ and 3.102 x 10^-2^, respectively; Mann-Whitney U test) (**Fig. S5.1**). We matched the homologous positions of discovered alternative alleles in GRCh38 and T2T-CHM13 across the three paralogs—including the previously missing paralog located in a centromere-associated region on chromosome 17p *KCNJ17* denoted *KCNJ18-1* in T2T-CHM13—and observed that even true variants (i.e., non-PSVs) had discordant allele counts in *KCNJ18* and *KCNJ12* between the two references. Examples of such variants impacting the coding sequence (CDS) of these genes are highlighted in **Fig. 5F** and labeled as 1–7, representing different categories:

- (1) false positives in *KCNJ12* and/or *KCNJ18* on GRCh38 resulting from PSVs in *KCNJ17*, which are corrected in T2T-CHM13;
- (2) and (4) variants erroneously detected in *KCNJ12* and *KCNJ18* on GRCh38 that actually are variants in *KCNJ17*;
- (3) false positives in *KCNJ12* and/or *KCNJ18* on GRCh38 resulting from PSVs in *KCNJ17* that also hide true variants in *KCNJ17, KCNJ12*, and/or *KCNJ18;*
- (5) variants not detected in *KCNJ12* or *KCNJ18* on GRCh38, likely due to skewed allele balance from mis-mapped reads;
- (6) and (7) variants erroneously detected in *KCNJ12* and/or *KCNJ18* on GRCh38 that actually are variants in *KCNJ17* with the allele detected in fewer haplotypes on GRCh38, likely due to skewed allele balance from mis-mapped reads.

We also identified more complex sites, such as the last base in **Fig. 5F** labeled with an asterisk, in which the reference base for *KCNJ18* changes to C in T2T-CHM13, in addition to novel alleles identified in *KCNJ17* and *KCNJ18*. Considering rare variants of *KCNJ18* contribute to muscle channelopathy-thyrotoxic periodic paralysis [58] including nine “pathogenic” or “likely pathogenic” variants in ClinVar, increased sensitivity to discover variants in patients using T2T-CHM13 would have a significant clinical impact. In summary, T2T-CHM13’s improved representation of this gene and other collapsed duplications not only eliminates false positives but also improves detection and genotyping of true variants.

#### Clinical gene benchmark demonstrates T2T-CHM13 reduces errors across technologies

Finally, to determine how the T2T-CHM13 genome improved our ability to assay variation broadly, we used a curated diploid assembly to develop a new benchmark for 269 challenging medically-relevant genes in GIAB Ashkenazi son HG002 [23], with comparable benchmark regions on GRCh38 and T2T-CHM13. We tested three short- and long-read variant callsets against this benchmark: Illumina-BWAMEM-GATK, HiFi-PEPPER-DeepVariant, and ONT-PEPPER-DeepVariant. Both the numbers of false positives and false negatives substantially decrease for all three callsets when using T2T-CHM13 as a reference instead of GRCh38 (**Fig. 5G** and **Table S5.3**). The number of false positives for HiFi decreases by a factor of 12 in these genes, primarily due to the addition of missing sequences similar to *KMT2C* (**Fig. S1.12**) and removal of false duplications of *CBS, CRYAA, H19*, and *KCNE1* (**Fig. 5G**). As demonstrated above, these genes and others are better represented in T2T-CHM13 than GRCh38 for a diverse set of individuals, so performance should be higher across diverse ancestries. The number of true positives also decreases by a much smaller fraction than the errors (~14%), due to a reduction of true homozygous variants caused by T2T-CHM13 possessing fewer ultra-rare and private alleles (**Fig. 2G**). This benchmarking demonstrates concrete performance gains in specific medically relevant genes resulting from the highly accurate assembly of a single individual.

## Discussion

Persistent errors in the human reference genome, ranging from collapsed duplications to missing sequences, have silently plagued human genetics for decades. The strong and arguably misguided assumption that most genomic analyses make about the correctness of the reference genome has led to spurious clinical findings and mistaken disease associations. For example, a few previous clinical exome studies have erroneously implicated PSVs of missing paralogs of GRCh38 collapsed duplications, such as *KCNJ18* and *GPRIN2*, as contributing to diseases [59–62]. Here, we resolve variation in these regions and show that the T2T-CHM13 reference genome universally improves genomic analyses for all populations by correcting major structural defects and adding sequences that were absent from GRCh38. In particular, we show that the T2T-CHM13 assembly (1) revealed millions of additional variants and the existence of new copies of medically relevant genes (e.g., *KCNJ17)* within the 240 Mbp and 189 Mbp of non-syntenic and novel sequence, respectively; (2) eliminated tens of thousands of spurious variants and incorrect genotypes per samples, including within medically relevant genes (e.g., *KCNJ18*) by expanding 203 loci (8.04 Mbp) that were falsely collapsed in GRCh38; (3) improved genotyping by eliminating 12 loci (1.2 Mbp) that were falsely duplicated in GRCh38; and (4) yielded more comprehensive SV calling genome-wide, with a greatly improved insertion/deletion balance, by correcting collapsed tandem repeats. Overall, the T2T-CHM13 assembly reduced false positive and false negative SNVs from short and long reads by as much as 12-fold in challenging, medically relevant genes. The T2T-CHM13 reference also accurately represents the haplotype structure of human genomes, eliminating 1,390 artificial recombinant haplotypes in GRCh38 that occurred as artifacts of BAC clone boundaries. These improvements will broadly enable new discoveries and refine analyses across all of human genetics and genomics.

Given these advances, we advocate for a rapid transition to the T2T-CHM13 genome. On a practical level, improvements to large genome segments, such as entire p-arms of the acrocentric chromosomes, and the discovery of new clinically relevant genes and disease-causing variants justify the labor and cost required to incorporate T2T-CHM13 into basic science and clinical genomic studies. On a technical level, T2T-CHM13 simplifies genome analysis and interpretation because it consists of 23 complete linear sequences and is free of “patch”, unplaced, or unlocalized sequences. In contrast, GRCh38 has 195 different contigs (with names such as chr14_GL000009v2_random and chrUn_GL000195v1) that are often excluded, as interpreting results in these regions is challenging. Many of the fixes made in T2T-CHM13 were previously curated by the Genome Reference Consortium and have fix patches, but few studies use these existing fix patches. The reduced contig set of T2T-CHM13 also facilitates interpretation and is directly compatible with essentially all of the most commonly used analysis tools (e.g., BWA-MEM, GATK, bedtools, samtools, bcftools, etc). To promote this transition, we provide variant calls and several other annotations for the T2T-CHM13 genome within the UCSC Genome Browser and the NHGRI AnVIL as a resource for the human genomics and medical communities.

Finally, our work underscores the need for additional T2T genomes. Most urgently, the CHM13 genome lacks a Y chromosome, so our analysis relied on the incomplete representation of chrY from GRCh38. We expect a T2T representation of chrY within the very near future, at which time we will study how including this sequence improves mapping and variant analysis on the Y chromosome. Furthermore, many of the new regions in T2T-CHM13 are present in all human genomes and enable variant calling with traditional methods from short and/or long reads. However, many novel regions exhibit substantial variation within and between populations, including satellite DNA [31] and SDs that are polymorphic in copy number and structure [32]. Relatedly, the T2T-CHM13 reference provides a basis for calling millions of new variants, but many of these variants are challenging to resolve accurately using current sequencing technologies and analysis algorithms. Robust variant calling in these regions will require many diverse haplotype-resolved T2T assemblies to construct a pangenome reference, in turn motivating the development of new methods for discovering, representing, comparing, and interpreting complex variation, as well as benchmarks to evaluate their performance [63,64].

Through our detailed assessment of variant calling at known and novel loci across global population samples, our study showcases T2T-CHM13 as the preeminent reference for human genetics. The annotation resources provided herein will help facilitate this transition, expanding knowledge of human genetic diversity by revealing functional variation that was previously hidden from view.

## Acknowledgements

We would like to thank Michael Zody, Bjöm Grüning, Heng Li, Sasha Langley, Chuck Langley, Geraldine Van der Auwera, Valerie Schneider, Steven Salzberg, Ben Langmead, Alexis Battle and several of their lab members for helpful discussions. Certain commercial equipment, instruments, or materials are identified to specify adequately experimental conditions or reported results. Such identification does not imply recommendation or endorsement by the National Institute of Standards and Technology, nor does it imply that the equipment, instruments, or materials identified are necessarily the best available for the purpose. This work utilized the computational resources of the NIH HPC Biowulf cluster (https://hpc.nih.gov) and the Maryland Advanced Research Computing Center (https://www.marcc.ihu.edu/).

## Author contributions

Annotation and comparison of GRCh38 and T2T-CHM13: AS, AR, DCS, DEM, DJT, SMY, MEGS, MRV, NFH, MYD, JMZ, RCM; Short-read analysis: SZ, SA, SMY, RCM, MCS; Long-read analysis: KS, MK, SA, JZ, MCS; Population genetics: SA, DJT, SMY, MCS, RCM; Clinical variant analysis: DCS, DEM, MYD, JMZ; Variant analysis: AMP, BP, CSC, CX, DCS, FJS, JL, JM, JMZ, JW, MM, MCS, MYD, NDO, PA, RCM, RL, SA, SK, SM; Project design: AMP, BP, FJS, JAR, JMZ, KHM, MYD, RCM, MCS, RL; Paper writing: RCM, MYD, JMZ, and MCS with input from all of the authors

## Data and materials availability

UCSC assembly hub browser: http://genome.ucsc.edu/cgi-bin/hgTracks?genome=t2t-chm13-v1.0&hubUrl=http://t2t.gi.ucsc.edu/chm13/hub/hub.txt

Aligned reads, variant calls, and other summarizations are available within the NHGRI AnVIL Platform, along with Jupyter notebooks for computing the analysis: https://anvil.terra.bio/#workspaces/anvil-datastorage/AnVIL_T2T

Variant calling workflows and downstream analysis scripts: https://github.com/schatzlab/t2t-variants https://github.com/mccoy-lab/t2t-variants

Genome in a Bottle Challenging Medically-Relevant Gene Benchmarks: https://ftp-trace.ncbi.nlm.nih.gov/ReferenceSamples/giab/release/AshkenazimTrio/HG002_NA24385_son/CMRG_v1.00/

