## Supplemental Text and Figures for "A complete reference genome improves analysis of human genetic variation"

#### Table of Contents

**Section I. Structural Comparisons of GRCh38 and T2T-CHM13** **3**
Mappability Analysis 3
Generating variant calls with assembly-assembly alignment of T2T-CHM13 to GRCh38 3
Inference of local ancestry and Neanderthal introgression for GRCh38 and T2T-CHM13 3
Identification of non-syntenic and novel regions 5
Quantifying artificial haplotype structure of GRCh38 5
Identification of collapsed duplications in GRCh38 5
Identifying false duplications in GRCh38 6
Genome Reference Consortium investigations into false duplications and contaminations in GRCh38 7
Liftover failures 8
Additional reference annotations and impacted genes 8

**Section II. T2T-CHM13 improves analysis of global genetic diversity based on 3,202 short-read samples from the 1KGP dataset** **9**
Short-read alignment, coverage analysis, and variant calling of the expanded 1KGP collection 9
Comparison of GRCh38/NY Genome Center Pipeline to T2T-CHM13/AnVIL Pipeline 11
Visibility and concordance of SNVs when using different reference genomes for variant discovery 11

**Section III. T2T-CHM13 improves structural variant analysis of 17 diverse long-read samples** **13**
Long-read alignment and coverage analysis 13
Long-read variant calling analysis 14
Optical mapping assembly and variant calling 14

**Section IV. Characterization of non-syntenic regions** **16**
Newly added region analysis 16
Identifying concordance between PacBio HiFi, ONT and Illumina variant calls 16

Distance and LD between non-syntenic regions and phenotype-associated SNVs 16
Identifying frequency-differentiated variants in non-syntenic T2T-CHM13 regions 17
Comparison of frequency-differentiated variants between T2T-CHM13 and GRCh38 17
Collapsed duplications and false duplication variant comparisons 18

**Section V. Impact of T2T-CHM13 on Clinical Genomics 19**
Annotation of T2T-CHM13 variants using Variant Effect Predictor (VEP) 19
Identification of medically relevant genes with impacted variant discovery 19
Creating a difficult medically relevant gene benchmark for HG002 on T2T-CHM13 19

**Supplementary Figures 21**

#### Section I. Structural Comparisons of GRCh38 and T2T-CHM13

##### Mappability Analysis

We calculated minimum unique k-mer (MUK) lengths for both T2T-CHM13 and GRCh38. MUKs represent the minimum distance from a position in the genome needed to identify a unique sequence, either upstream (right-anchored) or downstream (left-anchored) (**Fig. S1.2 and S1.3**). To compute these values, all chromosomes of the target genome were concatenated, followed by their reverse complements. A suffix array (SA) and longest common prefix (LCP) table were calculated from this single sequence using the algorithm and implementation adapted from Sapling [65]. For each position in the SA, the LCP value plus one represents the MUK at that position. If the SA value indicated that the sequence was a reverse complement, the unique sequence was right-anchored; otherwise, the sequence was left-anchored. Any minimum unique sequences containing an N or overlapping a chromosome end were removed and those positions were marked as having no minimum unique sequence. All scripts for recapitulating this analysis can be found at https://github.com/msauria/T2T_MUK_Analysis.

Examining the tails of the MUK distribution, e.g. the longest perfect repeats in these genomes, we found that GRCh38 contained 70 repeated sequences larger than 5 kb while T2T-CHM13 had 81 such sequence pairs (**Fig. S1.4 and S1.5**). Within GRCh38, more than half of these pairs (42) occurred between an unplaced contig and centromeric (or immediately adjacent), subtelomeric, or another unplaced contig's sequence, accounting for 29, 7, and 6 pairs, respectively. Another 18 sequence pairs in GRCh38 occurred between subtelomeric regions. Interestingly, 6 sequence pairs represented subtelomeric-centromeric sequences. There were 2 instances of sequences shared between centromeric regions, another between a subtelomeric region and the middle of chromosome 6, and a 12 kb sequence occurring on both chromosomes X and 8 in an intron of the gene *RBPMS*.

All but one sequence pair of long perfect repeats were removed in the T2T-CHM13 genome assembly as we were able to capture minute differences between these sequences using the long-read technologies. Of the subtelomeric/centromeric mixed pairs, 3 were resolved to the subtelomeric side, 1 to the centromeric side, and 2 had no corresponding mapping in T2T-CHM13. The centromeric/other pair resolved to chromosome 6. Finally, the chromosome X/6 sequence pair resolved to chromosome 6. T2T-CHM13 contains 81 long sequence pairs, 77 of which occur between acrocentric centromere pairs, Of the remaining 4, 3 overlap a single subtelomeric pairing from GRCh38 and the last pair occurs in the middle of chromosomes 2 and 22, exactly encompassing an L1HS LINE element.

##### Generating variant calls with assembly-assembly alignment of T2T-CHM13 to GRCh38

We generated variant calls between T2T-CHM13 and GRCh38 with dipcall (https://github.com/lh3/dipcall) [53] using the T2T-CHM13v1.0 assembly FASTA as haplotype 1 and haplotype 2 with GRCh38 as a reference, as in the NIST assembly benchmarking pipeline (https://github.com/usnistgov/giab-asm-benchmarking). Dipcall generates variant calls using any non-reference support in regions ≥50 kb with contigs having mapping quality ≥5. Dipcall also produces a BED which denotes confident regions covered by an alignment ≥50 kb with contigs having mapping quality ≥5 and with no other alignments ≥10 kb in length.

##### Inference of local ancestry and Neanderthal introgression for GRCh38 and T2T-CHM13

The full description of how the local ancestries and Neanderthal introgressions were identified is reported by Nurk *et al. [27]*. Briefly, we used RFMix v2.03-r0 [66] (https://github.com/slowkoni/rfmix) to infer the local ancestry of the T2T-CHM13 genome. As a set of reference samples for ancestry, we used the 2,504 individuals sequenced in the Phase 3 release of the 1KGP (http://ftp.1000genomes.ebi.ac.uk/vol1/ftp/data_collections/1000_genomes_project/release/20181203_biallelic_SNV/)

[18]. We assumed that T2T-CHM13 carried the reference allele at sites in the 1KGP dataset that were not called as T2T-CHM13 variants by dipcall (described above). All CHM13 genotypes were represented as phased homozygous diploid genotypes (either reference/reference or alt/alt), which resulted in two sets of identical inferred local ancestry segments for each haplotype. We obtained a genetic map for GRCh38, originally generated on build 35 of the human reference genome by the HapMap Consortium [67], from the Beagle resource bundle (http://bochet.gcc.biostat.washington.edu/beagle/genetic_maps/).

We ran RFMix with default parameters (conditional random field spacing = 50 SNPs; random forest window size = 5 SNPs), grouping the 1KGP reference panel into superpopulations (African, Ad Mixed American, East Asian, European, Southeast Asian). Because individuals with 46,XY karyotypes are haploid for variants on the X chromosome, we included only individuals with 46,XX karyotypes in the reference panel for local ancestry inference for the X chromosome. We excluded from the results regions overlapping with centromeres (UCSC Genome Browser; https://genome.ucsc.edu/cgi-bin/hgTables) or that were marked as inaccessible for the 1KGP (http://ftp.1000genomes.ebi.ac.uk/vol1/ftp/data_collections/1000_genomes_project/working/20160622_genome_mask_GRCh38/). The non-pseudoautosomal regions (PARs) of the X chromosome were also inaccessible for this analysis due to the absence of variant calls outside the PARs.

Inspired by related approaches [24,68], we used an alternative strategy for local ancestry inference of GRCh38. By restricting analysis to individual clones, our approach avoids strong assumptions about the genetic map, which could produce misleading results given the artificial haplotype that we previously described. Specifically, for each clone reported in the tiling path, we extracted phased haplotypes as binary vectors of the 1KGP samples from the corresponding genomic interval. We then computed the pairwise Hamming distances between each of these reference haplotypes and the reference GRCh38 haplotype (represented as an equal-length vector of zeros). We identified the nearest-neighbor reference haplotypes to GRCh38 as those with the smallest Hamming distance, recording the superpopulation membership of each corresponding reference individual. Ancestry of the clone was then assigned as the majority superpopulation of the nearest-neighbor haplotypes, normalizing by the relative abundances of superpopulations within the entire dataset. Clones for which no superpopulation achieved the majority among the nearest neighbors were designated as ambiguous, as were clones without SNPs in the 1KGP sample (i.e., occurring within inaccessible regions of the genome).

To identify Neanderthal-introgressed haplotypes, we used IBDmix (https://github.com/PrincetonUniversity/IBDmix), which detects introgressed archaic sequences in a set of individuals by identifying haplotypes that are identical-by-descent to a reference archaic sample [69]. Of the three Neanderthal individuals with available high-coverage sequences, we chose the Vindija Neanderthal as our reference archaic individual due to its inferred closest relation to the Neanderthal population that admixed with modern humans [70]. Because IBDmix identifies introgressed sequences in a set of modern samples, assumed to be from the same population, we additionally provided samples from the 1KGP as a background set of modern individuals. These samples were sequenced to 30x coverage by the New York Genome Center, and variants were called on the GRCh38 reference (http://ftp.1000genomes.ebi.ac.uk/vol1/ftp/data_collections/1000G_2504_high_coverage/working/20201028_3202_phased/) [28].

Because local ancestry analysis indicated that the majority of the CHM13 genome is composed of European ancestries, we used European samples as the background set of modern individuals for CHM13. For GRCh38, which comprises haplotypes from multiple populations, we identified introgressed segments by pairing it with each of the 1KGP superpopulations. In a given local ancestry tract, we only retained introgression calls from the background superpopulation matching the local ancestry in that region.

We required all Neanderthal-introgressed haplotypes to have a LOD score of at least 4 and a length of at least 50 kbp. As with the local ancestry segments, we removed introgressed segments within regions deemed inaccessible by the 1KGP.

Because we were only able to call introgression in the 72.9% GRCh38 clones with confident ancestry assignments, we also sought to verify that GRCh38's lower amount of introgressed sequence (26.7 Mbp, in comparison with CHM13's 51.0 Mbp) was not an artifact of restricting our analysis to a smaller set of regions. Using the UCSC Genome Browser's liftOver tool [71], we lifted over the boundaries of confidently assigned GRCh38 clones to T2T-CHM13, discarding the clones that failed to lift over. Within these lifted over regions, T2T-CHM13 carries 43.6 Mbp of Neanderthal-introgressed sequence, in line with our expectations due to GRCh38's higher proportion of African ancestry.

#### Identification of non-syntenic and novel regions

A complete description of how the non-syntenic and novel regions of T2T-CHM13 were identified is described by Nurk *et al.* [27]. Briefly, non-syntenic sequence in T2T-CHM13 was based on whole-genome LastZ [72] alignment of GRCh38 and T2T-CHM13, filtering for any regions without 1 Mbp of synteny. A more conservative set of novel regions was identified as those regions not covered by alignment of T2T-CHM13 to GRCh38 using Winnowmap v2.0.1 [44].

#### Quantifying artificial haplotype structure of GRCh38

We used PLINK v1.90b6.21 [73] to extract all autosomal SNP pairs with perfect LD ($R^2 = 1$) in the entire 1 sample set using recently published genotype data from the 30x sequencing by NYGC. By definition, GRCh38 carries the reference allele at all sites, so we computed the frequency of REF-REF haplotypes at each of these SNP pairs among the 1KGP samples. We then used HTSlib v.1.12 [74] to query T2T-CHM13 alleles at each corresponding site, again computing the frequency of the observed T2T-CHM13 haplotypes among the 1KGP sample set. This analysis of T2T-CHM13 alleles on GRCh38 coordinates was based on the aforementioned variant calls produced by dipcall, restricting to portions of the alignment involving only a single contig (i.e., the region defined by the dip.bed output file) and excluding regions within 10 bp of an indel that distinguishes the two assemblies. All queried genomes were subjected to these same filtering criteria to ensure a common baseline for comparison. For the GRCh38 haplotypes that were absent (i.e. frequency = 0) among the 1KGP sample set, we intersected the intervals defined by the SNP pairs with the clone tiling path used for the construction of the GRCh38 reference genome (https://hgdownload-test.gi.ucsc.edu/goldenPath/hg38/bigZips/hg38.agp.gz), thereby quantifying the number of rare allelic combinations explained by transitions between adjacent clones.

To compare these results to samples from the 1KGP, we performed a leave-one-out analysis in which for each iteration, we randomly selected a single phased "haploid" genome from the 1KGP sample set and re-computed LD for the full dataset minus that individual. For all SNP pairs in perfect LD ($R^2 = 1$) in the new sample (N = 2503 individuals), we queried the allelic state of the selected haploid sample in the manner previously described. We repeated this analysis for 100 randomly sampled individuals, generating superpopulation-specific distributions of LD-discordant SNP pairs per haploid genome. We again applied the same filtering criteria as were applied to T2T-CHM13 and GRCh38 to ensure a fair comparison.

#### Identification of collapsed duplications in GRCh38

We simulated Illumina-like reads (400 million PE 150 bp reads) from T2T-CHM13 reference v1.0 including the GRCh38 Y chromosome using Mason (https://github.com/seqan/seqan/tree/master/apps/mason2) and aligned them to GRCh38 (no alt or decoy contigs) and T2T-CHM13 v1.0 including the GRCh38 Y chromosome using BWA-MEM [40] (**Fig. S1.10**). Likewise, previously published CHM13 PacBio HiFi reads (~24X, SRA: SRX5633451) [75] were aligned to GRCh38 using minimap2 [45] with the -ax map-pb setting. We called SNVs in both datasets with GATK v4.1.8.1 [41] using minimum MAPQ 30, ploidy 2 and otherwise default parameters. Only PASS variants were used for downstream analyses. Heterozygous variants called by each platform were merged into one multi-sample VCF file with bcftools merge, and the number of heterozygous variants per kbp was calculated using bedtools coverage. For both references, we first defined problematic regions as regions ≥2 kbp with ≥2 heterozygous calls in the CHM13 sample. From this, we connected regions separated by ≥5 kbp, and then filtered for regions ≥5 kbp in size.

Focusing on GRCh38-derived problematic regions, we intersected them with previously published RepeatMasker and SD annotations obtained from UCSC Table Browser, as well as known GRC issues [27](ftp://ftp.ncbi.nlm.nih.gov/pub/grc/human/GRC/Issue_Mapping/). For each region, we determined association with SDs [32], centromeres [31], and non-syntenic and novel regions in T2T-CHM13 reference [27], using combined lifted coordinates obtained from all minimap2 hits and UCSC LiftOver [71].

Additionally, we obtained variants flagged with excess of heterozygosity by the gnomAD database (InbreedingCoeff in the FILTER field), defined as variants with an inbreeding coefficient < -0.3, but were not filtered due to low read depth, genotype quality, or minor-allele fraction [6]. Empirical enrichment of variants with excessive heterozygosity within problematic regions was obtained by calculating the number of variants in 10,000 randomly sampled regions of the genome using bedtools shuffle. The empirical *p*-value was calculated as (M+1)/(N+1), where M is the number of iterations yielding a number of features greater than observed and N is the number of iterations.

We identified each homologous GRCh38 problematic region in T2T-CHM13 using the following approach: (1) coordinates obtained by UCSC LiftOver using available chain [27] if the size of the lifted region was within 80-120% of the original size, (2) Minimap2 longest hit if the size of the lifted region was within 80-120% of the original size, and closest hit when more than two options were available, and (3) manual selection and curation of remaining coordinates. To assess functional impact, these likely problematic regions were intersected with all gene features in Gencode v36, as well as a curated list of medically relevant genes [23].

Using available read-depth copy number estimates in T2T-CHM13 [32], we obtained the overall copy number of the lifted regions as the median window copy number for a "*k*-merized" version of GRCh38 and T2T-CHM13 references, as well as 268 individuals from the SGDP dataset (excluding sample LP6005442-DNA_A08). Regions where copy number in GRCh38 was lower than T2T-CHM13 and also nearly all SGDP individuals (allowing for one individual with lower copy number) were considered putative collapsed duplications in the GRCh38 reference. Additionally, we intersected lifted coordinates with T2T-CHM13 SDs [32] and centromere annotations [31].

The same analysis was performed to identify putative collapsed duplications in T2T-CHM13, but without the need to liftover homologous coordinates.

#### Identifying false duplications in GRCh38

To identify falsely duplicated regions in GRCh38, we compared copy number estimates of GRCh38 to copy number estimates of 268 genomes from the SGDP dataset using short reads, using a method analogous to comparative read-depth approaches described previously [35,76]. We first averaged the copy number estimates for each genome across 1 kbp windows. For each 1 kbp region, we flagged it as a potential false duplication if the copy number in GRCh38 was greater than the copy number in 99% of the other genomes. Flagged regions were assigned a value of 1 and unflagged regions were assigned a value of 0. To filter the flagged regions, we used a median filter approach with a window size of 3 kbp, where the binary value of each 1 kbp region was replaced with the median value of the complete window. We then merged all adjacent flagged regions and reported the start and end coordinates with respect to T2T-CHM13. To find the corresponding locations of the duplications on GRCh38, we used minimap2 [45] version 2.17-r941 with parameter -p 0.25. Some regions mapped to more than two locations on GRCh38 due to true SDs in the genome. We curated these regions with more than two alignments, and identified the incorrect region(s) as the region(s) that did not have an assembly-assembly alignment from T2T-CHM13 or the HG002 haplotype. We identified the affected, correct region as the region that aligned most closely to the T2T-CHM13 region, which also had reduced HG002 read coverage. Upon curation of the regions with only two alignments on GRCh38, we selected as correct the region that was on the same chromosome arm as the corresponding T2T-CHM13 region. When both regions were on the same chromosome arm, we selected as correct the region that was not adjacent to or between gaps in GRCh38. One false duplication was a tandem duplication, and we arbitrarily selected one copy as correct. Upon curation, we also removed one small 8 kb region (chr19:14,359,000-14,367,000 on T2T-CHM13) that was incorrectly identified as falsely duplicated.

#### Genome Reference Consortium investigations into false duplications and contaminations in GRCh38

Since the release of GRCh38, the GRC has received a number of user reports alerting us to a potential false duplication involving chr 21p and 21q. Users noted that reads were aligning to both regions in GRCh38, but not GRCh37/hg19, resulting in a decreased mapping score and difficulties in variant calling throughout. Additionally, user analyses involving Multiplex Ligation-dependent Probe Amplification (MLPA), a technique for gene copy number detection, and exome studies indicated potential false duplications. The implicated regions contained several genes, including *CBSL* (Gene ID: 102724560), *U2AF1L5* (Gene ID: 102724594) and *KCNE1B* (Gene ID: 102723475). The GRC has investigated the matter and concurs that the GRCh38 assembly contains sequence on the short arm of chr 21 that should be excluded from analyses.

The short arm of human chromosome 21, like that of the four other human acrocentric chromosomes, is where genes associated with rDNA synthesis are localized, and is characterized by highly repetitive heterochromatic sequence. The repetitive nature of these sequences, coupled with limitations in sequencing technology, have until recently made the representation of these regions in genome assemblies very difficult. As a consequence, the GRCh37 representation of the chromosome 21 p-arm contained only 11 clone sequences. Seven were clones from the HSA21-specific BAC library CHORI-507 that had previously been experimentally localized to 21p [77]. In an effort to add additional sequence to this repetitive region, 23 additional components were added to 21p for GRCh38, including 18 additional CHORI-507 clones, 4 RPCI-11 clones, and 1 ABC9 fosmid. Admixture mapping localized some of these clones to this region.

In response to the user reports, the GRC re-reviewed the sequences added to 21p in GRCh38. Haploid CHM13hTERT Illumina reads generated by The McDonnell Genome Institute were aligned to GRCh38 by NCBI, and evaluated for read mapping and coverage. This analysis supported the user reports, suggesting that 5 of the newly added CHORI-507 clones (FP565260.4, CU639417.17, FP236240.8, FP475955.4 and CU633980.13) were actually redundant with sequences on chr 21q, and thus represented false duplications in GRCh38.

The GRC has now removed these sequences from the files that it uses to generate the reference assembly. However, they cannot remove them from the GRCh38 assembly without triggering the next major release of the human assembly as the coordinates will shift for all downstream sequences. In order to help users recognize these regions and avoid them in their analyses, the GRC produced a v1 masking file to be used as a companion to GRCh38 at https://ftp.ncbi.nlm.nih.gov/genomes/all/GCA/000/001/405/GCA_000001405.15_GRCh38/seqs_for_alignment_pipelines.ucsc_ids/GCA_000001405.15_GRCh38_GRC_exclusions.bed. This file provides the assembly coordinates of the 5 clones incorrectly localized to chr 21p. The Genome in a Bottle Consortium produced a fasta version of GRCh38 with these regions masked, as well as v2 masked GRCh38 fasta with additional false duplications identified in this manuscript under https://ftp-trace.ncbi.nlm.nih.gov/ReferenceSamples/giab/release/references/. The v2 masking file is available at https://ftp-trace.ncbi.nlm.nih.gov/ReferenceSamples/giab/release/references/GRCh38/GCA_000001405.15_GRCh38_GRC_exclusions_T2Tv2.bed.

In addition to these sequences, the file also includes 2 other assembly scaffolds that were found, after the release of GRCh38, to be contaminated with non-human sequence. These include a chrUn scaffold (KI270752.1/NT_187507.1), whose sole component (AF065393.1) is now known to represent sequence from Chinese hamster [78], likely derived from the human-hamster CHO cell line that was the clone source, and an alternate loci scaffold (KI270825.1/NT_187580.1) whose non-anchor component (AC225822.3) was shown to be chimeric. In AC225822.3, the first 25,375 bases are human sequence matching GRCh38 chr10 reference component and alternate scaffold anchor sequence AL391421.27, while the rest match *Acidithiobacillus thiooxidans* sequences from multiple WGS projects [79]. Although all sequences in the reference assembly are screened for foreign contamination, these two were not detected at the time of release (2014). Prompted by these findings, the GRC has more recently re-screened the assembly with updated contamination databases and has not detected additional issues. As these two scaffolds are not human sequence, very few reads are likely to map well to them, but users may still want to make note of them in their analyses. In total, the contamination represents ~800 Kb, or 0.02% of the total sequence length and the GRC remains committed to addressing assembly errors and making sure it serves as the most reliable analysis substrate possible.

#### Liftover failures

We lifted over dbSNP build 154 (ftp.ncbi.nih.gov/snp/archive/b154/) [80], the March 8th, 2021 release of Clinvar (ftp.ncbi.nlm.nih.gov/pub/clinvar/vcf_GRCh38/weekly/) [81], and GWAS Catalog v1.0 (ebi.ac.uk/gwas/home) [50] from the GRCh38 assembly to the T2T-CHM13 assembly. We did not attempt to lift over any variants on primary or non-primary Y chromosome contigs as this sequence has not changed. To generate a VCF file for portion of the GWAS Catalog that could be lifted over, we first identified all RefSeq IDs (rs ids) in the v1.0 associations file (ebi.ac.uk/gwas/api/search/downloads/full), in the SNPS, SNP_ID_CURRENT, and SNP-RISK ALLELE fields. This yielded 154,054 unique rs ids, which we then intersected with all dbSNP variants on the primary contigs for Chromosomes 1-22, Chromosome X and the mitochondrial chromosome, subsetting to those rsids that matched only a single record in dbSNP. This resulted in 151,876 unique variants from the GWAS Catalog that were used for liftover. We identified “pathogenic” Clinvar variants as any Clinvar variants that were annotated as “pathogenic”, “likely_pathogenic”, or “pathogenic/likely_pathogenic”, had conflicting annotations including at least one of these three annotations, or were part of a haplotype annotated with one of these three annotations (identified through the CLINSIG, CLNSIGCONF, and CLNSIGINCL info fields respectively).

We performed liftover using the GATK release 3.1.9 [82] LiftoverVcf (Picard) tool using the available GRCh38 to T2T-CHM13 chain file [27] with the default parameters. This successfully lifts over variants that map exactly from GRCh38 to T2T-CHM13 but does not recover variants with swapped reference and alternative alleles. To recover variants with swapped ref/alt alleles, we ran LiftoverVCF again, with the “RECOVER_SWAPPED_REF_ALT” flag. Notably, this feature does not recover multiallelic variants, so to recover these variants, we first separated them into multiple biallelic variants, performed liftover using the “RECOVER_SWAPPED_REF_ALT” tag, and converted them back to their multiallelic representations. Variants whose position lifted over, but whose reference allele or any alternative alleles did not match the T2T-CHM13 allele were not recovered.

To determine how many variants failed to lift over because they overlap an indel that distinguishes T2T-CHM13 and GRCh38, we intersected those variants whose position failed to lift over (i.e., those failures not annotated as a ref/alt swap or ref mismatch) with the T2T-CHM13-on-GRCh38 alignment generated using dipcall (described above).

#### Additional reference annotations and impacted genes

We annotated GRCh38 and T2T-CHM13v1.0 references using previously published genomic features (**Fig S1.1**). For GRCh38, we included known GRC issues [27] (ftp://ftp.ncbi.nlm.nih.gov/pub/grc/human/GRC/Issue_Mapping/), regions blacklisted by ENCODE [83], and SDs annotations from the UCSC Table Browser. For T2T-CHM13, we included known T2T-CHM13v1.0 issues [30], and SD annotations from [32]. To obtain the number of genes impacted, we intersected each genomic feature to Gencode v35 and CATv4 gene annotations for GRCh38 and T2T-CHM13v1.0 respectively, using bedtools intersect [84] with default parameters.

#### Section II. T2T-CHM13 improves analysis of global genetic diversity based on 3,202 short-read samples from the 1KGP dataset

##### Short-read alignment, coverage analysis, and variant calling of the expanded 1KGP collection

In order to perform large-scale alignment and variant calling, we used the NHGRI Genomic Data Science Analysis, Visualization, and Informatics Lab-Space (AnVIL) [39], which allowed us to analyze nearly 100 terabytes of raw sequence data and perform thousands of analyses in just a few days of wall-clock time. Using the AnVIL/Terra platform, we aligned the 3,202 samples (**Fig. S2.1**) from the 1KGP [28] to the T2T-CHM13 reference genome using the functional equivalence pipeline standard established by the Centers for Common Disease Genomics (CCDG) project using GRCh38 [85] as a framework. For T2T-CHM13 we used assembled v1.0 (20200921) main chromosome sequences with an addition of chrY, chrY_KI270740v1_random, and chrEBV from GRCh38, whereas the prior analysis of the 1KGP samples used GRCh38 including alt, decoy, random, unknowns, and HLA sequences.

For our analysis, we followed the processing of the previous pipeline, although we updated certain elements of the pipeline in order to improve efficiency while maintaining accuracy. Most notably, we substituted samtools v1.11 [86] for Picard when fixing mate information in the BAM file, merging lane-level BAMs, marking duplicates, and coordinate-sorting. Otherwise, we followed the CCDG pipeline as used by the New York Genome Center (NYGC), including all pertinent parameters: lane-level alignment of the 1KGP FASTQ files to T2T-CHM13 using BWA-MEM v0.7.17-r1188 [40], fixing mate information in the BAMs, coordinate-sorting the BAMs, merging lane-level BAM files to sample-level BAM files, marking duplicates, and converting to CRAM format. Though the original pipeline performed base quality score recalibration at the BAM level, we forewent this step because the subsequent variant inference part of the pipeline utilizes a newer version of GATK framework (4.1.9 for AnVIL vs 3.5.0 for NYGC). This workflow is documented in **Fig. S2.2** and can be found at https://github.com/schatzlab/t2t-variants/blob/master/wdls/t2t_alignment.wdl. The NYGC workflow is documented here: https://ftp.1000genomes.ebi.ac.uk/vol1/ftp/data_collections/1000G_2504_high_coverage/working/20190425_NYGC_GATK/1000G_README_2019April10_NYGCjointcalls.pdf. When possible, we used the same flags and options as the original study at the NYGC used.

After alignment, we quantified the mapping statistics using samtools stats (**Fig. S2.4**). In addition to the improvements reported in **Fig. 2A**, several other metrics improved, such as a decrease in the standard deviation of the insert size and an improvement in the paired-end concordance. Lastly, because of the resolved/introduced repeat sequences in T2T-CHM13, we observe an increase in the number of reads with a mapping quality value of 0, signalling a repetitive alignment.

We then computed coverage statistics using mosdepth [87] on 500 bp non-overlapping genome-wide windows on individual sample BAM files (**Fig. S2.5**). For coverage analysis, we only considered autosomal chromosomes for coverage stability analysis, and removed from consideration 500 bp windows with 250+ Ns in the respective references (GRCh38: 260,251 windows, T2T-CHM13: 22,950 windows). We then consider the interquartile range to annotate 500 bp windows with outlier/abnormal coverage using a threshold of 1.5x IQR. We then intersected these regions against several annotations to evaluate coverage uniformity.

This included intervals defined by:

- Genome In A Bottle (GIAB):
    - HC_smallvar: High Confidence Small Variants
    - HC_SV: High Confidence Structural Variations
    - gene_medical: Medically relevant genes

- RepeatMasker:
    - r_LINE: LINE elements
    - r_SINE: SINE elements
    - r_low_complexity: Low complexity repeats
    - r_satellite: Satellite sequences
    - r_simple: Simple repeats
- CAT:
    - genic: Gene sequences
- SD:
    - synteny: Syntenic between GRCh38 and T2T-CHM13
    - Synteny_not: Not synthetic between GRCh38 and T2T-CHM13
- Superpopulation local ancestry derived from 1KG
    - a_AFR: African ancestry
    - a_AMR: American ancestry
    - a_EAS: East-Asian ancestry
    - a_EUR: European ancestry
    - a_SAS: South-Asian ancestry
    - a_neanderthal: Neanderthal ancestry

For each of these annotations, we computed the overlap of every stratification context with 500 bp windows for which the coverage was computed and retained context-specific windows with 1+/500 bp context overlaps.

***Variant Calling:*** To call small variants, we again followed the framework previously used, updating to GATK v4.1.9 [82]. Briefly, we performed raw variant calls using GATK HaplotypeCaller on a per-chromosome basis for each sample (https://github.com/schatzlab/t2t-variants/blob/master/wdls/t2t_haplotype_calling.wdl). We then generated GATK GenomicsDB files on each chromosome, across all 3,202 samples, per interval, with 1kb of padding on either side of each interval (https://github.com/schatzlab/t2t-variants/blob/master/wdls/t2t_genomics_db.wdl), performed joint genotyping using GATK GenotypeVCFs, and trimmed the padding on the resulting VCFs to produce interval-level joint VCF files across all 3,202 samples (https://github.com/schatzlab/t2t-variants/blob/master/wdls/t2t_interval_calling.wdl) [41]. We then used bcftools v1.9 [86] to concatenate the interval-level joint VCF files to produce chromosome-level VCF files and to obtain statistics on all VCF files. Lastly, we performed recalibration using GATK VariantRecalibrator (https://github.com/schatzlab/t2t-variants/blob/master/wdls/t2t_recalibration.wdl). This workflow is documented in **Fig. S2.3**, and the NYGC workflow is documented here: https://ftp.1000genomes.ebi.ac.uk/vol1/ftp/data_collections/1000G_2504_high_coverage/working/20190425_NYGC_GATK/1000G_README_2019April10_NYGCjointcalls.pdf. When possible, we used the same flags and options as previously used.

***Variant filtering and QC:*** To remove low quality, spurious variants, we first filtered these VCF files by the FILTER field, using only variants labeled as PASS by GATK. This eliminated a number of variants in highly repetitive regions, such as in satellite regions (**Fig. S2.6-7**). For most subsequent analysis, we only considered the 2,598 founder (unrelated) samples in the 1KGP. Using bcftools v1.9, we calculated the number of variants per sample, performed AF analysis on autosomes, and performed local ancestry analysis on the population-level VCF files. To identify variants with AF of 1.0 and of 0.5 +/ 0.05, we used bcftools v1.12 norm to stratify multiallelic variants and bcftools view. We also identified Mendelian discordance in all 602 trios using the MendelianViolationEvaluator evaluation module of GATK VariantEval.

#### Comparison of GRCh38/NY Genome Center Pipeline to T2T-CHM13/AnVIL Pipeline

The main differences between the original NYGC pipeline for short-read data processing and our AnVIL workflow using the T2T-CHM13 reference (**Fig. S2.2-3**) lie in: (i) tool/tool versions and (ii) the starting organization of the sample-specific read data. In contrast to the NYGC pipeline, which started with sequenced flow-cell- and lane-specific FASTQ files straight from the sequencing instruments, the AnVIL setup started from a composite FASTQ file downloaded from the European Nucleotide Archive. These files were then decomposed into individual flow-cell- and lane-specific FASTQ files.

For subsequent alignment steps, the AnVIL pipeline follows the NYGC pipeline, although uses a newer BWA v0.7.17 vs v0.7.15 in the NYGC pipeline and substitutes steps performed with Picard Tools in the NYGC pipeline (e.g., sorting, merging, deduplication) with samtools' analogs. Our AnVIL framework also forgoes the alignment recalibration step, as the subsequent variant inference part of the pipeline utilizes a newer version of GATK framework (4.1.9 for AnVIL vs 3.5.0 for NYGC), which utilizes elements of the alignment recalibration workflow directly during the variant inference.

Our AnVIL setup also utilized segmentation (1 Mbp) of the T2T-CHM13 reference with reciprocal overlapping (extra "extending" margins of 10kbp) between consecutive segments for the HaplotypeCaller step, followed by a joint genotyping step using GATK GenomicsDB. This was necessary because the default segmentation strategy for GRCh38 partitions the genome at "N" characters, but the T2T-CHM13 has nearly none such characters. Each genotyped segment-based variant file is then trimmed off of the extending 10 kbp extending margins on each segment, and the AnVIL pipeline concatenates segments in a sequential manner into chromosomes. The variant recalibration steps in our AnVIL setup follow the outline of the NYGC pipeline with the resource files lifted over from GRCh38 reference onto the T2T-CHM13, with only unambiguously lifted entries considered.

***Pipeline comparison:*** To evaluate how the alignment part of the pipeline differs between the AnVIL setup and the original NYGC pipeline, we performed GRCh38-based alignment with the AnVIL workflow for 3 arbitrarily selected samples. We then computed samtools alignment stats on both the original NYGC CRAM files and on the CRAM files generated on AnVIL. We observed that the alignments are almost identical across the two configurations, with most differences being very minor and observed in the number of MAPQ0 alignments (<0.2% reduction), number of reads marked "duplicated" (<0.01% reduction), insert size standard deviation (<1.2% reduction), pairs on different chromosomes (<1.2% reduction), and other metrics. (**Fig. S2.4**)

We further compared the AnVIL small-variant inference with the original NYGC workflow and observed genome-wide variant inference concordance of 98.7% (via bcftools stats). We further confirmed that the difference is driven by the GATK v3.5 in NYGC pipeline being updated to GATK 4.1.9 in AnVIL setup: identical versions of GATK run on alignments produced by the NYGC vs the AnVIL pipeline yielded >99.9% genome-wide concordance. We also observed that the proposed segmentation-with-margins GATK variant inferences approach utilized in the AnVIL setup yielded absolutely identical results when compared to complete-chromosome based segmentation on 200+ samples with T2T-CHM13v1.0 reference.

#### Visibility and concordance of SNVs when using different reference genomes for variant discovery

To determine the impact of using the T2T-CHM13 reference for variant calling and then annotating the results using resources available for GRCh38, we used the Picard LiftoverVCF tool to lift single-sample (HG002) and multi-sample (1000 Genomes Project samples, or "1KG") T2T-CHM13-based SNV calls over to the GRCh38 reference, and then compared the lifted variants to variant calls that were directly made using GRCh38. When lifted to GRCh38, the T2T-CHM13-derived single-sample callset for HG002 did not include 30.4% (1,051,004) of the 3,460,148 GRCh38-derived SNV calls made using the GRCh38 reference. This was largely due to the 946,126 sites in HG002 that are homozygous for the T2T-CHM13 allele, and therefore are called only when using GRCh38 as the reference. Likewise, the set of SNVs called against T2T-CHM13 and then lifted to GRCh38 contained an additional 721,547 SNVs, which were not called against GRCh38, and this was due largely to the 672,341 SNVs for which HG002 is homozygous for the GRCh38 allele. When we compare the remaining SNV sites for which HG002 has alleles, which are variant with respect to both reference genomes, the set of SNVs detected against T2T-CHM13 and then lifted to GRCh38 contains 95.8% of the SNV sites detected using GRCh38, and 98.0% of the SNV sites lifted from T2T-CHM13 are contained within the GRCh38 set.

While the difference in visibility of a single sample's variants with respect to the two reference sequences is significant, variant calls for larger multi-sample sets are less affected since large sets of samples are more likely to contain detectable variants with respect to both references in places where the two references exhibit different alleles. The 1KG multi-sample callset lacks far fewer of the GRCh38 calls due to the absence of an alternate allele with respect to T2T-CHM13: only 13,001 SNVs of the 106,581,271 GRCh38 1KG SNV calls were not detected with the T2T-CHM13 reference for this reason. Conversely, 12,124 1KG SNVs that were originally called using T2T-CHM13 were not detected when using GRCh38 due to a lack of 1KG samples with an allele that differs from the GRCh38 allele.

We performed analyses of the visibility of SNVs for single and multiple samples on the GATK short-read VCF files. For single-sample results, we used the calls from HG002, and for the 1KGP calls, the full set of variants for all 3,202 samples was used. To count the number of HG002 SNVs called using GRCh38 as a reference which are not callable using the T2T-CHM13 reference, the VCF file of HG002 GRCh38 SNVs was lifted to T2T-CHM13 using Picard v2.25.0's LiftOverVCF tool (http://broadinstitute.github.io/picard/) with the -RECOVER_SWAPPED_REF_ALT option. Any SNV whose genotype with respect to T2T-CHM13 was homozygous reference, or "0/0", and whose VCF record contained the Picard-assigned "SwappedAlleles" INFO tag, was counted among the 946,126 SNVs that are not callable against T2T-CHM13 due to the lack of a variant allele in HG002. Likewise, to calculate the number of HG002 SNVs called using T2T-CHM13 that lift over to GRCh38 positions for which the reference allele is not represented in HG002, we again ran Picard LiftOverVCF on the T2T-CHM13 VCF with the -RECOVER_SWAPPED_REF_ALT option. SNVs for which the lifted SNV's GRCh38 genotype was homozygous reference, or "0/0", and which were assigned the INFO field value "SwappedAlleles", were counted among the 672,341 SNVs that are not callable in GRCh38 due to the lack of a variant allele in HG002. To compute similar metrics for the 1KGP SNVs, the VCF files of T2T-CHM13 variant calls without genotypes were lifted from T2T-CHM13 to GRCh38, and the GRCh38 variants were similarly lifted from from GRCh38 to T2T-CHM13. Instead of examining genotypes, the lifted AFs (the "AF" values in the VCF INFO field) and the unadjusted alternate and total allele counts ("AC" and "AN") were examined using a custom perl script to count variants that were 100% fixed for one or the other reference allele.

To compare variants between the called and the lifted sets of variants for a given reference, we ran hap.py version 0.3.9 [88] on the complete sets of "PASS" variants from both references, as well as VCF files which had been filtered to remove variants visible only with respect to the T2T-CHM13 reference, only with respect to the GRCh38 reference, or visible with respect to both.

#### Section III. T2T-CHM13 improves structural variant analysis of 17 diverse long-read samples

##### Long-read alignment and coverage analysis

To assess the impact of using T2T-CHM13 as a reference on long-read alignment and variant calling, we aligned data from two independent sequencing platforms - PacBio HiFi and ONT long reads - to both GRCh38 and T2T-CHM13. For the GRCh38 reference, we excluded alternate and decoy sequences as current long-read mappers are not alt-aware, but included random and unknown sequences. For the T2T-CHM13 reference, we used the T2T-CHM13 v1.0 assembly with the addition of chrY, chrY_KI270740v1_random, and chrEBV from the GRCh38 reference.

We used HiFi sequencing data from 17 samples and ONT data from 14 of those samples (**Fig. 3a**). The average read length of HiFi data across the 17 samples was 18,130 bp, and the average read length of ONT data across the 14 samples for which it was available was 21,913 bp (**Fig S3.1**). We performed alignments using minimap2 v.217 [45] and Winnowmap v2.0.1 [44], and mapping statistics were compared between the two references using samtools stats (**Fig. S3.2**). The number of reads mapped was similar between the two references across all combinations of technology and aligner. However, compared to GRCh38, we observed a lower error rate in mapping when using T2T-CHM13 due to the more accurate and complete sequence. At the same time, alignments to T2T-CHM13 yielded more reads with a mapping quality of zero due to the increased number of resolved repeats which resulted in multiple identical targets for mapping. Finally, there were fewer non-primary alignments to T2T-CHM13 because the addition of reference sequence absent from GRCh38 enables better end-to-end primary alignments of long reads derived from these newly resolved regions.

In addition, we investigated the effect of the new reference on coverage anomalies by using mosdepth [87] to compute coverage statistics across non-overlapping 500 bp bins in each reference. Only autosomal chromosomes were considered, and bins with 250 or more N's were removed (260,251 bins in GRCh38 and 22,950 bins in T2T-CHM13).

We first computed the standard deviation of the per-bin coverage in each combination of reference, technology, aligner, and sample and compared the results between references. We observed a similar mean coverage in T2T-CHM13 and GRCh38, but found a significant reduction in the standard deviation when aligning to T2T-CHM13 (**Fig. S3.3**), indicating alignments distributed more uniformly across the genome.

In addition to these genome-wide results, we compared the mean and standard deviation among bins overlapping (≥1 bp) with a number of genomic contexts (**Fig. S3.3**):

- Satellite repeats
- Genes
- Non-syntenic regions with respect to the other reference
- Syntenic regions with respect to the other reference
- Abnormal coverage bins, or bins in which the coverage is outside the range [Median - 1.5 * (Median - Q1), Median + 1.5 * (Q3 - Median)] among all bins in the same reference with the same technology, aligner, and sample.

We found that the coverage standard deviation in GRCh38 was particularly elevated relative to T2T-CHM13 in satellite repeats, non-syntenic regions, and regions of abnormal coverage, likely due to repetitive sequences not properly characterized in GRCh38.

Finally, for those bins with abnormal coverage, we counted the number of distinct samples in which it had abnormal coverage (**Fig. S3.4**). We found that many abnormal coverage regions in T2T-CHM13 had such coverage in only a single sample, which could be caused by rare structural variation disrupting alignments to certain bins. On the other hand, GRCh38 had a higher number of bins with abnormal coverage in every sample, indicating incorrectly resolved repeats or other errors in the reference.

#### Long-read variant calling analysis

To evaluate the impact of T2T-CHM13 on structural variant (SV) calling from long reads, we called SVs in all long-read samples from the Winnowmap and minimap2 alignments described above. Variants were called separately in each unique combination of (sample, reference, technology, aligner) using Sniffles v1.0.11 [46] with sensitive parameters - a minimum variant length of 20 bp and a minimum read support of two reads. Variants were marked as high-confidence if they were at least 30bp in length, were annotated with the PRECISE INFO field by Sniffles, and were supported by a sufficient number of reads: at least 10 or 25% of that sample's average coverage, whichever is smaller. Then, we refined insertion calls using Iris and removed duplicate variant calls within the same callset using Jasmine v1.1.0 [47] with a constant breakpoint distance threshold of 200 bp. The alignments and variant calls were computed using the default recommended parameters from our optimized Jasmine-SV pipeline: https://github.com/mkirsche/Jasmine/tree/master/pipeline.

To compare SV calling between the two references in a large cohort setting, we constructed cohort-level SV calls with Jasmine from HiFi reads separately for each reference. We merged each sample's SV calls derived from both Winnowmap and minimap2 alignments, and only retained calls detected from both sets of alignments. Then, we merged the per-sample callsets to obtain a unified cohort-level SV callset. SVs not annotated as high-confidence in at least one sample in which they were present were discarded, resulting in final callsets of 124,566 SVs in GRCh38 and 141,193 SVs in T2T-CHM13. The cohort-level callset was computed using Jasmine with the default recommended parameters.

Using these cohort-level callsets, we computed the AF distribution in each reference (**Fig. 3C**) as well as the number of SVs present in each sample (**Fig. S3.5 and S3.6**). We also counted the number of insertions and deletions, respectively, in each reference to compute indel balance (**Fig. 3D**). To show the effects of SV calls in novel regions on the balance of insertions and deletions, we annotated SV calls present in non-syntenic regions of T2T-CHM13 with respect to GRCh38. We also looked specifically at the distributions of SV calls in T2T-CHM13 which intersected (1+ bp) a number of different genomic contexts, including genes and exons, syntenic and non-syntenic regions (**Fig. S3.7**), as well as centromeres and different classes of repeats (**Fig. S3.8**).

***Trio analysis:*** To evaluate the ability to detect *de novo* SVs in T2T-CHM13 in trio settings, we constructed trio callsets for two different trios from the Genome in a Bottle Consortium - the HG002 trio of Ashkenazim ancestry and the HG005 trio of Han Chinese ancestry. To construct a callset for each trio, we merged the callsets of the child and both parents derived from all four combinations of aligner (Winnowmap and minimap2) and sequencing technology (HiFi and ONT). Then, we discarded any variants not supported by both technologies with Winnowmap in at least one of the samples in which they were present. As in the population-level analysis above, we also only retained SVs which were annotated as high-confidence in at least one sample. This yielded final trio callsets of 56,414 variants in the HG002 trio and 56,449 variants in the HG005 trio. We inspected all SVs in IGV in these sets present in only the child of the trio (36 in HG002 and 45 in HG005 with respect to T2T-CHM13; 40 in HG002 and 29 in HG005 with respect to GRCh38). Similar analysis in GRCh38 yielded 40 candidates in HG002 and 29 candidates in HG005. In both references, these candidate sets include a previously unreported potential *de novo* variant in HG005, a 1,571 bp deletion at chr17:49,401,990 in T2T-CHM13 (**Fig. S3.9**).

#### Optical mapping assembly and variant calling

We generated Bionano optical mapping data for HG002/GM24385 using the DLS chemistry and Bionano Saphyr system, available at https://ftp-trace.ncbi.nlm.nih.gov/ReferenceSamples/giab/data/AshkenazimTrio/analysis/Bionano_haplotype_SV_DLS_06172019/. A haplotype-aware optical assembly of Bionano data for HG002 was generated using default parameters with Bionano Solve 3.6. We aligned the assembly against both the GRCh38 and T2T-CHM13v1.0 references, and performed

SV calling using Bionano Solve3.6 with default parameters. For mapping, the T2T-CHM13 reference was *in silico* digested from the sequence t2t-chm13.20200921.withGRCh38chrY.chrEBV.chrYKI270740v1r.fasta.

There were 2865 non-redundant variants called against T2T-CHM13, and the number of insertions and deletions >500 bp in size called against T2T-CHM13 are more balanced than those against GRCh38, consistent with the observation that T2T-CHM13 corrects collapses in GRCh38 (**Fig. S3.14**). There are 1431 insertions called against the T2T-CHM13 reference compared to the 2771 insertions called against GRCh38.

Among the variants called against the T2T-CHM13 reference, 306 of them overlap non-syntenic regions. Some of these SVs involve gaps in GRCh38 that are fixed in the T2T-CHM13 reference (**Fig. S3.15**). Although the SVs crossing gaps in GRCh38 can still be called with respect to GRCh38, T2T-CHM13 improves resolution for these SVs by adding markers that can be aligned between the optical assembly and the reference. Future work could include evaluating the utility of calling SVs from Bionano data in many individuals against T2T-CHM13 to assess improvements across ancestry groups.

#### Section IV. Novel variants and evolutionary signatures within newly accessible regions of the genome

##### Newly added region analysis

We intersected the 1KGP population-wide joint genotyped VCF files with the non-syntenic and novel regions of the T2T-CHM13 genome, which we used to calculate the number of bases newly added in this assembly and the number of variants in these newly accessible regions. We also intersected these regions with the T2T-CHM13 v4 gene annotations to get the number of genes intersecting with novel regions as well as the number of newly accessible variants within protein-coding genes.

##### Identifying concordance between PacBio HiFi, ONT and Illumina variant calls

To evaluate the variant calls in non-syntenic regions, we derived concordance between variant calls generated with HiFi, ONT, and Illumina reads. For each sample, we used bcftools to filter the non-PASS and uncalled variants from each variant callset. We then used hap.py [88] to derive the precision, recall, and F1-score between each variant callset two determine how many variants are common between each pair of sets. We restricted this analysis to autosomes and only used SNP calls, excluding indels in the pre-processing step.

In each comparison, we set HiFi as the baseline and Illumina or ONT as query VCFs. The reported true positive events are common variants between two platforms, false positives are variants missing from the PacBio HiFi platform, and false negatives are variants present in PacBio HiFi platform but missing from the query platform.

##### Distance and LD between non-syntenic regions and phenotype-associated SNVs

We used bedtools v2.3 intersect [84] with default parameters to identify those regions in the T2T-CHM13 assembly that are annotated as both novel and non-syntenic, and then used bedtools subtract with default parameters to identify regions that were annotated only as novel or only as non-syntenic. With these three non-overlapping regions, we used bedtools closest with the "-t first" argument to find the closest GWAS Catalog [50] and Clinvar variants [81] (lifted over from the GRCh38 assembly to the T2T-CHM13 assembly) to each of these regions.

The proximity of some GWAS Catalog and ClinVar SNVs to non-syntenic regions suggests that these regions may contain variation of phenotypic and medical relevance. To more directly address this question, we used short-read variant calls from the 1KGP cohort to investigate patterns of LD between SNVs from the GWAS Catalog (https://www.ebi.ac.uk/gwas/home) [50] and variants in non-syntenic regions. Non-syntenic variants in strong LD with GWAS hits represent candidate causal variants that may have been inaccessible to previous studies that used GRCh38 or earlier assemblies. We chose not to include the ClinVar database in this analysis because putatively pathogenic variants should segregate at very low AFs and are unlikely to be represented in the 1KGP individuals.

Using v1.0.2 of the GWAS Catalog's list of associations and v1.0 of its ancestry metadata, we subsetted the database to studies conducted in populations that broadly corresponded to a set of 1KGP populations. This remaining set of associations comprised 902 of the 4,422 total studies in the catalog and 30,181 of the 257,947 total SNVs.

| GWAS catalog broad ancestry | Corresponding 1KGP populations |
|---|---|
| European | CEU, FIN, GBR, IBS, TSI |
| East Asian | CHS, CHB, JPT |
| African American or Afro-Caribbean | ACB, ASW |

| | |
|---|---|
| South Asian | BEB, GIH, ITU, PJL, STU |
| Hispanic or Latin American | CLM, MXL, PEL, PUR |
| Hispanic or Latin American, Native American | CLM, MXL, PEL, PUR |
| Native American | CLM, MXL, PEL, PUR |
| Sub-Saharan African | ESN, GWD, LWK, MSL, YRI |

We matched the rsIDs of SNVs identified in this subset of studies to rsIDs in the NCBI dbSNP database, lifted over to T2T-CHM13. Based on position, we then matched these lifted over GWAS variants to variants called in the 1KGP dataset. Using PLINK v1.90b6.21 [89], we calculated LD between each GWAS variant and all non-syntenic variants within a 1 Mbp window, reporting only non-syntenic variants in strong LD ($R^2 \geq 0.5$).

#### Identifying frequency-differentiated variants in non-syntenic T2T-CHM13 regions

Within regions of the T2T-CHM13 assembly that are non-syntenic with GRCh38, we searched for variants with large AF differences between populations, which may represent evolutionarily interesting loci that were inaccessible using GRCh38. Using PASS filtered variant genotypes generated with the 1KGP dataset, we applied Ohana [51,90,91], a maximum likelihood-based method that models individual genomes as possessing ancestry from combinations of *k* ancestry components. Ohana then tests whether individual variants adhere to this genome-wide null model, or are better explained by an alternative model in which their AF differs in one ancestry component. Outlier loci identified through this method exhibit extreme levels of frequency differentiation compared to the genome-wide average.

We used the inferred admixture relationships generated by [52] for the core 2,504 samples from the 1KGP, modeling these individuals' genomes as combinations of 8 ancestry components. This choice of *k* follows the precedent of the 1KGP [18] and replicates known patterns of population structure at continental scales, as well as expected signatures of admixture within the Ad Mixed American and African American populations (**Fig. S4.5**). In order to generate "selection hypothesis" matrices to search for selection in a specific ancestry component, we modified the neutral covariance matrix by allowing one component at a time to have greater covariance. A scalar value of 10, representing the furthest possible deviation that a variant could have in a population under selection, was added to elements of the neutral covariance matrix depending on the population of interest.

For each variant of interest, Ohana computes the likelihood of the observed ancestry component-specific AFs under both the selection and neutral models, then compares the two by computing a likelihood ratio statistic (LRS), which quantifies relative support for the selection hypothesis. We filtered these results to remove extreme outliers in null model log-likelihoods (global log likelihood estimate (LLE) < -1000), which were unremarkable in their patterns of AF and instead indicated a failure of the neutral model to converge for a small subset of variants. We also required variants to be genotyped in at least 95% of samples. Using PLINK v1.90b6.21 [89], we calculated population-specific AFs and removed variants that were extremely rare in all 1KGP populations (minor AF in all populations < 0.05 or > 0.95).

We intersected variant positions with the T2T-CHM13 v4 gene annotations to identify variants that overlapped with genes and exons, and we identified outliers for each ancestry component by selecting variants with LRS values in the 99.9th percentile.

#### Comparison of frequency-differentiated variants between T2T-CHM13 and GRCh38

Next, we further characterized the 5,154 most highly differentiated variants across all 8 ancestry components by comparing them to variants called on GRCh38. Using the UCSC Genome Browser's liftOver tool [71], we were able to lift over 3,038 (58.9%) of these 5,154 variants from T2T-CHM13 to GRCh38.

We intersected the remaining 2,116 variants that could not be lifted over with T2T-CHM13 "novel regions," a stricter set of novel sequences in the T2T-CHM13 genome that do not align to GRCh38 using Winnowmap. In order to further subset to regions accessible for short-read variant calling, we also restricted our analysis to variants in regions mappable by unique 100-mers (see **Section I: Mappability Analysis**). This left 943 unique variants that are likely to be truly novel variant calls made on T2T-CHM13. We manually verified the novel variant calls highlighted in **Fig. 4F-G** by subsetting the adjacent region in the CRAM file with samtools v1.12 [86] and visualizing read alignments with IGV v2.8.2 [92].

For the 3,038 variants that successfully lifted over to GRCh38, we were able to directly compare their selection likelihood ratio statistics (LRS) computed by Ohana to LRS values computed for GRCh38 variant calls in Yan *et al*. [52]. The GRCh38 variant calls were generated by the 1KGP Consortium, by aligning the 1KGP Phase 3 data to GRCh38 and calling variants, and are restricted to biallelic SNVs and indels (http://ftp.1000genomes.ebi.ac.uk/vol1/ftp/data_collections/1000_genomes_project/release/20190312_biallelic_SNV_and_INDEL/). LRS values for these variants were then calculated with Ohana using the same 2,504 samples and matrix of admixture relationships as in this study. These identical methods of scoring frequency-differentiated variants allow for a direct comparison of LRS values between variants on T2T-CHM13 and GRCh38.

For each of the highly differentiated non-syntenic variants that lifted over to GRCh38, we identified all GRCh38 variants that were 1 kbp upstream or 1 kbp downstream of the lifted over coordinate. 2,302 variants had at least one GRCh38 variant within this window. In order to determine whether the window contained SNPs with AF differentiation signals that would have been recognized on GRCh38, we then checked the LRS values for GRCh38 variants in the same ancestry component (**Fig. S4.6**). Setting our threshold to:

$$\text{GRCh38 LRS} \geq \text{T2T-CHM13 LRS - 10},$$

we found that there were 1,255 T2T-CHM13 variants where neighboring GRCh38 variants had comparable or greater LRS values, suggesting that these loci could have been identified in studies using the GRCh38 assembly.

Interestingly, we observed many cases where the lifted over T2T-CHM13 variant matched nearly identically to a GRCh38 variant in both position and LRS score, indicating that these likely represent the same variant between assemblies (**Fig. S4.5**). This redundancy is a consequence of the broadness of our definition of non-syntenic regions, which capture many sequences that are also represented in GRCh38, and demonstrates the reproducibility of LRS scores generated by Ohana.

#### Collapsed duplications and false duplication variant comparisons

To count the number of variants per gene, we used GRCh38 Genecode (v35) and lifted over using Picard v2.25.0's LiftOverIntervalList tool to identify corresponding locations in T2T-CHM13v1.0 and bedtools coverage intersected with VCFs of 1KGP variants from GRCh38 and T2T-CHM13. We calculated variant density as the number of variants in the interval divided by gene size, multiplied by a factor of 1000. Using only lifted over gene positions, we calculated statistical differences in the density distributions between references using a Wilcoxon signed rank test (wilcox.test function in R).

#### Section V. Impact of T2T-CHM13 on Clinical Genomics

##### Annotation of T2T-CHM13 variants using Variant Effect Predictor (VEP)

VEP [54] (version 102.0) was used to annotate variants generated by dipcall [53] when aligning the T2T-CHM13 reference genome (chm13_v1.0_plus38Y.fa) to the GRCh38 reference genome (hg38.no_alt.fa). VCF files were annotated without the --filter_common and --canonical flags. CADD [93] v1.6 and raw SpliceAI [94] scores were added using both the CADD and SpliceAI plugins. Variants were filtered based on predicted HIGH functional impact.

##### Identification of medically relevant genes with impacted variant discovery

Reference artifacts and errors were intersected with a previously curated list of medically relevant genes [23]. GRCh38 coordinates for genes were lifted over using Picard v2.25.0's LiftOverIntervalList tool to identify locations in T2T-CHM13v1.0. For SVs, we intersected gene coordinates after expanding breakpoints for each variant ± 5 kbp.

***TNNT3 analysis***: Genomic sequencing containing *TNNT3* implicated with SVs in GRCh38 were extracted from each reference (chr11:1,892,362-1,946,566, GRCh38; chr11:1,978,257-2,034,355, T2T-CHM13v1.0) and homologous regions compared using miropeats -s 400 [95].

***KCNJ18 analysis:*** SDs containing *KCNJ18* were identified in GRCh38 (chr17:21,687,227-21,736,311; CDS: chr17:21,702,787-21,704,088) and T2T-CHM13v1.0 (chr17:21,636,382-21,685,461; CDS: chr17:21,651,942-21,653,243) and paralogs pinpointed using the UCSC Genome Browser SD annotations [96–98]. With *KCNJ18*, Genome coordinates of SDs containing *KCNJ12* (GRCh38: chr17:21,399,778-21,450,415 (whole) and chr17:21,415,343-21,416,644 (CDS); T2T-CHM13v1.0: chr17:21,348,977-21,399,639 (whole) and chr17:21,364,558-21,365,859 (CDS)) and the novel *KCNJ17* (T2T-CHM13v1.0: chr17:22,634,421-22,683,415 (whole) and chr17:22,666,582-22,667,880 (CDS)) were used to extract 1KGP AFs from GRCh38-called and T2T-CHM13-called variants intersecting each locus. Histograms of the minor-allele frequencies were plotted and distributions compared using a Mann-Whitney U test (wilcox.test R function, with unpaired and two-sided settings). Finally, direct comparison of AFs for variants falling within the CDS were compared using 1KGP datasets from the NYGC (GRCh38), produced here (described above for T2T-CHM13), and the liftover of T2T-CHM13 to GRCh38 (described above).

##### Creating a difficult medically relevant gene benchmark for HG002 on T2T-CHM13

The Genome in a Bottle Consortium (GIAB) recently created a highly-curated benchmark for 273 challenging and medically relevant genes with GRCh37 and GRCh38 as references [23]. This benchmark used variant calls generated by dipcall [53] when aligning the paternal and maternal haplotypes assembled using hifiasm with PacBio HiFi reads for HG002 and Illumina k-mers from the parents, HG003 and HG004 [99]. Calling variants in these 273 genes tends to be challenging due to structural variants, SDs, and/or reference errors. The final benchmark regions for these genes excluded a small number of errors in homopolymers longer than 20 bp and errors found during extensive manual curation.

Here we create a comparable benchmark for HG002 on the T2T-CHM13 reference with the following steps:

1. We find the gene coordinates on T2T-CHM13 for the 273 genes using the T2T-CHM13 v3 gene annotations.
2. We generate variant calls with dipcall using the same HG002 trio-hifiasm assembly with T2T-CHM13 as a reference, as in the NIST assembly benchmarking pipeline (https://github.com/usnistgov/giab-asm-benchmarking). Dipcall generates variant calls using any non-reference support in regions that are ≥50 kbp with contigs having mapping quality ≥5. Dipcall also produces a BED denoting confident regions covered by an alignment ≥50 kb with contigs having mapping quality ≥5 and with no other >10 kbp alignments.

3. We identify that dipcall/hifiasm fully resolves 269/273 genes on T2T-CHM13 (i.e., the gene region and any SDs overlapping the gene region are encompassed by the dip.bed file output by dipcall, so that one and only one contig from each haplotype fully covers the region). We exclude 4 genes (*ESPN*, *IFITM3*, *FCGR3A*, and *FCGR2B*) due to large structural variation in T2T-CHM13 relative to GRCh38 and HG002.
4. We lift over the HG002 medically relevant genes v1.0 small variant benchmark regions from GRCh38 to T2T-CHM13 and intersect with T2T-CHM13 gene coordinates.
5. We manually exclude new SVs >49bp in HG002 on T2T-CHM13 as well as any overlapping tandem repeats.
6. We lift over T2T-CHM13 benchmark regions from T2T-CHM13 to GRCh38 and intersect with GRCh38 benchmark regions to get equivalent benchmarks on both references.

The resulting benchmark regions are <1% different in size, 11,548,799 bp and 11,482,860 bp on GRCh38 and T2T-CHM13, respectively, enabling benchmarking performance metrics to be directly compared between the references.

To compare variant call accuracy when using T2T-CHM13 vs. GRCh38 as a reference, we follow the best practices for benchmarking from the Global Alliance for Genomics and Health (GA4GH), using hap.py with vcfeval to generate performance metrics [88]. We use 3 callsets for HG002 generated similarly for this work on both GRCh38 and T2T-CHM13: Illumina-BWA-MEM-GATK, HiFi-PEPPER-DeepVariant, and ONT-PEPPER-DeepVariant. For Illumina-BWA-MEM-GATK, we use single-sample variant calls before filtering, because HG002 was not included in the 1KGP population analysis for GRCh38.

The challenging medically relevant gene benchmark variants and regions for HG002 on GRCh38 and T2T-CHM13, as well as dipcall results for HG002 on both references, are available under https://ftp-trace.ncbi.nlm.nih.gov/ReferenceSamples/giab/release/AshkenazimTrio/HG002_NA24385_son/CMRG_v1.00/. To have a benchmark on GRCh38 that is directly comparable to the T2T-CHM13 benchmark, the GRCh38 bed file https://ftp-trace.ncbi.nlm.nih.gov/ReferenceSamples/giab/release/AshkenazimTrio/HG002_NA24385_son/CMRG_v1.00/CHM13v1.0/SupplementaryFiles/HG002_CHM13_CMRG_smallvar_v1.00_GRCh38-equiv-regions_draft.bed can be used with the GRCh38 VCF at https://ftp-trace.ncbi.nlm.nih.gov/ReferenceSamples/giab/release/AshkenazimTrio/HG002_NA24385_son/CMRG_v1.00/GRCh38/SmallVariant/HG002_GRCh38_CMRG_smallvar_v1.00.vcf.gz.

### Figure 1. Supplemental

Structural comparisons of GRCh38 and T2T-CHM13

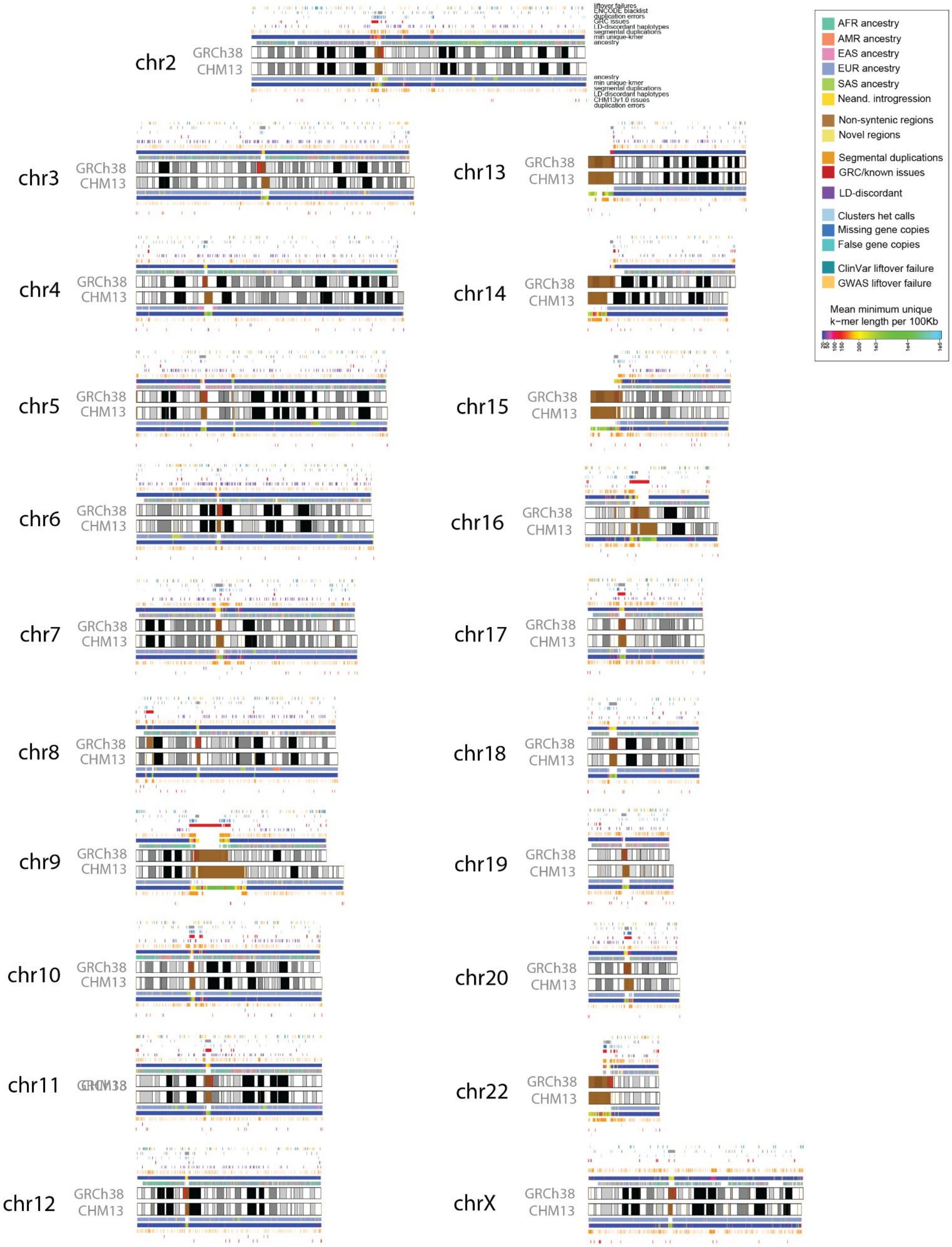


**Fig. S1.1 Annotation of all chromosomes**

Overview of annotations available for GRCh38 and CHM13 all chromosomes (chromosomes 1 and 21 shown in Figure 1A) with colors indicated in legend. Cytobands are pictured as gray bands with red bands representing centromeric regions within ideograms.

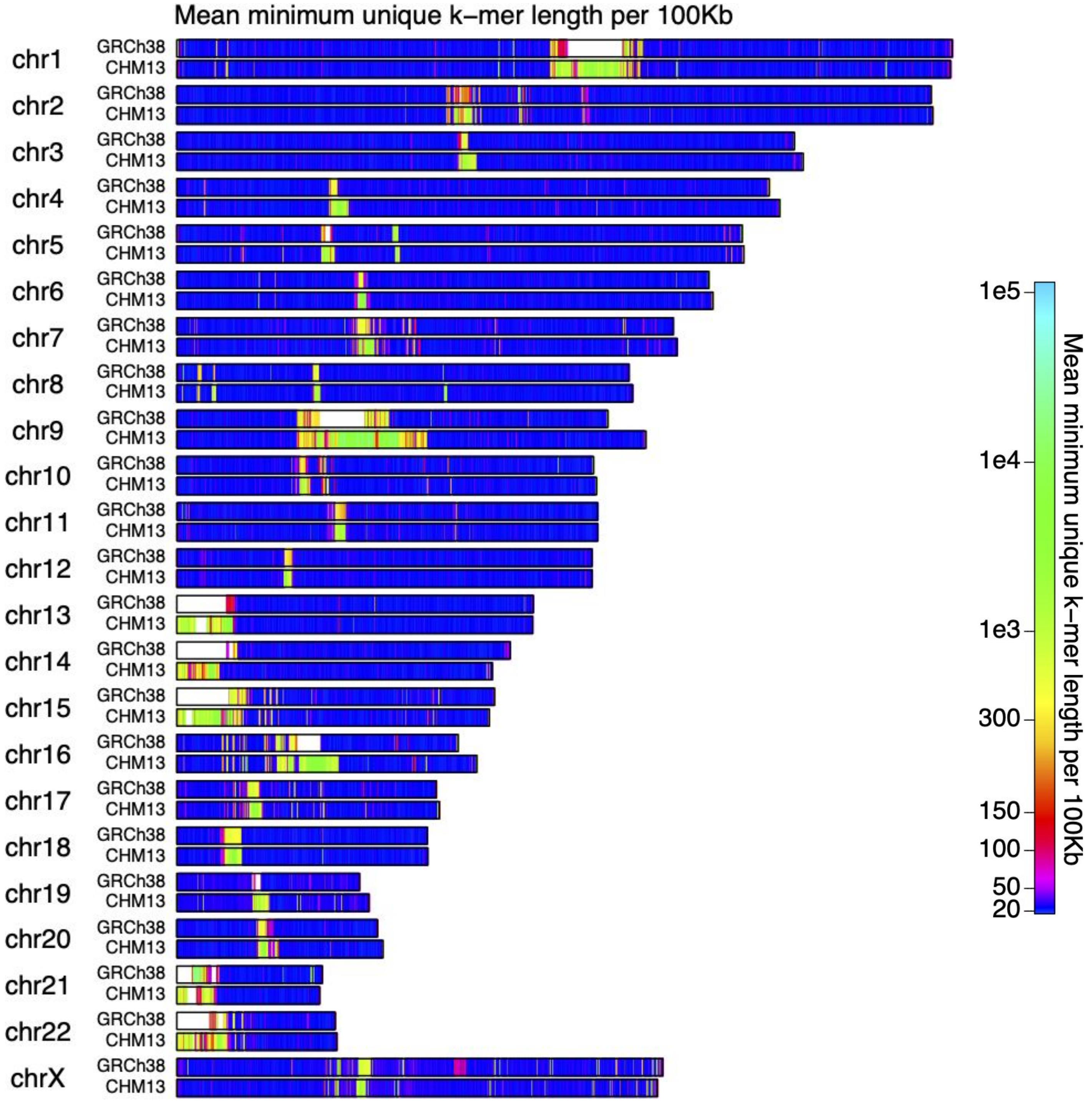


**Fig. S1.2 Minimum unique k-mer chromosome score maps**

Minimum unique k-mer scores (left-anchored) were averaged in 100 Kb bins and plotted along the length of each chromosome. Regions denoted in white indicate that no valid score exists, either because the k-mer sequence contains at least one N or the length of the k-mer would cause it to overlap a chromosome end.

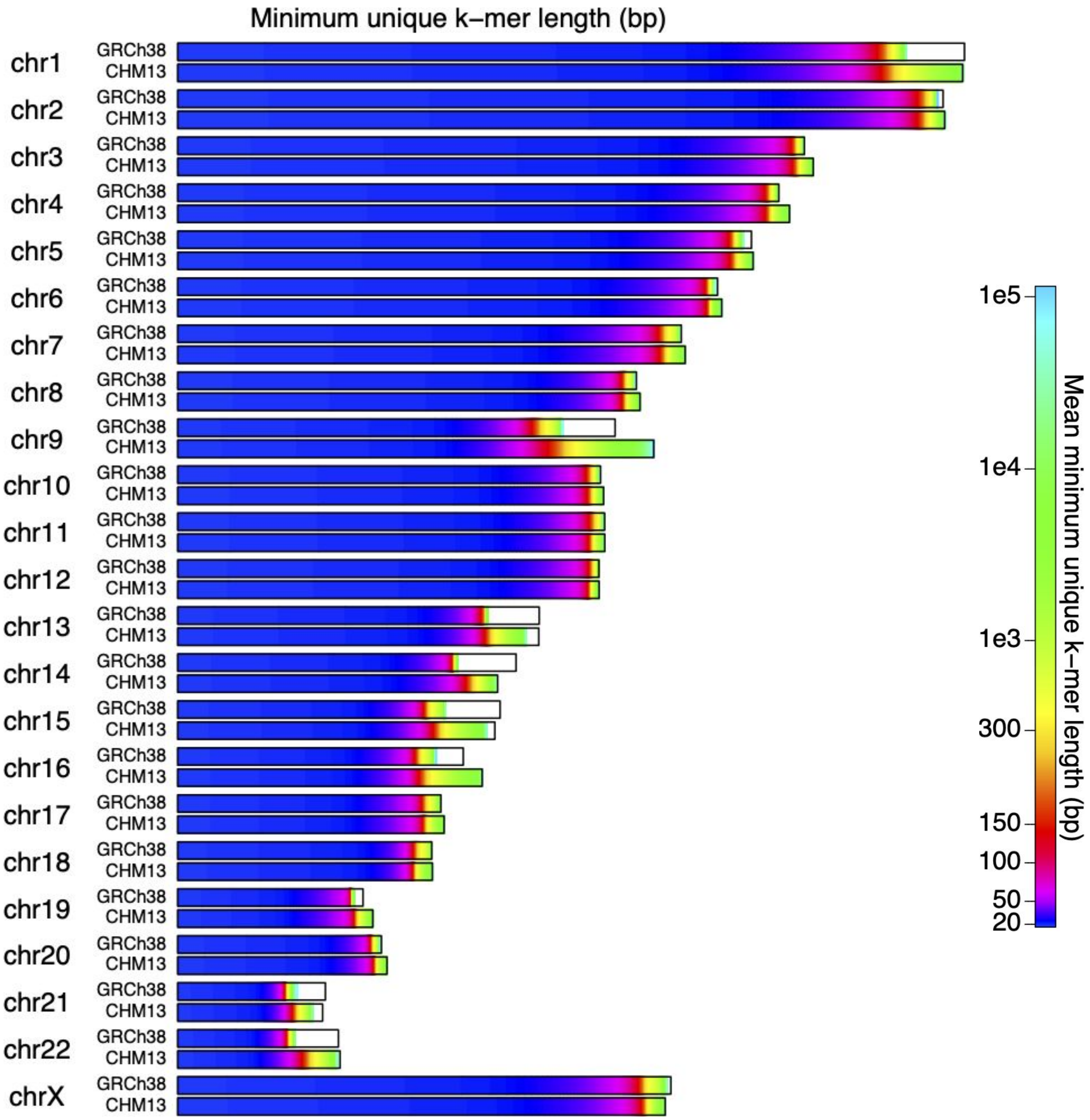


**Fig. S1.3 Minimum unique k-mer chromosome histograms**

Minimum unique k-mer scores (left-anchored) were calculated for each base position, sorted on a per-chromosome basis, and plotted. A black outline represents the reported length of each chromosome. Regions of white indicate invalid scores due to either overlapping sequence containing at least one N or overlapping the end of a chromosome.

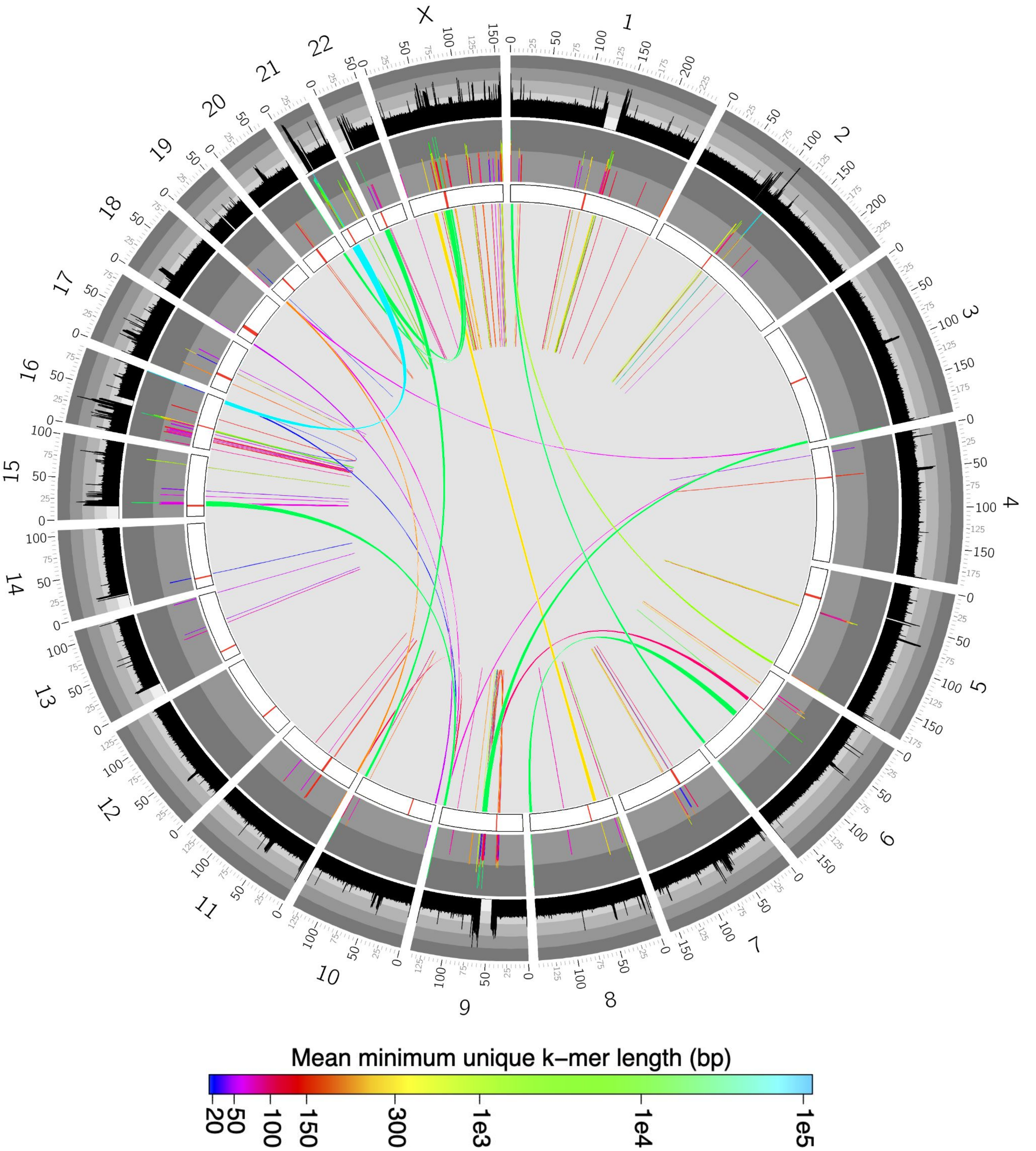


**Fig. S1.4 Large exact match sequence pairs in GRCh38**

For all minimum non-unique sequences greater than 5 Kb (minimum unique k-mer length plus one), the pair of positions corresponding to each sequence were determined. Sequence sizes are denoted by both color and a colored barplot (middle ring) that ranges from 1 Kb to 100 Kb in $\log_{10}$ scale. The inner ring denotes the relative length of each chromosome with the annotated centromeric region indicated in read. The outer ring shows the minimum unique k-mer score, binned at 100 Kb intervals and is shown with a range of 1 to 100 Kb in $\log_{10}$ scale.

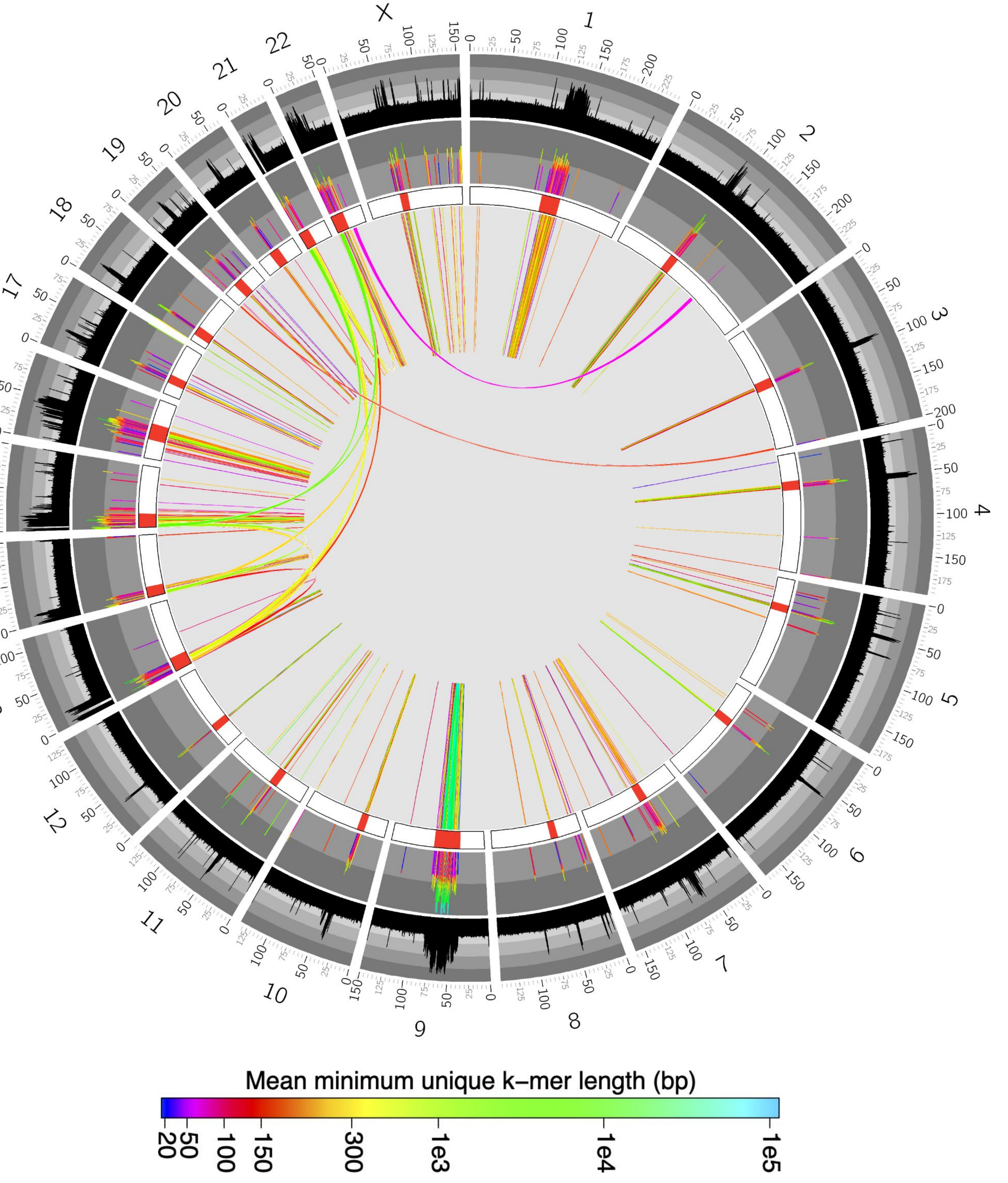


**Fig. S1.5 Large exact match sequence pairs in CHM13**

For all minimum non-unique sequences greater than 5 Kb (minimum unique k-mer length plus one), the pair of positions corresponding to each sequence were determined. Sequence sizes are denoted by both color and a colored barplot (middle ring) that ranges from 1 Kb to 100 Kb in $\log_{10}$ scale. The inner ring denotes the relative length of each chromosome with the annotated centromeric region indicated in read. The outer ring shows the minimum unique k-mer score, binned at 100 Kb intervals and is shown with a range of 1 to 100 Kb in $\log_{10}$ scale.

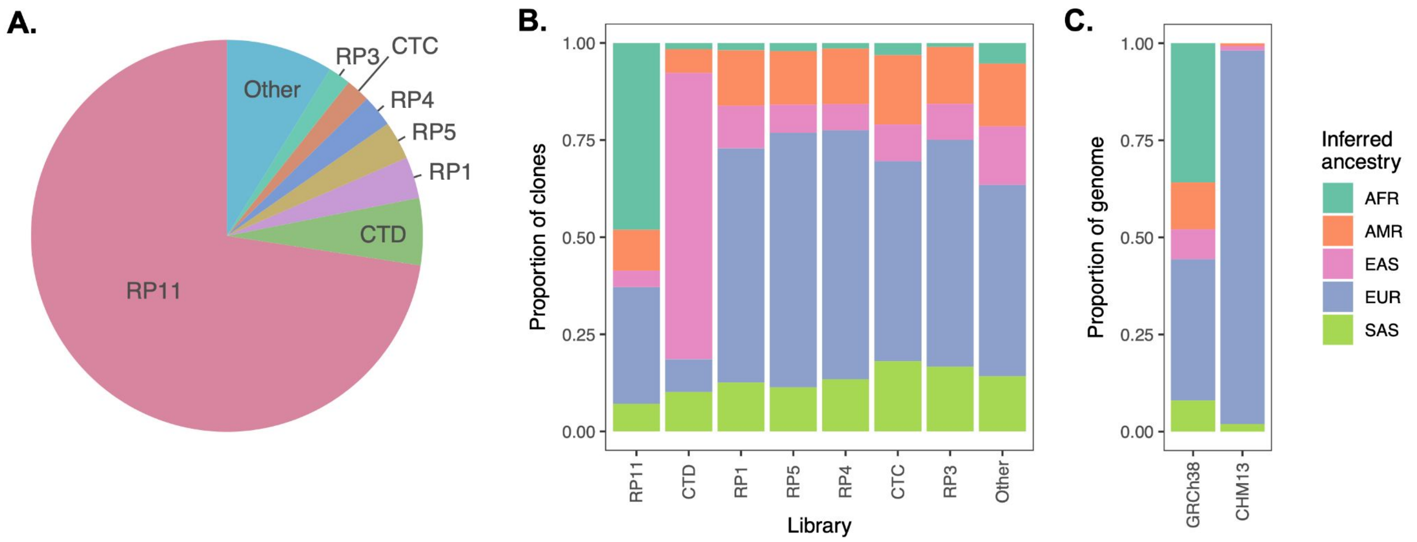


**Fig. S1.6. Local ancestry analysis of GRCh38 and CHM13**

**A.** Proportion of the GRCh38 reference genome composed of clones from various libraries, each derived from DNA obtained from a distinct diploid donor. **B.** Inferred local ancestry proportions for BAC libraries derived from different donor individuals that contributed to GRCh38. **C.** Total inferred local ancestry proportions for the GRCh38 and CHM13 reference genome.

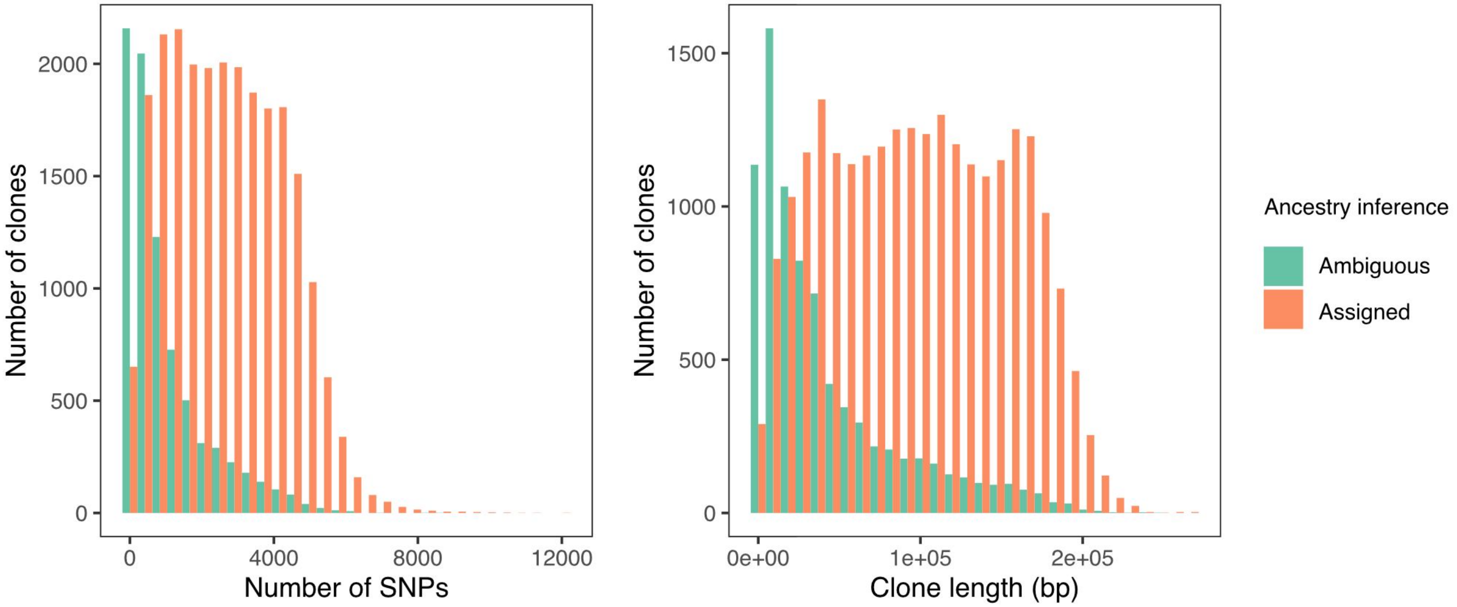


**Fig. S1.7. Ambiguity in ancestry inference for short GRCh38 clones with few markers**

Number of SNPs (left panel) and length (right panel) of each GRCh38 clone for which ancestry was or was not inferred based on majority vote of nearest neighbor haplotypes in the phased 1000 Genomes reference panel.

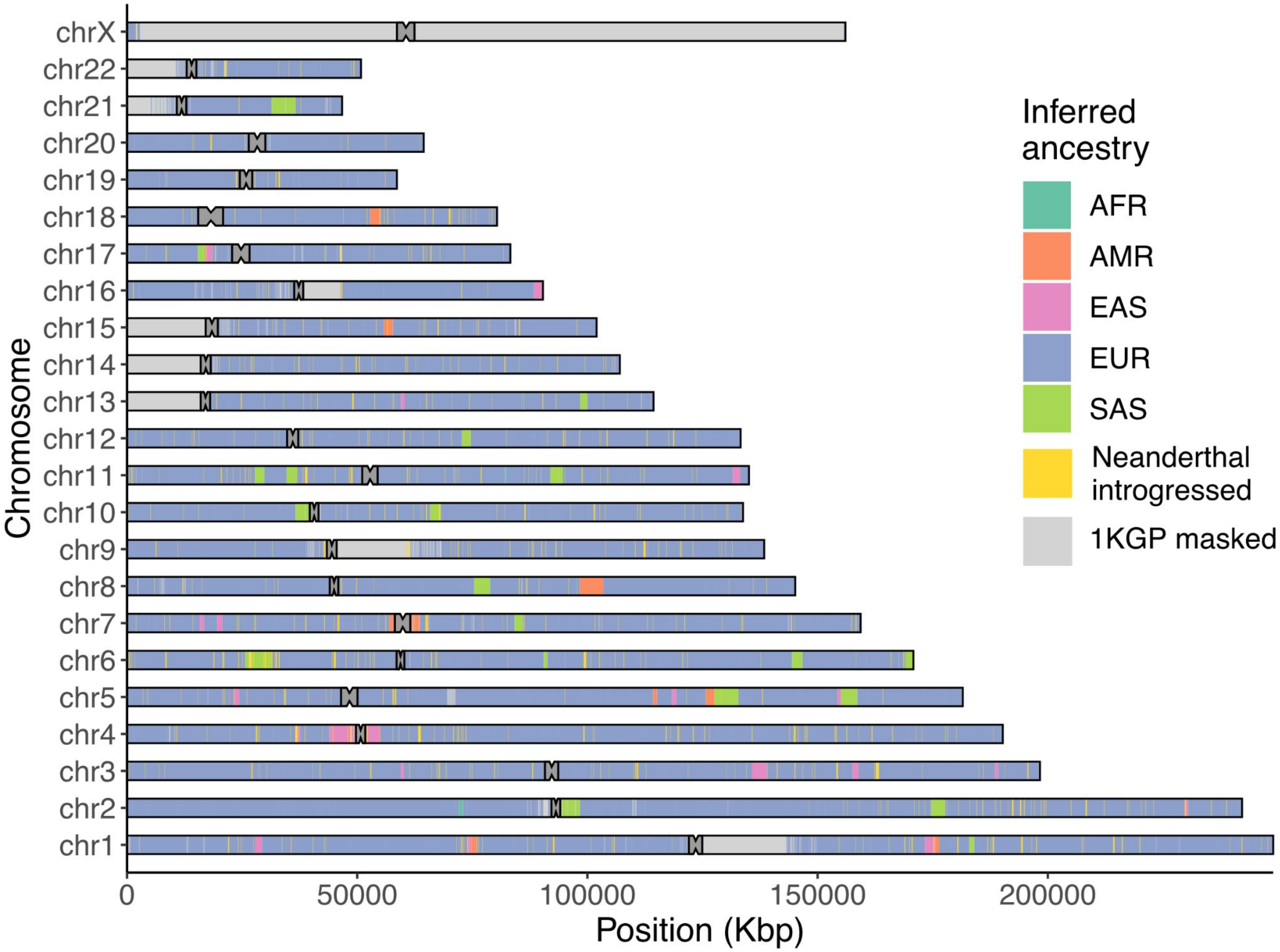


**Fig. S1.8. Local ancestry analysis of CHM13**

Ideogram depicting RFMix-inferred local ancestry tracts for CHM13. Neanderthal introgressed haplotypes, inferred with IBDMix, and regions masked by the 1000 Genomes Project (1KGP) are superimposed.

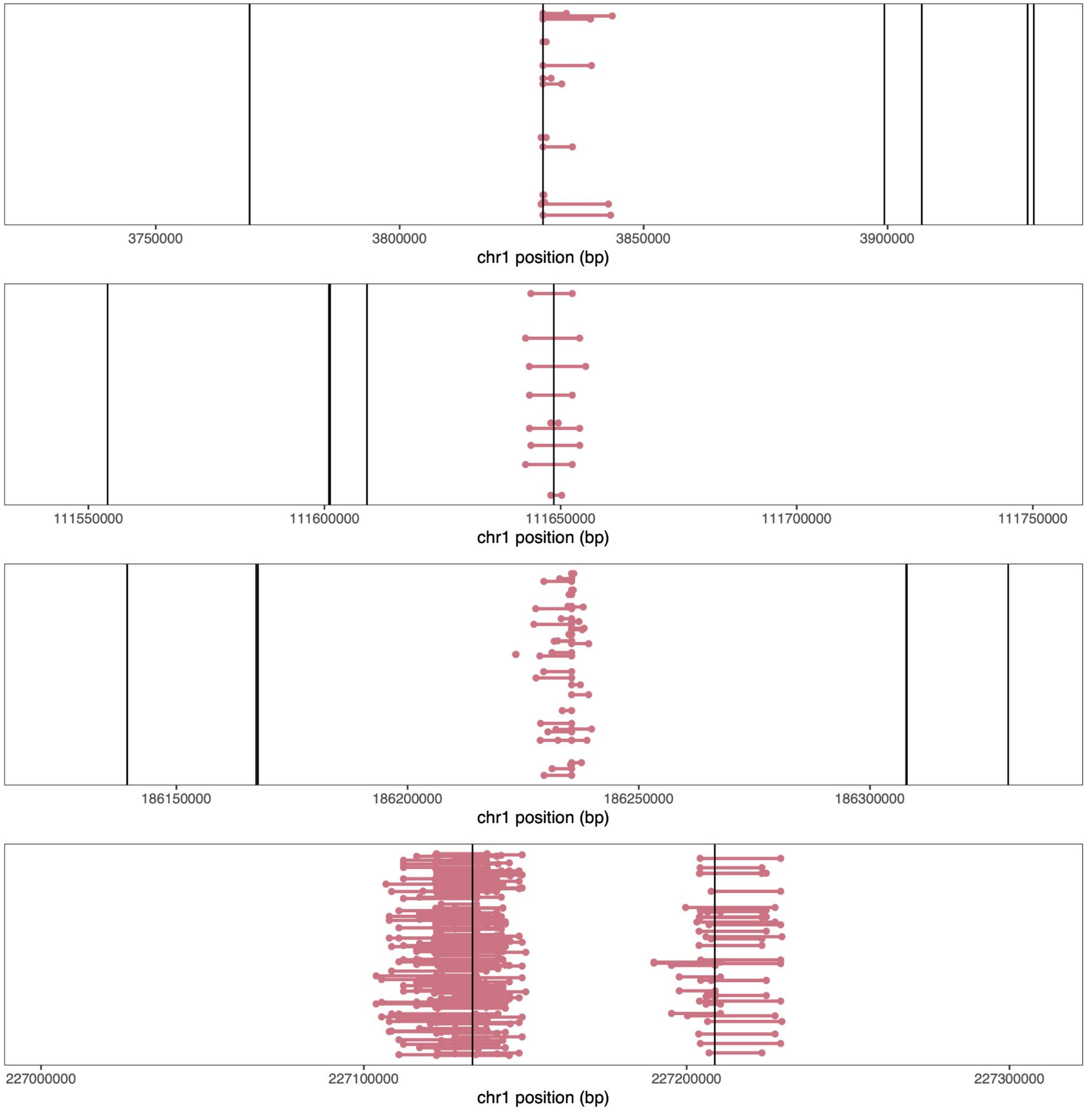


**Fig. S1.9. LD-discordant SNP pairs frequently span clone boundaries**

Depiction of four representative "islands" of LD-discordant SNP pairs, where for common SNPs in perfect LD ($R^2$ = 1), GRCh38 possesses a combination of alleles that is never observed among the 1000 Genomes sample. Linked SNP pairs are represented as dots connected by lines. Clone boundaries are represented as vertical lines. In all but one case (third row), SNP pairs straddle the annotated boundary of BAC clones.

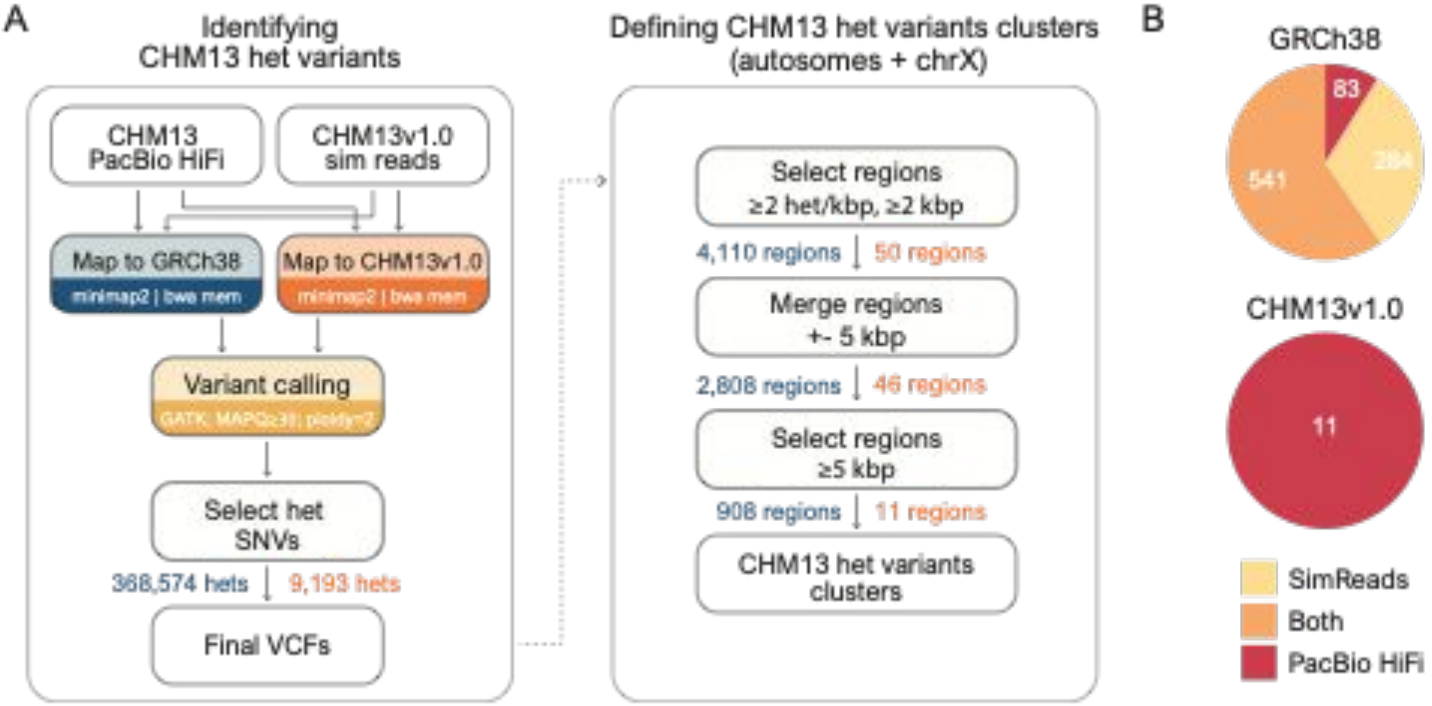


**Fig. S1.10. Strategy used to detect CHM13 heterozygous variant clusters**

(**A**) The workflow to identify CHM13 heterozygous false positive (FP) heterozygous (het) regions in GRCh38 (blue) and CHM13v1.0 + Y chromosome (red) from Illumina simulated reads (from T2T-CHM13v1.0 + Y chromosome) and PacBio (PB) HiFi reads generated from the CHM13 cell line. Numbers of features (excluding those associated with chrY) are indicated after each step. (**B**) The sequencing-platform source of the FP het regions are shown in a Venn diagram for GRCh38 and CHM13v1.0.

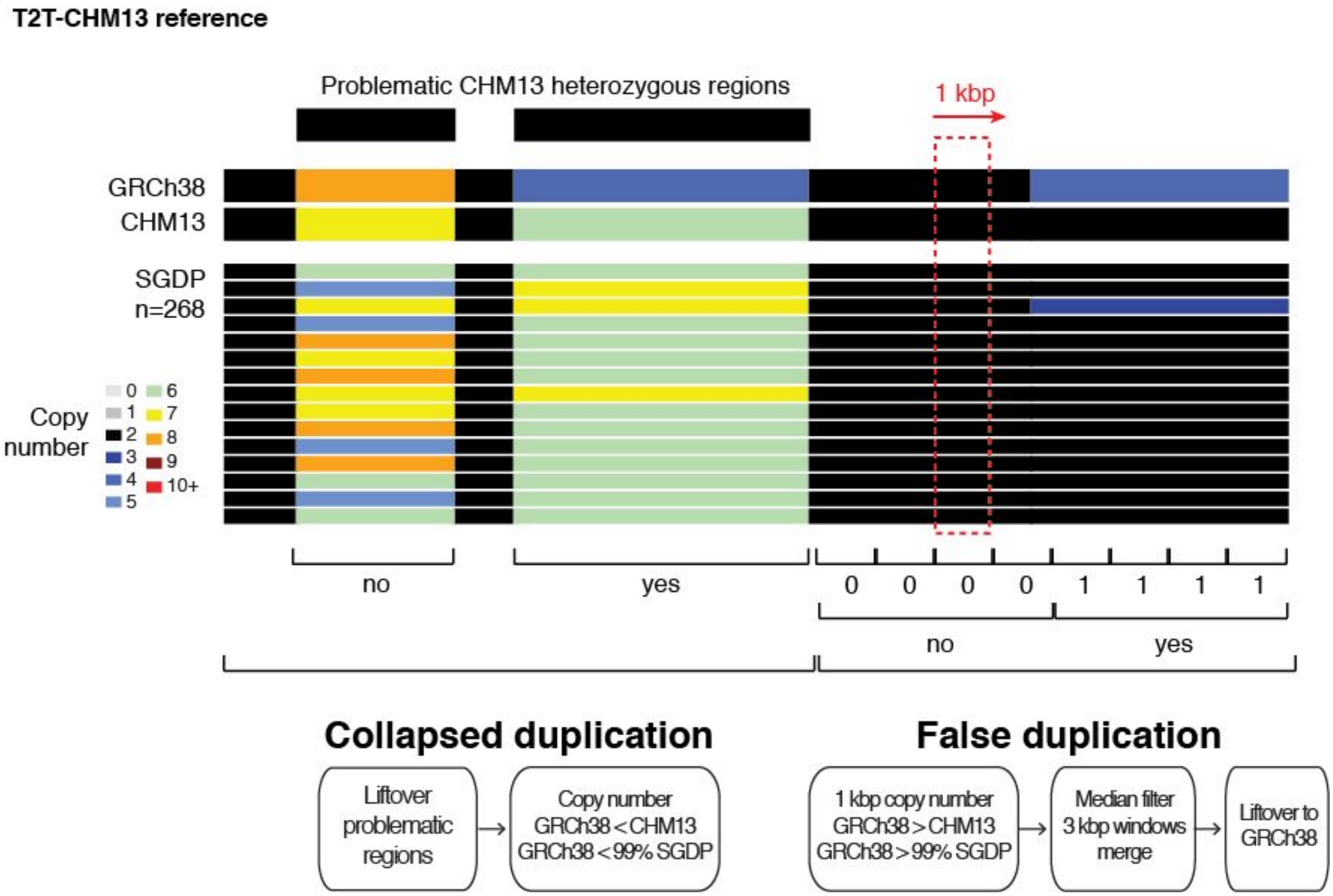


**Fig. S1.11. Strategy used to detect collapsed and false duplications in GRCh38**

WSSD read-depth copy-number estimates were obtained for ‘k-merized’ versions of GRCh38 and T2T-CHM13v1.0 references, and Illumina reads from 268 SGDP individuals in the CHM13 reference. To identify putative collapsed duplications, the median copy-number of *k*-merized GRCh38 was compared to population and *k*-merized CHM13v1.0 copy numbers for each CHM13 problematic heterozygous region identified in either GRCh38 (lifted coordinates) or CHM13v1.0. To identify false duplications, *k*-merized GRCh38 copy-number estimates were compared to population and *k*-merized CHM13v1.0 copy-numbers using 1-kbp windows genome-wide.

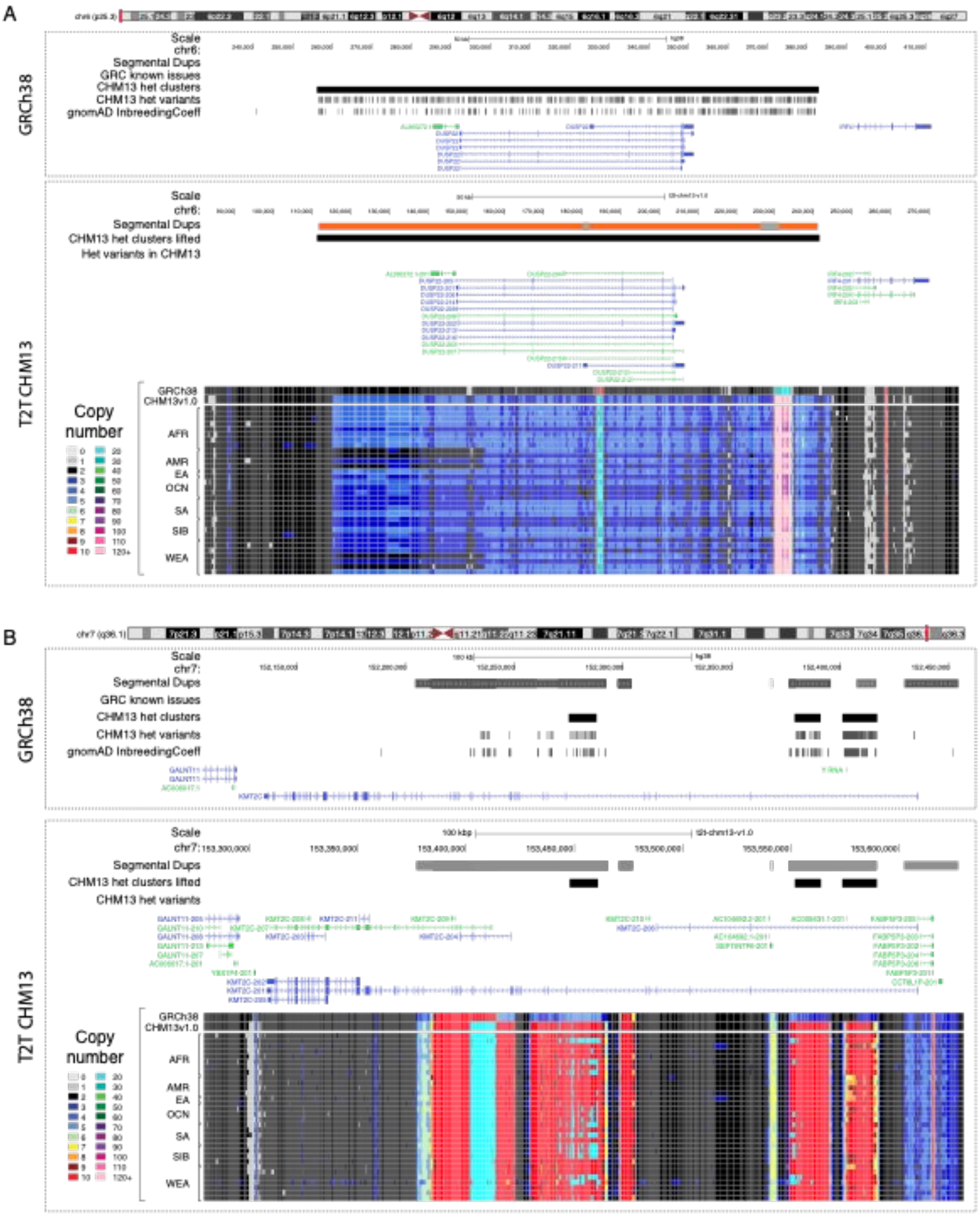


**Fig. S1.12. Examples of collapsed duplications impacting genes in GRCh38**

**A**. Collapsed duplication in GRCh38 corrected in CHM13 impacting *DUSP22.* **B**. Collapsed duplication in GRCh38 corrected in CHM13 impacting *KMT2C.* For both examples, CHM13 heterozygous variant clusters, CHM13 het variants, and gnomAD variants with the InbreedingCoeff flag are displayed on top, while read-depth copy-number estimates in CHM13 are shown at the bottom for 'k-merized' versions of GRCh38 and T2T-CHM13v1.0 references, and Illumina reads from a diverse subset (n=34) of SGDP individuals.

### Figure 2. Supplemental

T2T-CHM13 improves analysis of global genetic diversity based on 3,202 short-read samples from the 1KGP dataset

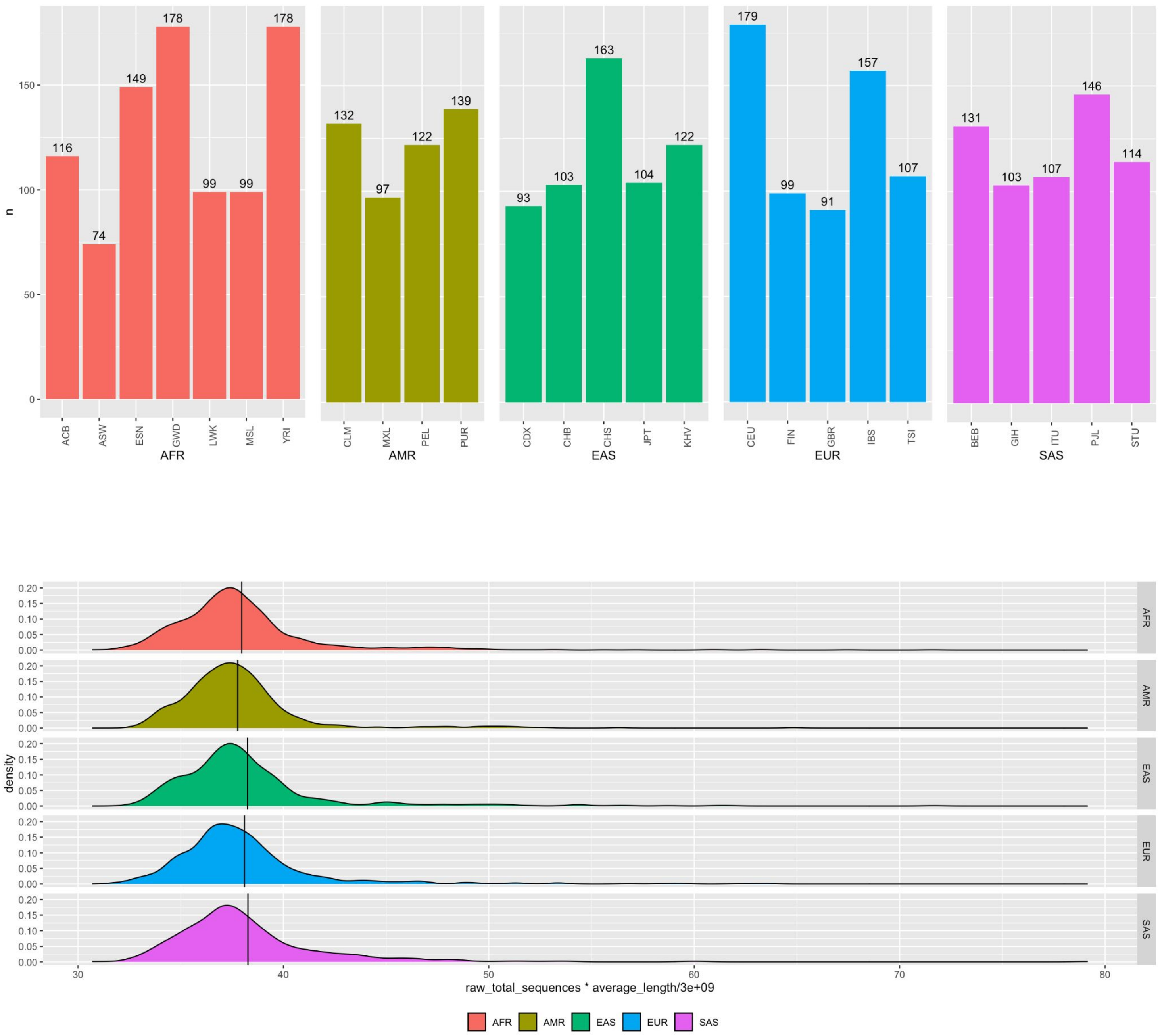


**Fig. S2.1. 1KGP Sample overview**

(top) Total number of samples per population, including children in trios, grouped by superpopulation. (bottom) Violin plot of the total amount of raw sequencing coverage available per sample, assuming a 3.0Gbp genome size. Black vertical line indicate the mean coverage per superpopulation.

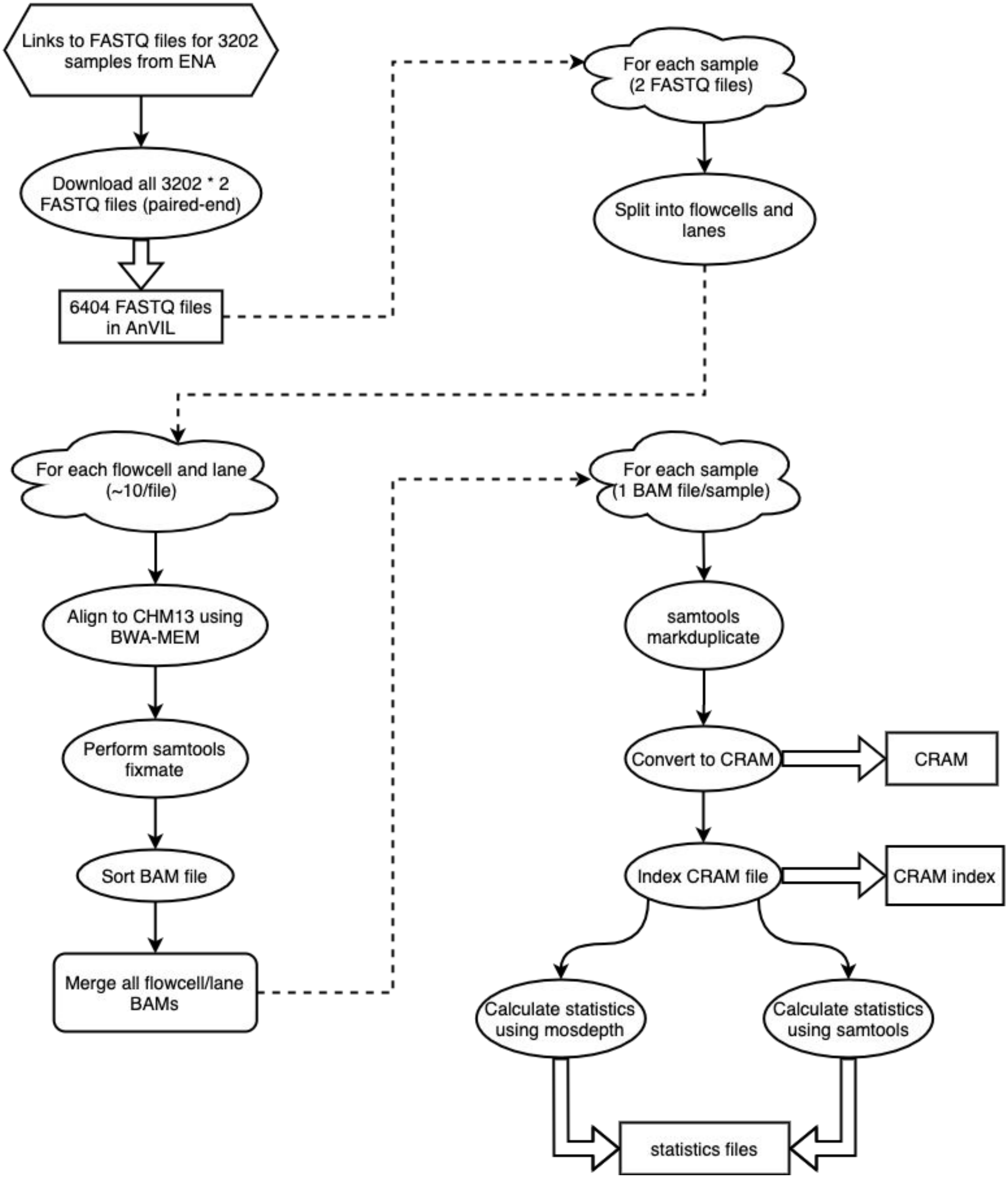


**Fig. S2.2. 1000 Genomes Project Alignment Pipeline**

Overview of the workflow for aligning 1000 Genomes Project samples to CHM13, as adapted from the NYGC's pipeline for variant calling 1KGP samples on GRCh38 data.

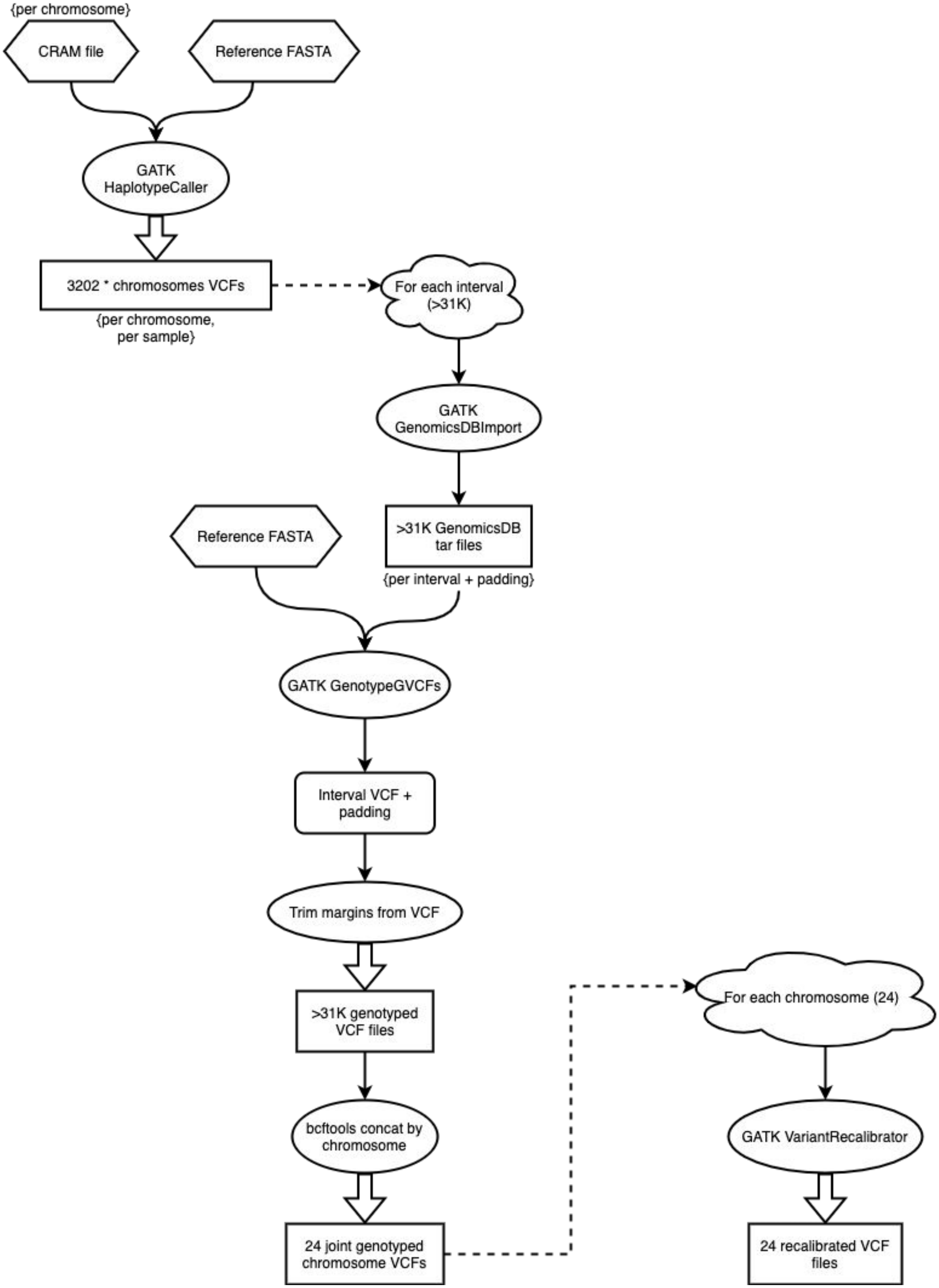


**Fig. S2.3. 1000 Genomes Project Variant Calling Pipeline**

Overview for the workflow for performing variant calling and joint genotyping on 1000 Genomes Project samples after alignment to CHM13, as adapted from the NYGC's pipeline for variant calling 1KGP samples on GRCh38 data.

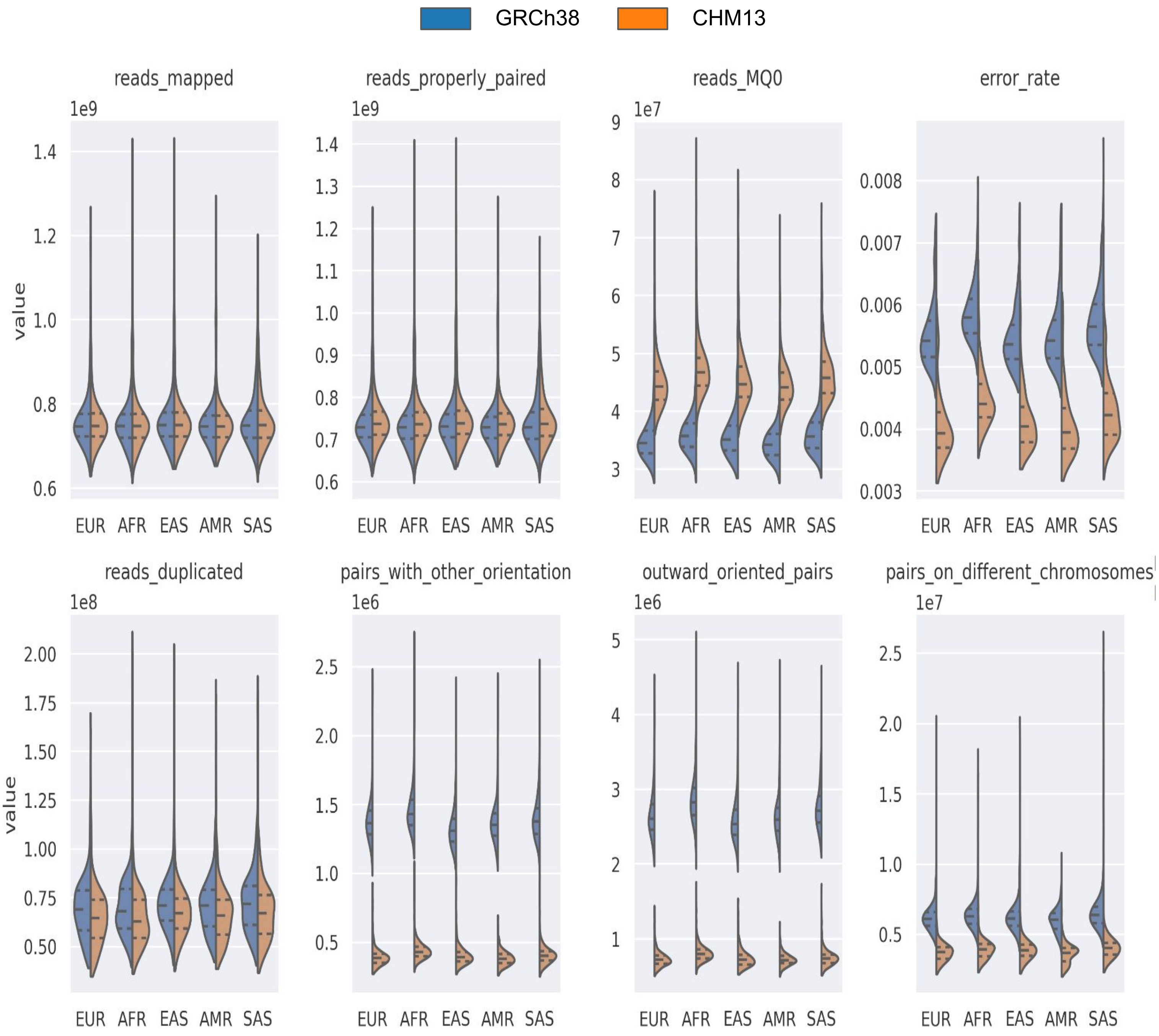


**Fig. S2.4. Short-read mapping statistics generated using samtools stats with GRCh38 and CHM13 as alignment target references**

Results are stratified by superpopulation codes as per 1KGP dataset. Distribution quartiles values are shown as dashed lines inside violin plots.

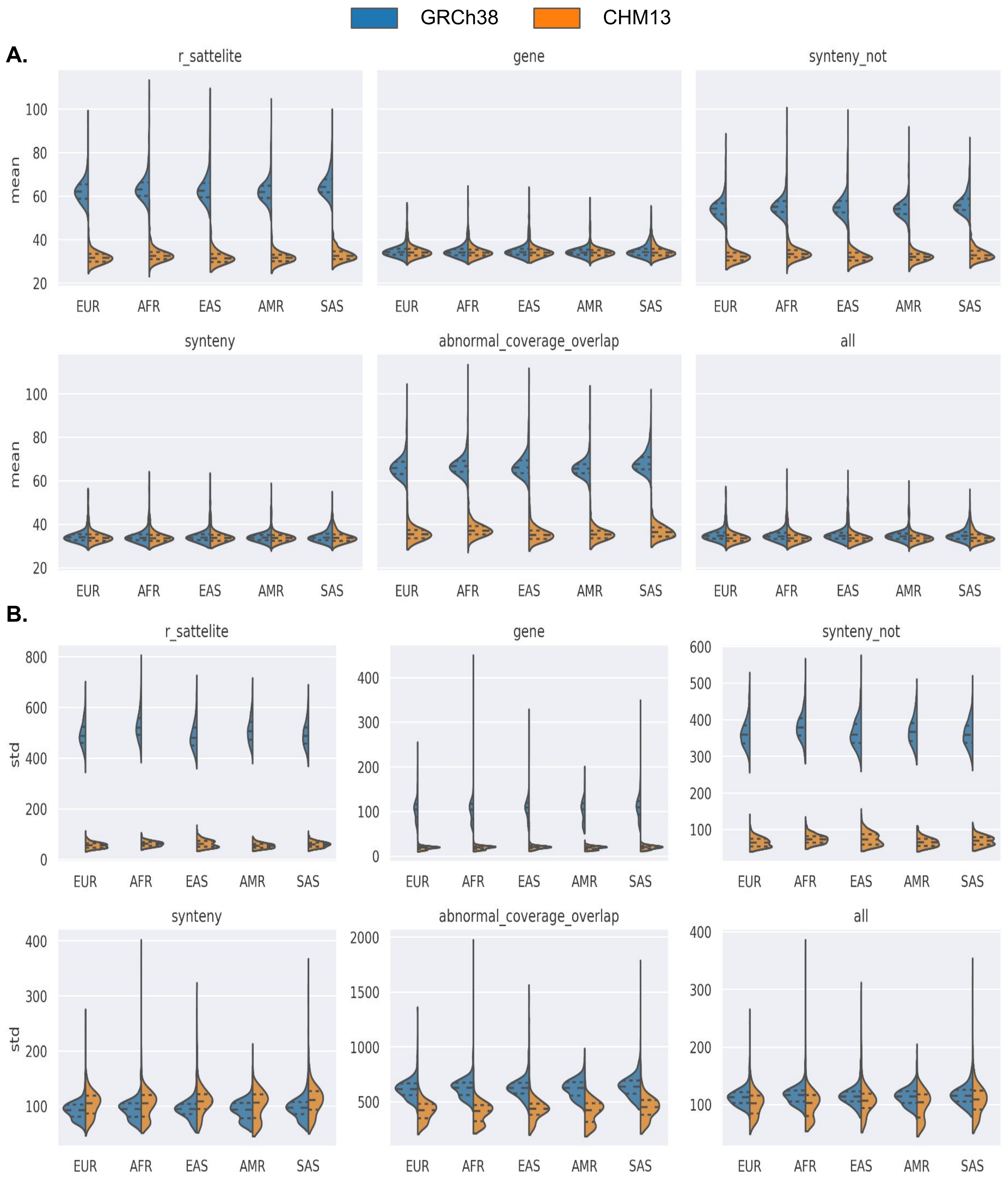


**Fig. S2.5. Short-read coverage statistics**

Distributions of per-sample mean (A) and standard deviation (B) values for read depth context-stratified 500bp-windowed intervals with alignment target references GRCh38 (blue) and CHM13 (orange) are grouped based on 1KGP superpopulation sample annotations. Distribution quartiles are shown as dashed lines.

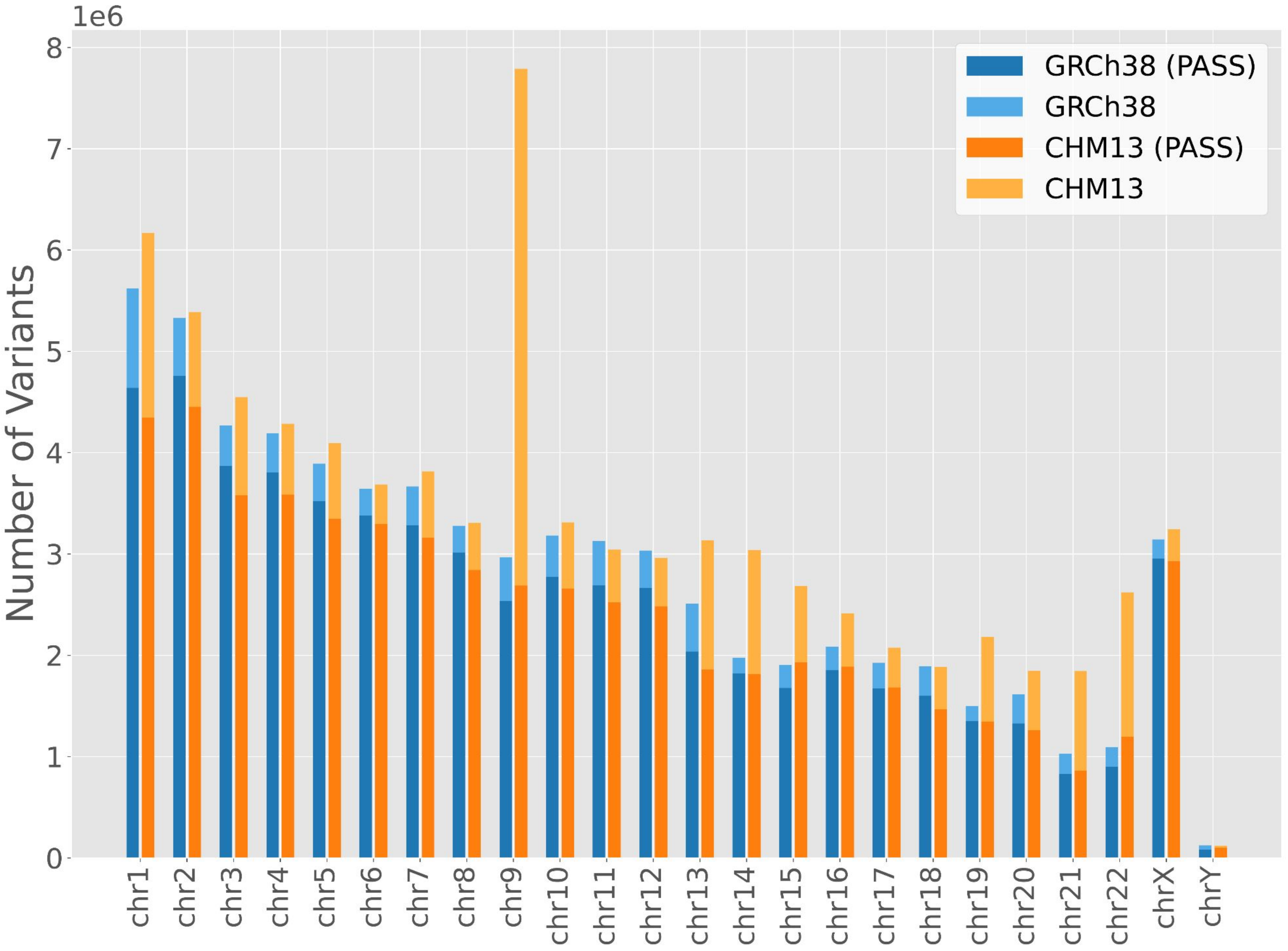


| | GRCh38 | CHM13 |
|---|---|---|
| All | 67,007,576 | 79,491,005 |
| Filtered by PASS | 59,051,733 | 57,315,786 |

**Fig. S2.6. Effect of PASS filtering**

The per-chromosome (top) and genome-wide (bottom) number of variants in the 1000 Genomes Project samples (allele frequency > 0) with respect to GRCh38 and CHM13, before and after filtering by GATK's "PASS" annotation.

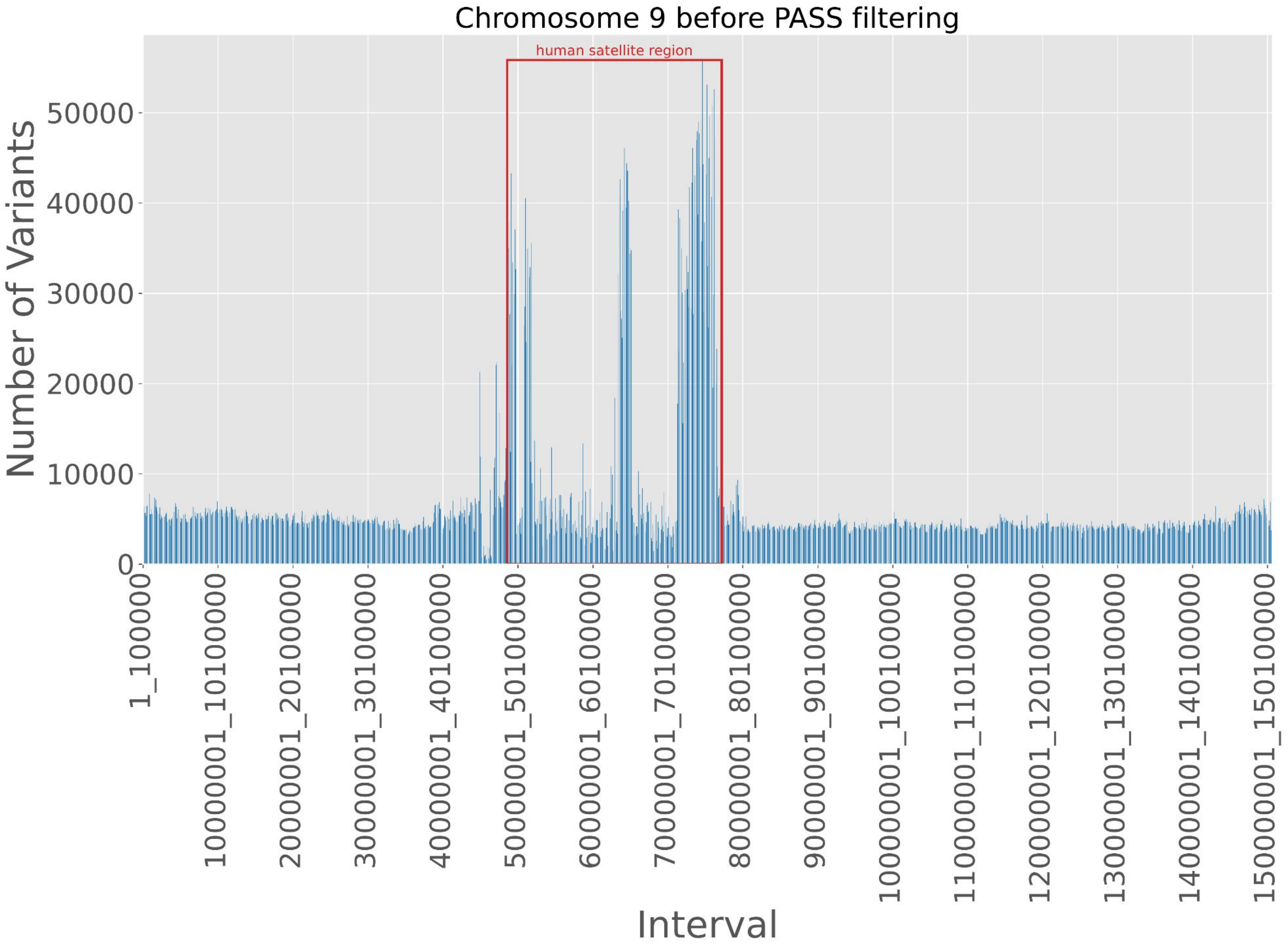


**Fig. S2.7. Complex regions affected by PASS filtering**

The number of variants in chromosome 9 when aligned to CHM13 before filtering by the "PASS" annotation, with a complex human satellite region annotated. Most of these variants are filtered out with the "PASS" annotation.

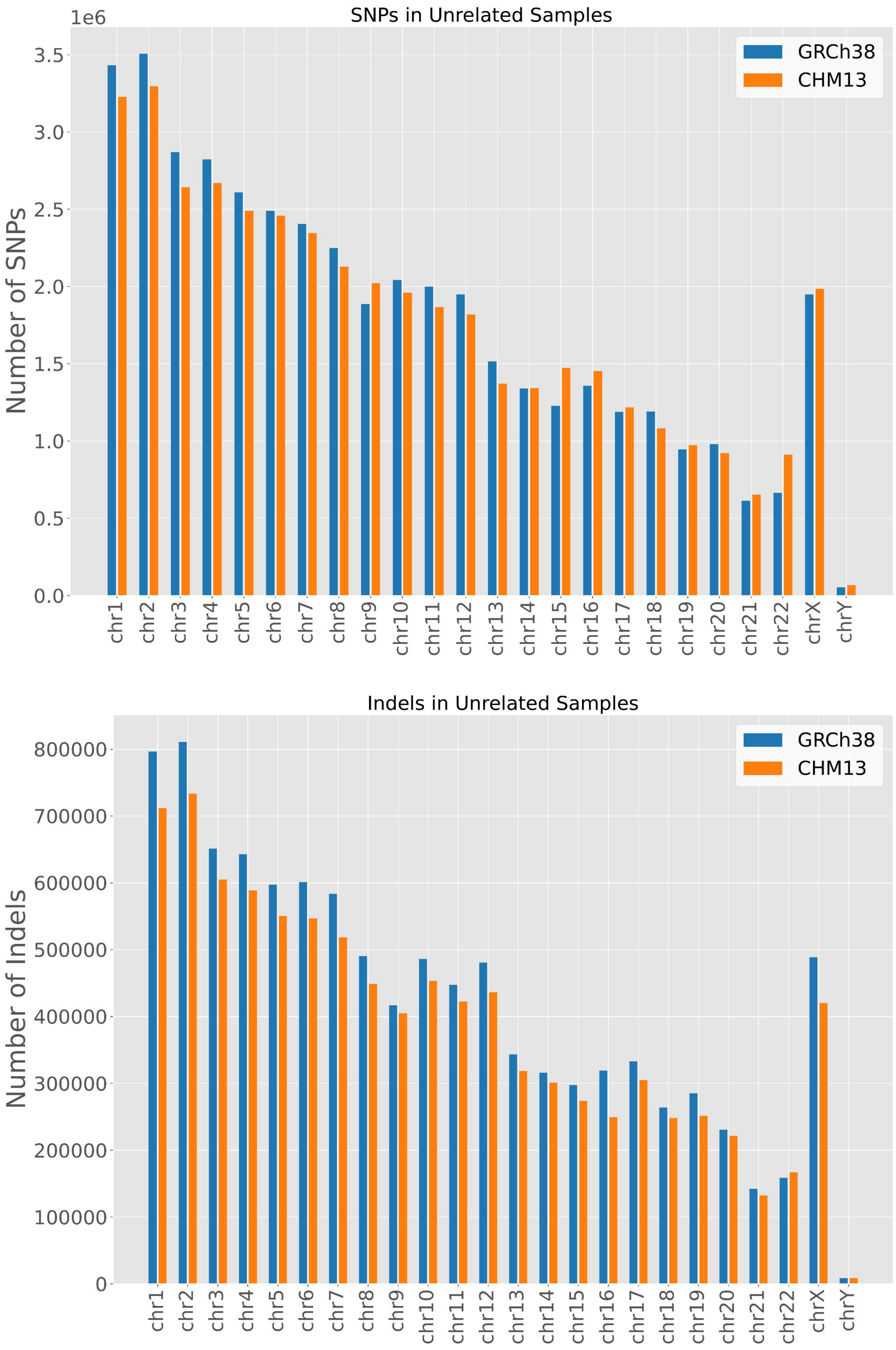


**Fig. S2.8. SNPs and indels in unrelated samples**

The number of SNPs (top) and indels (bottom) across all 1KGP samples (allele frequency > 0) when aligned to GRCh38 and CHM13.

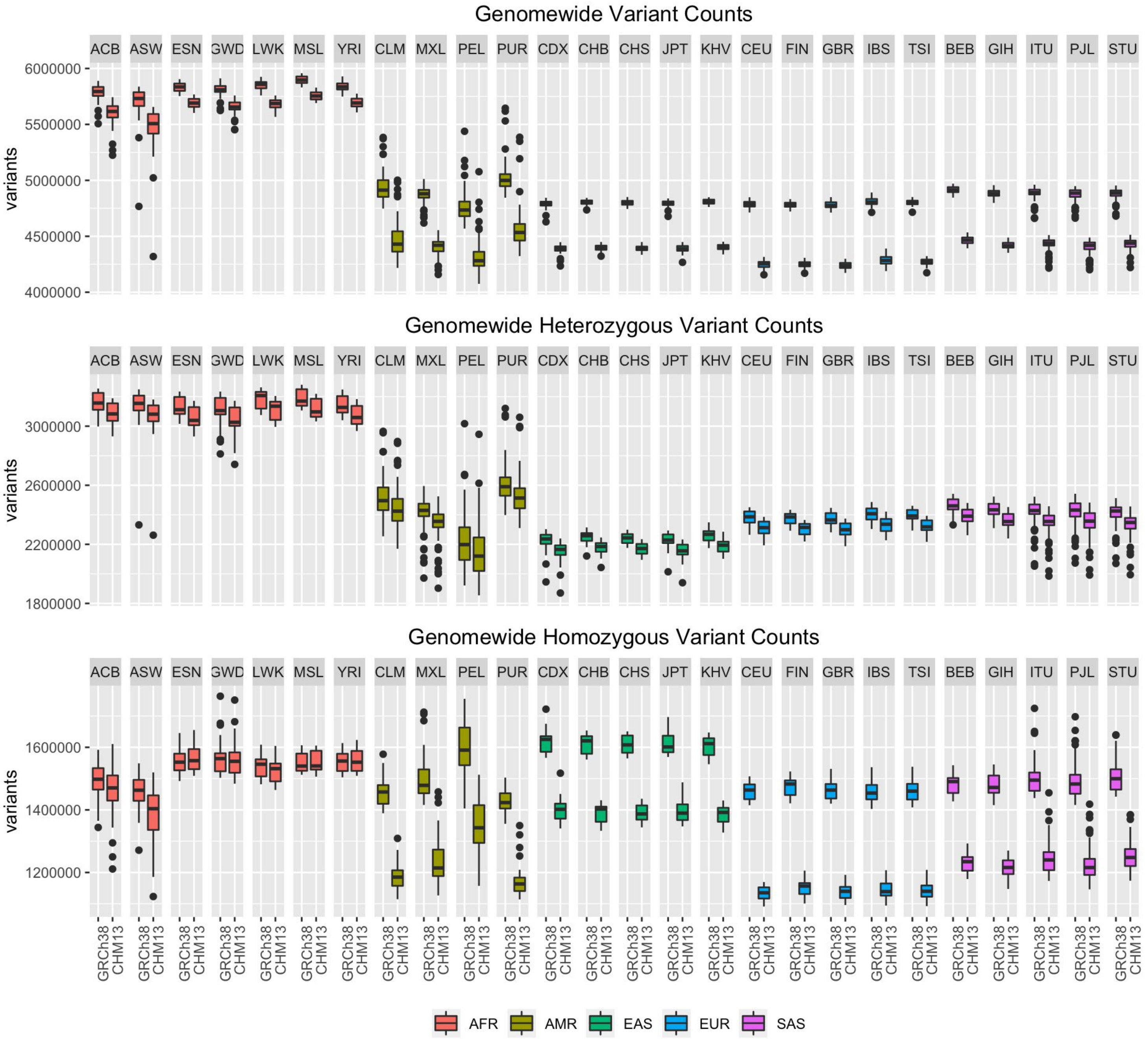


**Fig. S2.9.Population-specific variant counts**

Population-specific boxplots of the number of all variants (top), heterozygous variants (middle), and homozygous variants (bottom) per sample, as computed in Figure 2B.

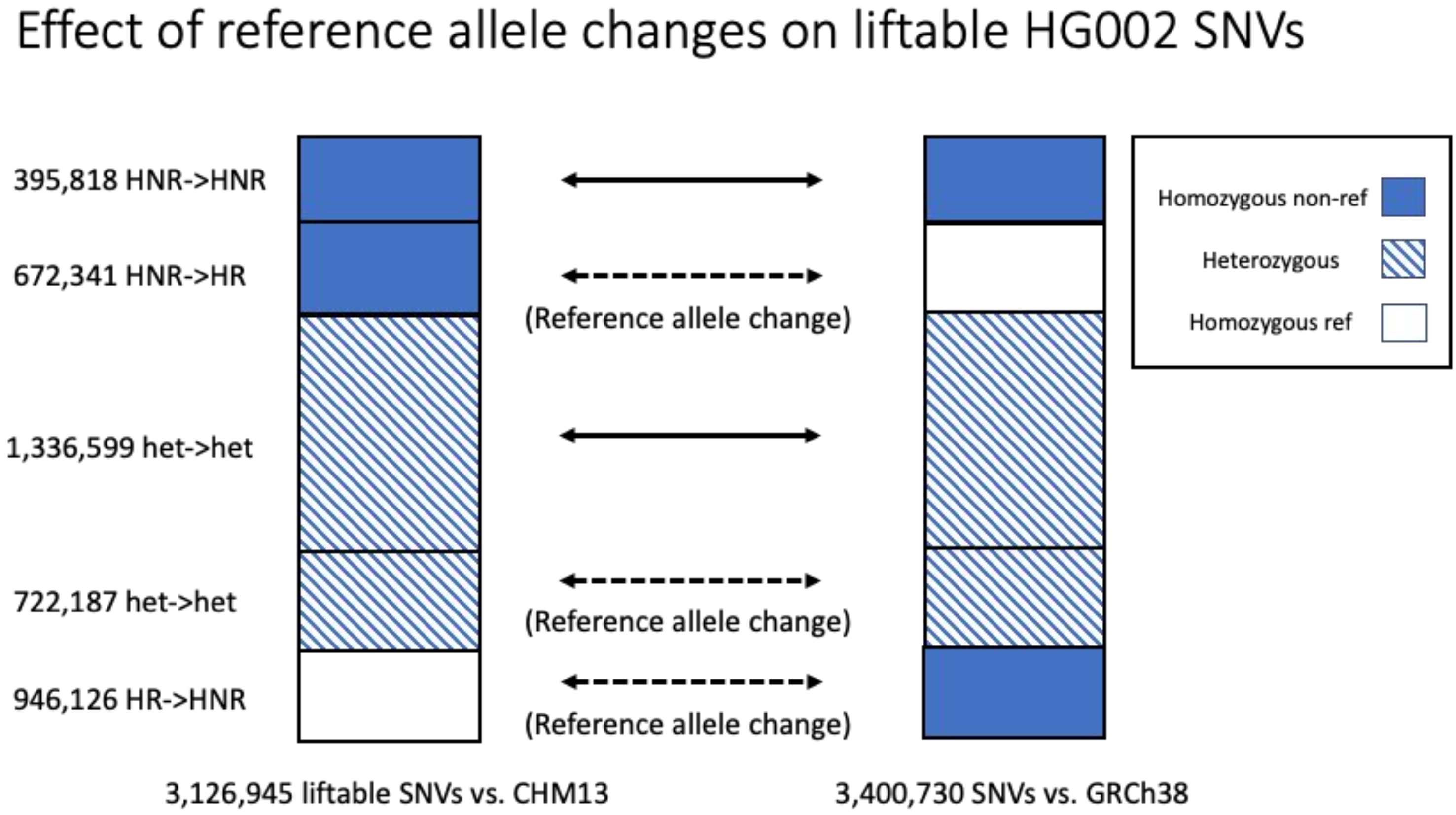


**Fig. S2.10. Effect of reference allele changes on SNV visibility in HG002.**

Many SNVs in the sample HG002 are not visible as variants with respect to either T2T-CHM13 or GRCh38 due to changes in the reference base such a homozygous variant call (HNR) changes to homozygous reference (HR). Many heterozygous SNVs (hets) also change due to reference base changes but are visible as variants on both references. Reported SNV counts are only among those with positions that lift successfully between references.

### Figure 3. Supplemental

T2T-CHM13 improves structural variant analysis of 17 diverse long-read samples

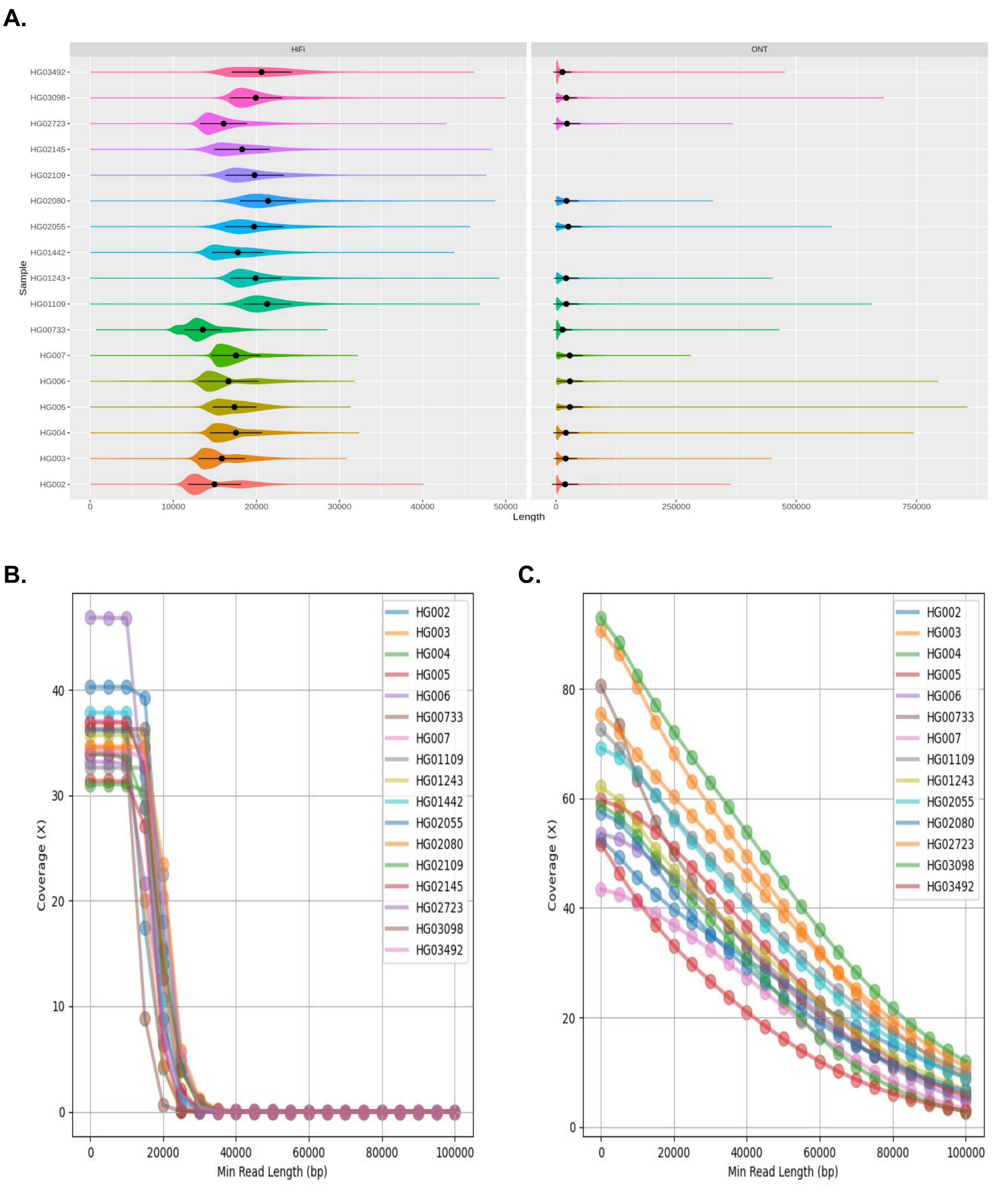


**Fig. S3.1. Read length distributions**

**A)** The distribution of HiFi read lengths in 17 samples and ONT read lengths in 14 of those samples. Points and error bars represent mean and standard deviation lengths in each sample. The mean of the per-sample mean lengths of HiFi reads is 18,130.2 bp. The mean of the per-sample mean lengths of ONT reads is 21,912.9 bp. **B)** HiFi coverage in each sample as a function of minimum read length. **c.)** ONT coverage in each sample as a function of minimum read length.

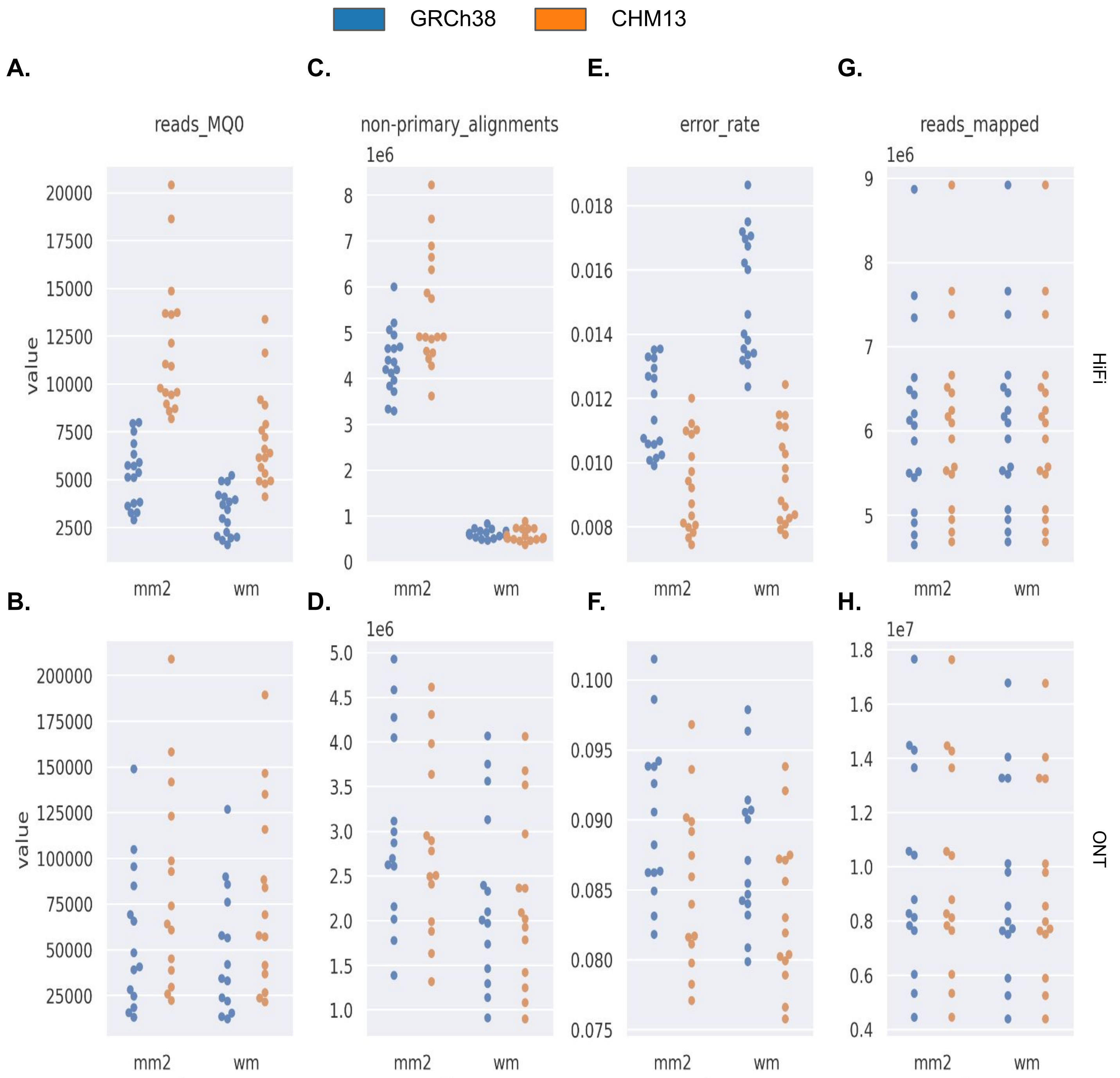


**Fig. S3.2. Long-Read mapping statistics generated with samtools stats**

**A)** The number of HiFi reads with 0 mapping quality across 17 samples in each reference.
**B)** The number of ONT reads with 0 mapping quality across 14 samples in each reference.
**C)** The number of HiFi reads with non-primary alignments across 17 samples in each reference.
**D)** The number of ONT reads with non-primary alignments across 14 samples in each reference.
**E)** The average error rate of HiFi reads across 17 samples in each reference.
**F)** The average error rate of ONT reads across 14 samples in each reference.
**G)** The number of HiFi reads mapped across 17 samples in each reference.
**H)** The number of ONT reads mapped across 14 samples in each reference.

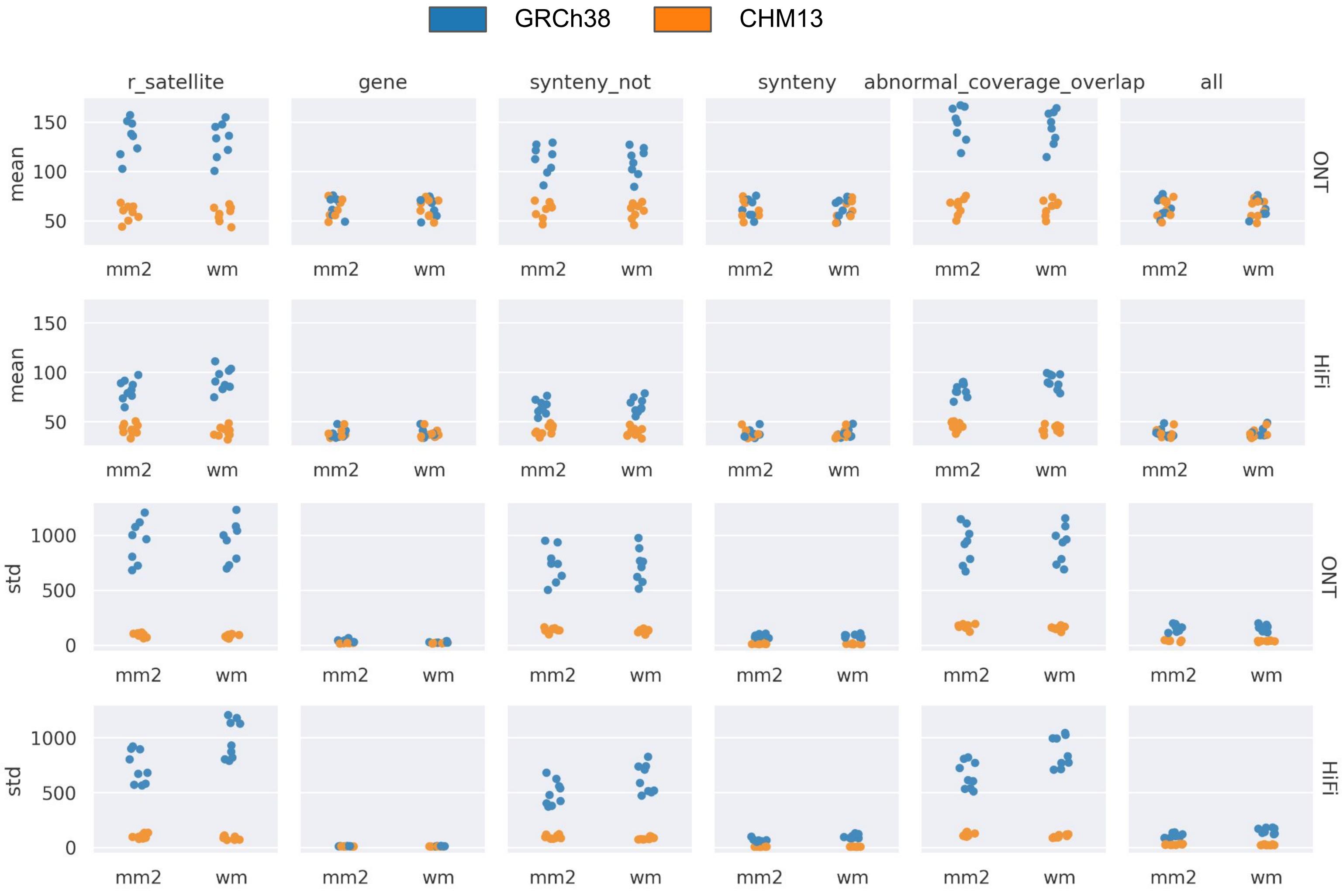


**Fig. S3.3. Long-read abnormal coverage statistics**
The mean and standard deviation of coverage among 500bp bins in CHM13 and GRCh38 when using different combinations of aligners and sequencing technologies. The overall counts are displayed, as well as bins which overlap satellite repeats, genes, non-syntenic regions with respect to the other reference, syntenic regions, and bins with abnormal levels of coverage.

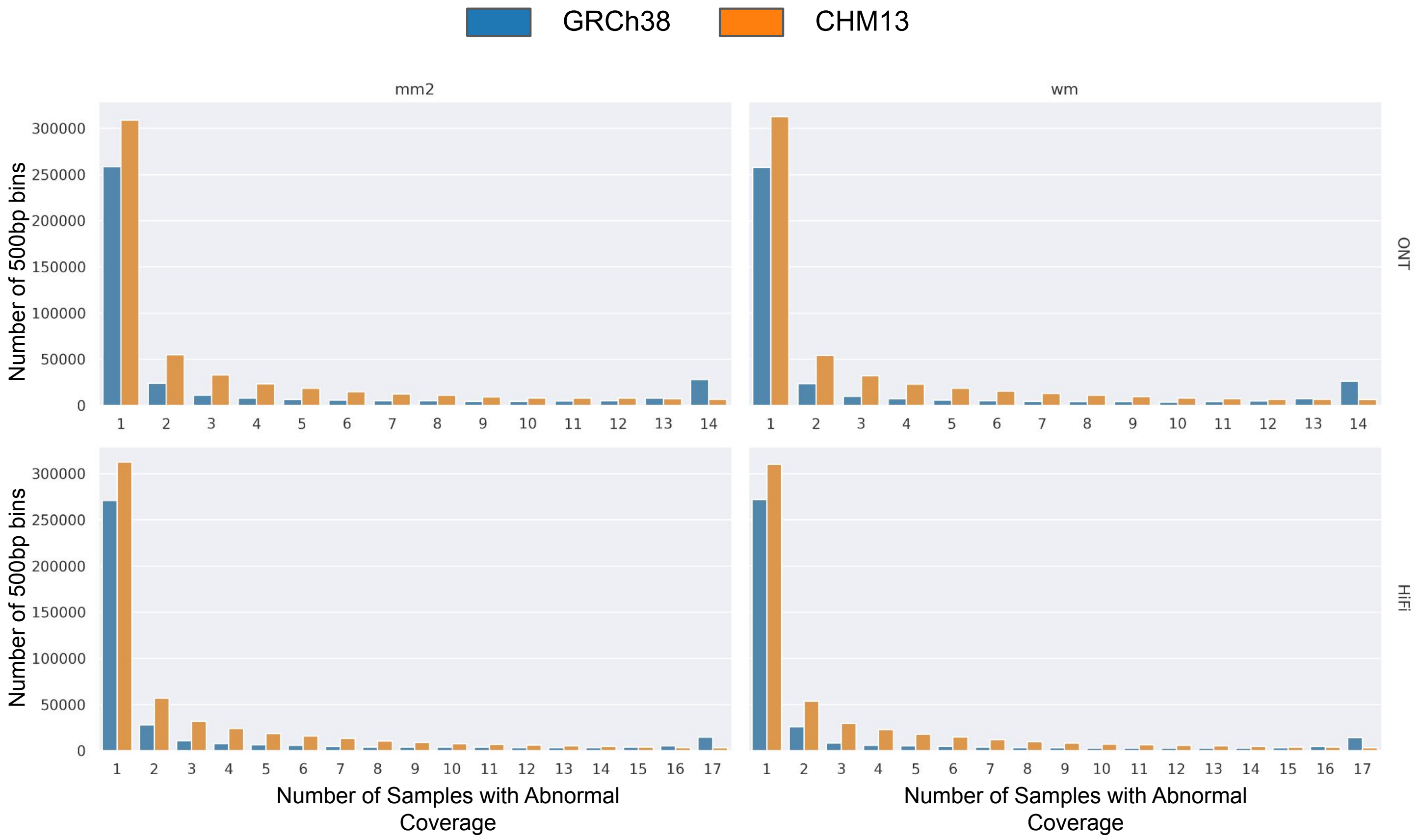


**Fig. S3.4. Long-read abnormal coverage frequency**
A histogram of the number of 500bp bins with different frequencies of abnormal coverage among the samples studied. Here abnormal coverage is defined as outside the range [Median - 1.5 (Median - Q1), Median + 1.5 (Q3 - Median)] among all bins in the same reference.

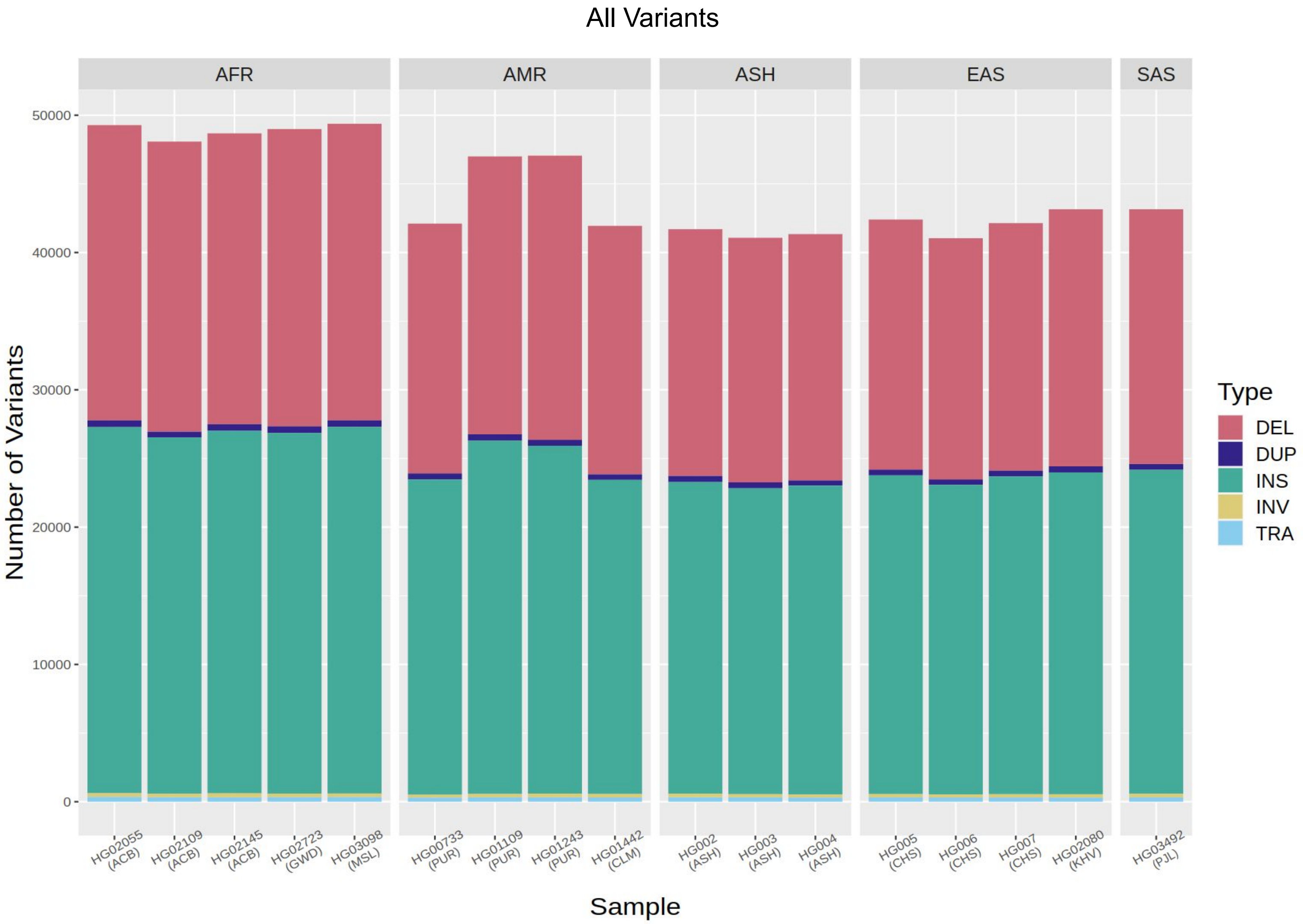


**Fig. S3.5. Per-sample SV counts**

The counts of HiFi-derived SV calls in each of the 17 samples studied. Counts are post-merging, so include variants which were called with high-confidence in that sample, as well as variants which were called with low-confidence in that sample but which were merged with high-confidence calls in other samples.

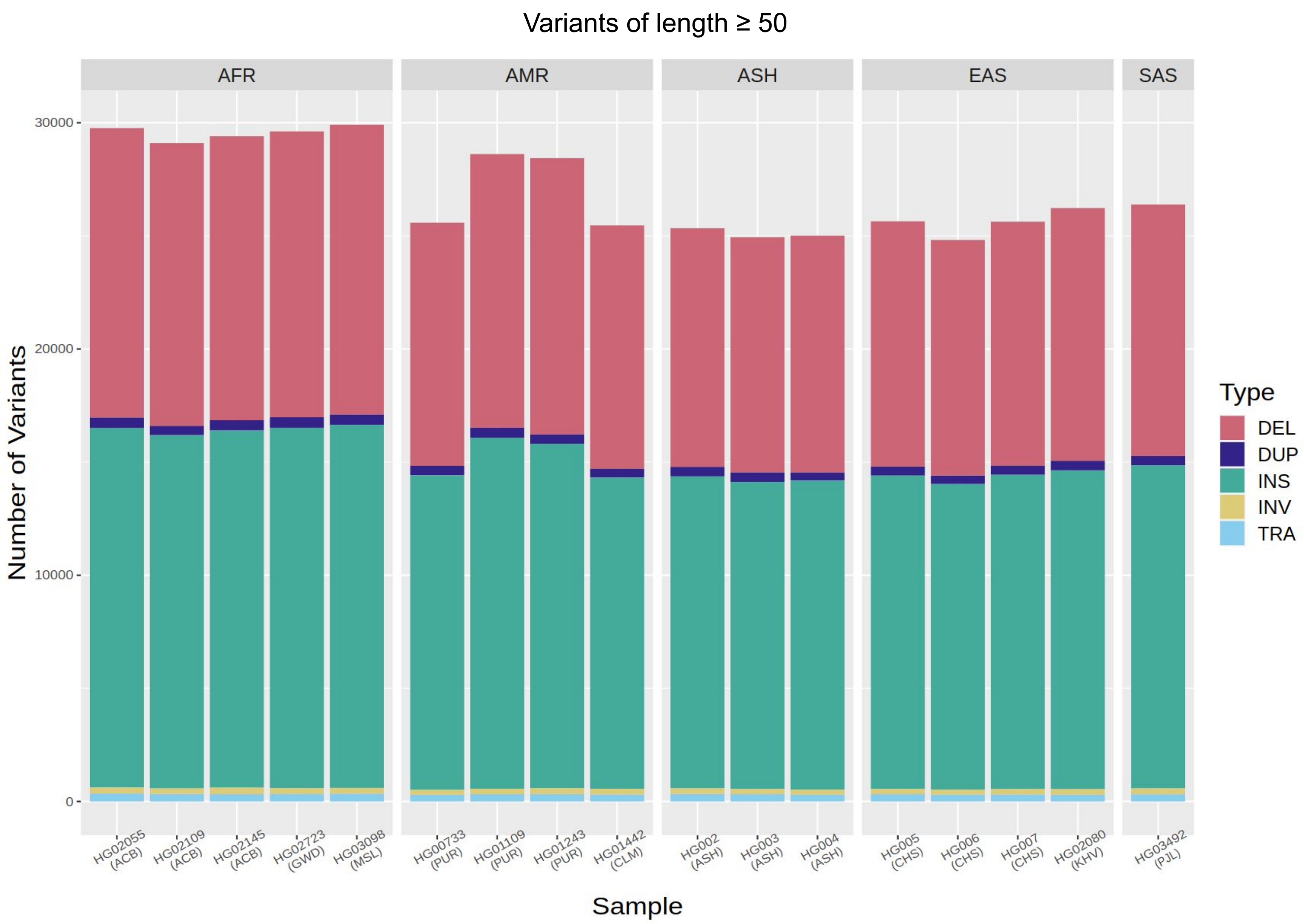


**Fig. S3.6. Per-sample variant counts (length ≥ 50)**
The counts of HiFi-derived SV calls with length at least 50, plus translocations, in each of the 17 samples studied. Counts are post-merging, so include variants which were called with high-confidence in that sample, as well as variants which were called with low-confidence in that sample but which were merged with high-confidence calls in other samples.

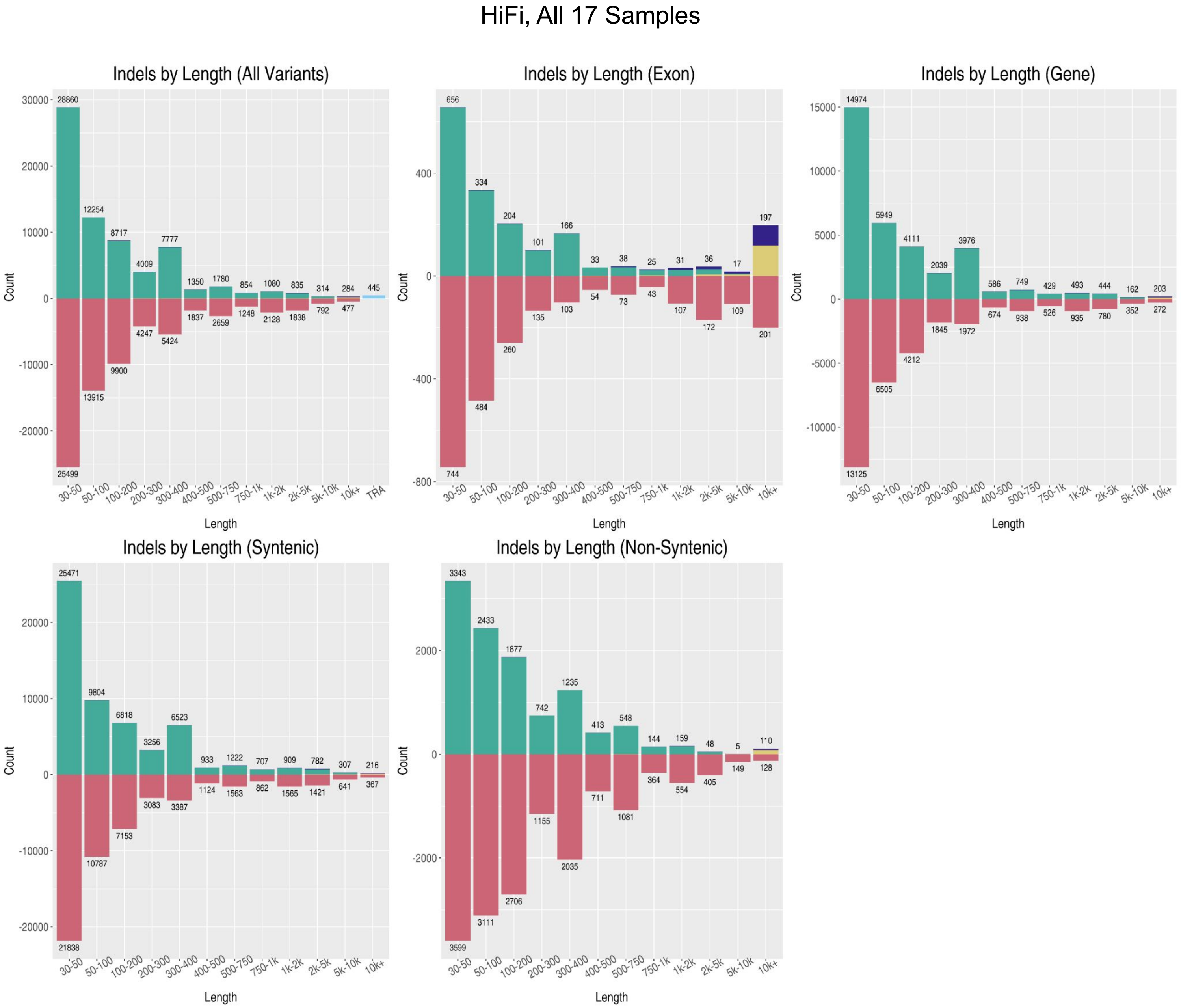


**Fig. S3.7. CHM13 variants across genomic contexts**

The lengths and types of SVs in CHM13 overlapping genes, exons, regions syntenic to GRCh38, and regions non-syntenic to GRCh38 called from HiFi data in our cohort of 17 samples.

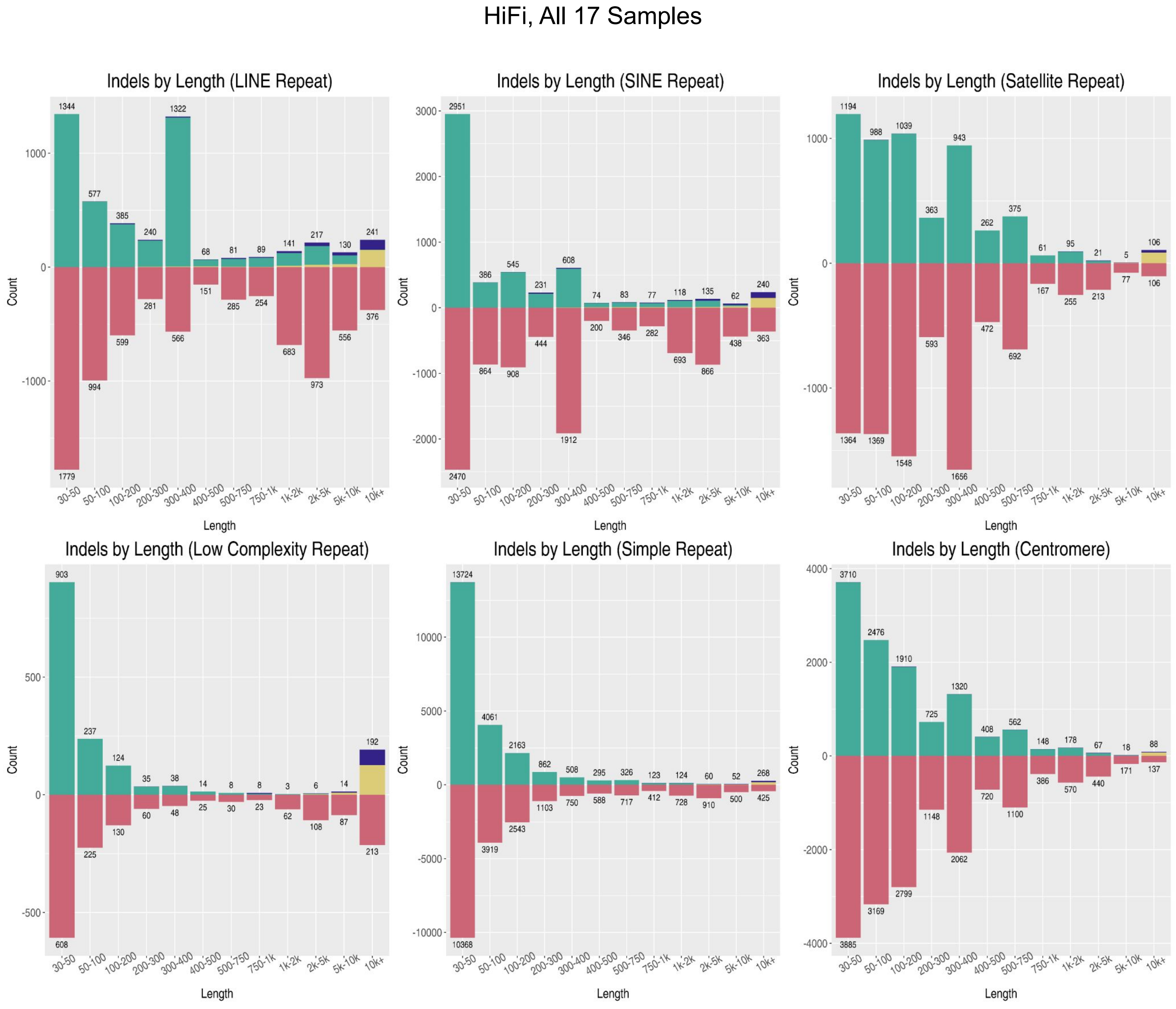


**Fig. S3.8. CHM13 variants in repeats**

The lengths and types of SVs in CHM13 overlapping various repeat classes called from HiFi data in our cohort of 17 samples.

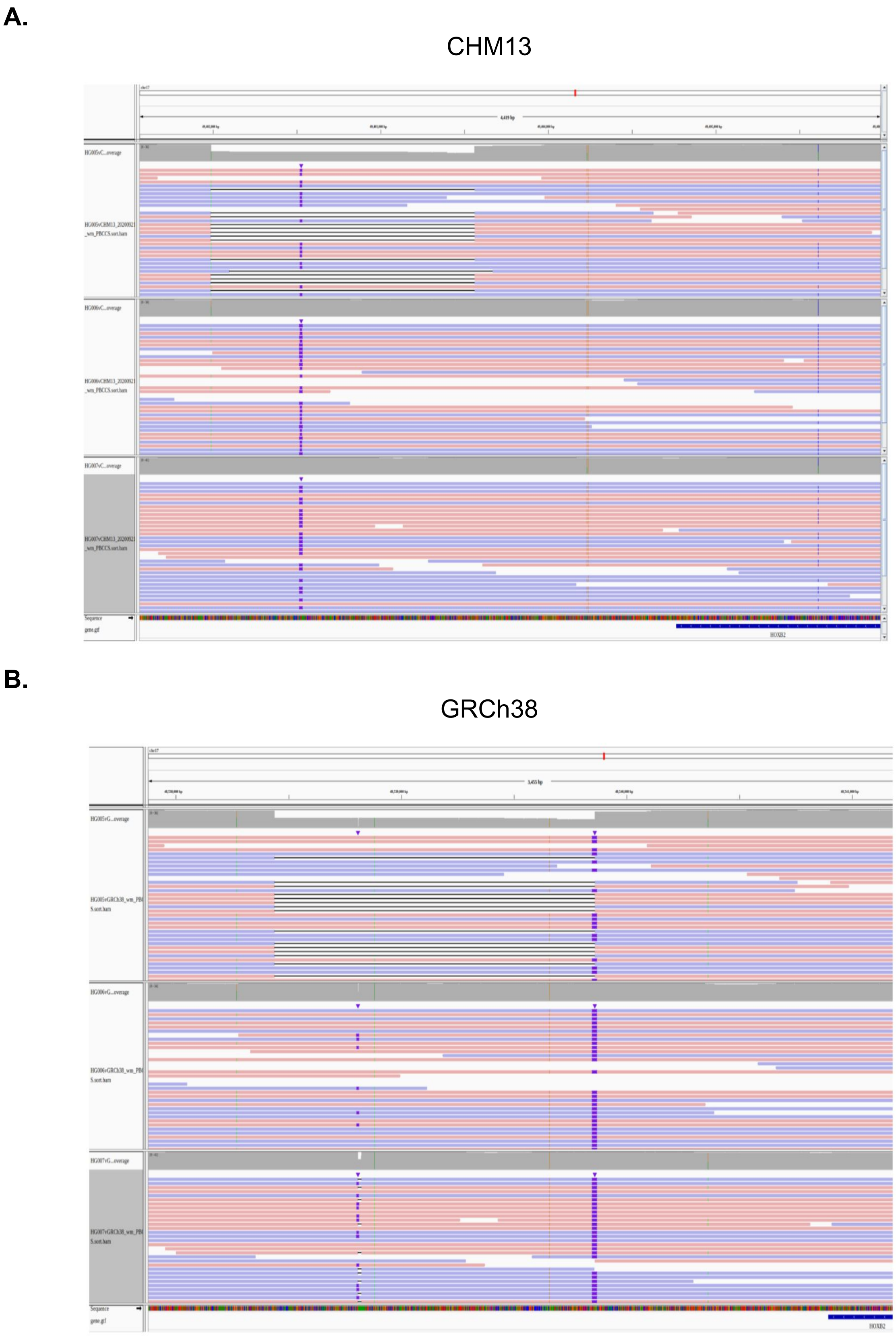


**Fig. S3.9. Potential de novo SV in HG005**

A putative de novo 1,571 bp deletion in HG005 at chr17:49,401,990 in CHM13. **a.)** The alignments of the reads of HG005 (child), HG006 (parent, 46XY), and HG007 (parent, 46XX) to CHM13 near the SV call, indicating the SV's presence in HG005 and absence in the parents. **b.)** The alignments of the same reads to GRCh38

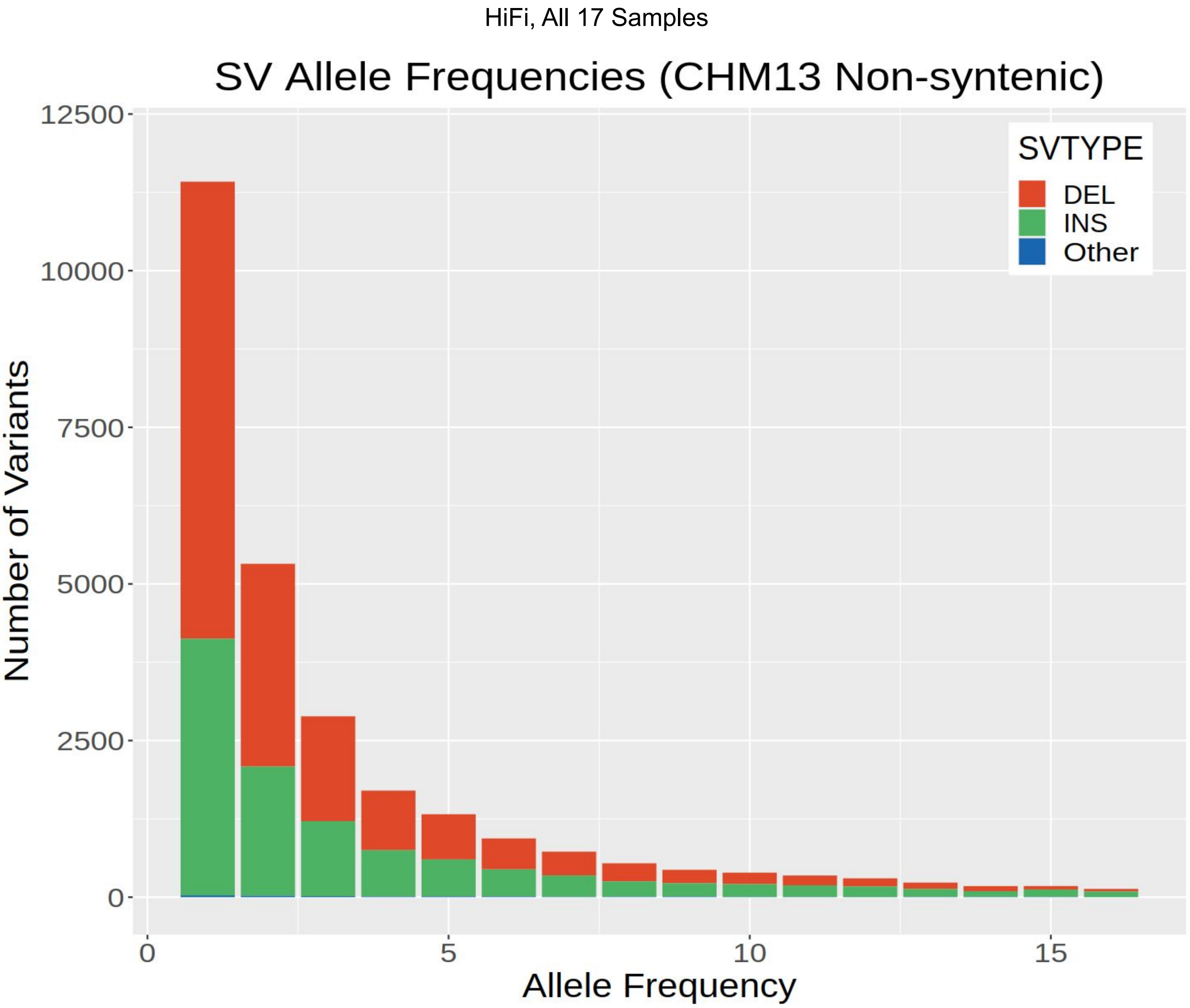


**Fig. S3.10. Allele frequencies of SVs in non-syntenic regions**
The allele frequency of structural variants overlapping non-syntenic regions in CHM13, called from HiFi data in our cohort of 17 samples.

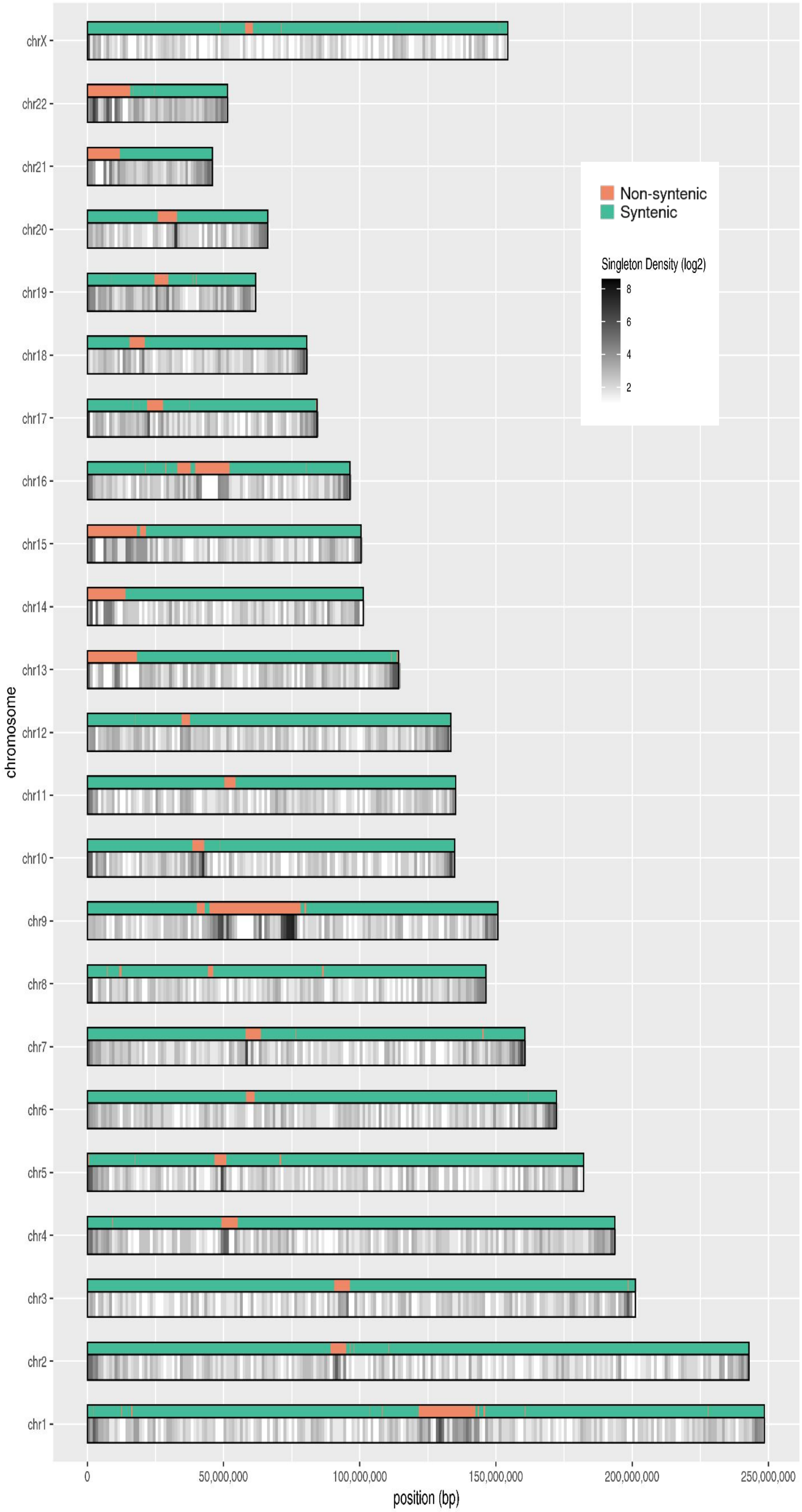


**Fig. S3.11. Singleton SV density across CHM13**

The density of singletons SVs (i.e. present in only a single sample), across 1 Mbp bins of CHM13.

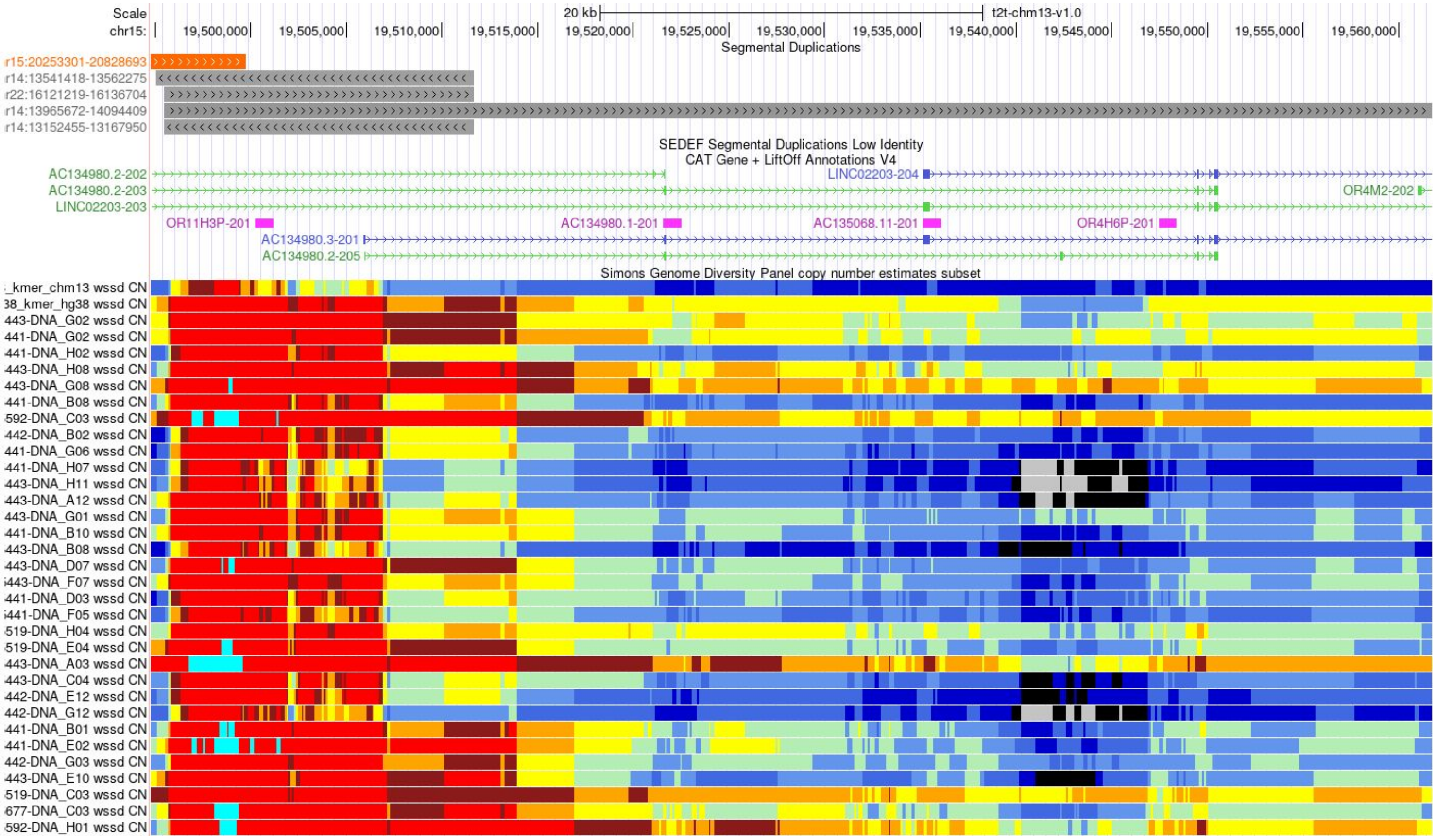


**Fig. S3.12. Copy number variation in AC134980.2**

Copy number variation of the region surrounding an exon of AC134980.2 (shown in Figure 4g and Figure 4h) among CHM13 and GRCh38, as well as samples from the Simons Diversity Panel.

**A.**

CHM13

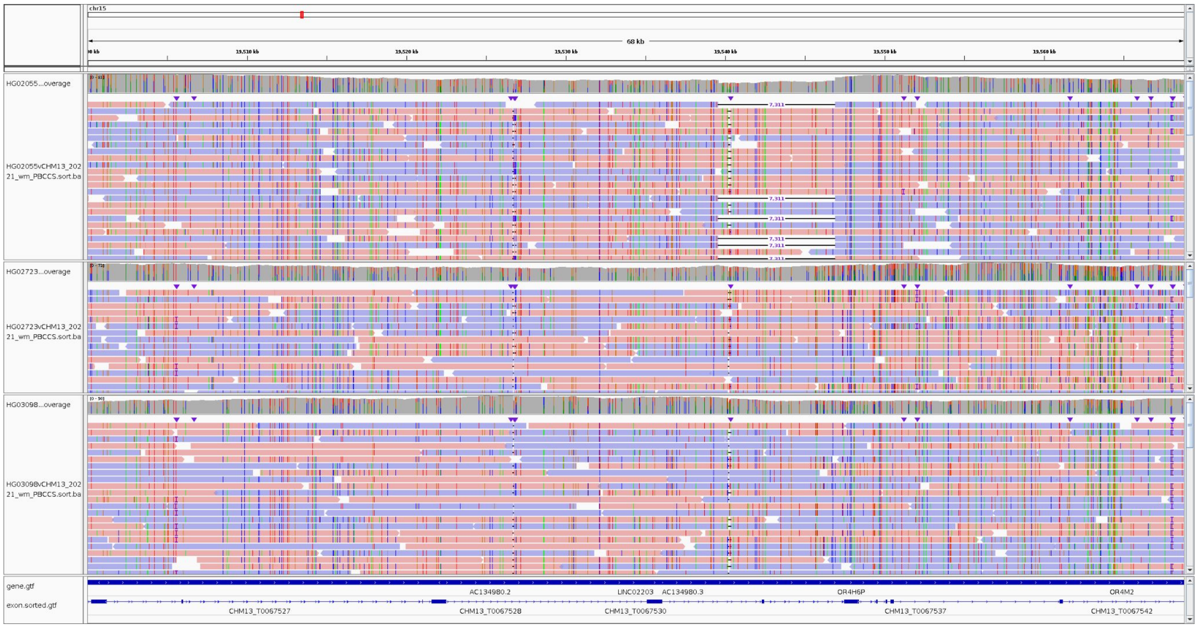


**B.**

GRCh38

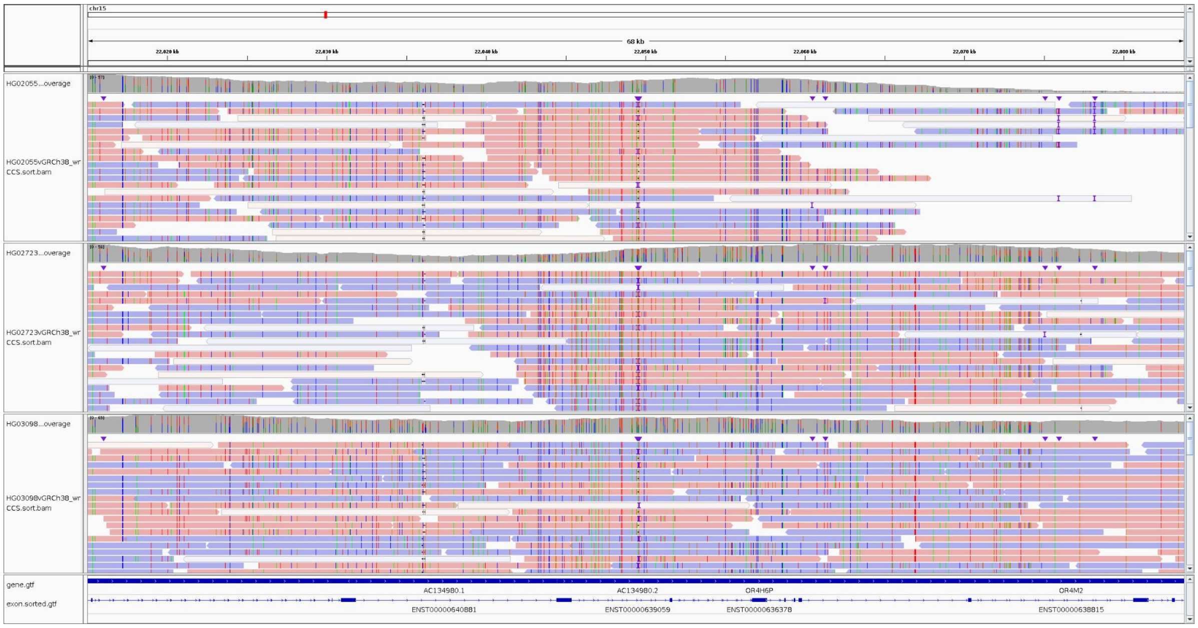


**Fig. S3.13. Alignments to AC134980.2 among samples of African ancestry**

Alignments of three samples of African ancestry - HG02055, HG02723, and HG03098 respectively - to the region around an exon of AC134980.2 (shown in Figure 4g and Figure 4h) to **a.)** CHM13 and **b.)** GRCh38.

#### SV Counts Overview

| | HG002 Assembly | | | | | | |
|---|---|---|---|---|---|---|---|
| | SV hg38 | SV Chm13 | SV unique to chm13 overlapping non-synteny | SV unique to chm13 overlapping non-synteny and at seg dups but not at centromere | SV found in both ref (hg38, chm13) | SV unique to hg38 | SV unique to chm13 |
| Deletion | 1199 | 1379 | 147 | 37 | 501 | 698 | 878 |
| Insertion | 2771 | 1431 | 129 | 26 | 588 | 2182 | 843 |
| Duplication | 38 | 35 | 19 | 6 | 14 | 23 | 21 |
| Inversion breakpoints | 22 | 20 | 11 | 0 | 6 | 16 | 14 |

(SV counts after confidence filtering and clustering)

**Fig. S3.14. SV counts from Bionano**

Comparison of SV calls >500 bp in HG002 from Bionano on GRCh38 (hg38) vs. T2T-CHM13, after filtering for quality and clustering equivalent calls. Balance of insertions and deletions is substantially improved. We also found number of SVs uniquely called on T2T-CHM13 in non-syntenic regions, and in non-syntenic regions that are in segmental duplications but not in centromere-satellite regions.

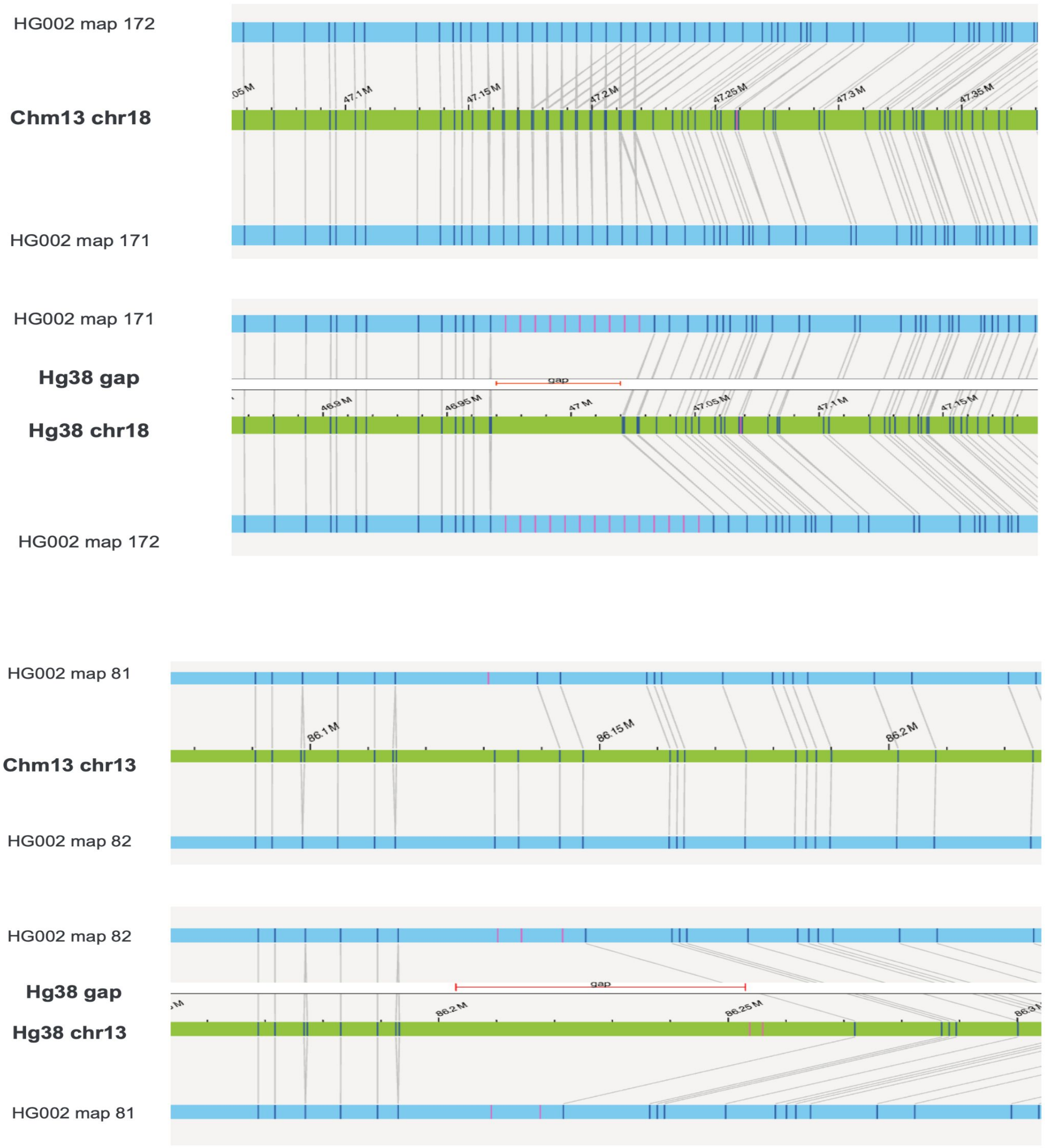


**Fig. S3.15. Improved Bionano alignment in CHM13 by closing gaps in GRCh38**

Two examples showing improved resolution of SV calls that were gaps on chr18 and chr13 in GRCh38 but are fully sequenced in CHM13. Lines connect sequence markers in HG002 maternal and paternal assemblies to *in silico* digested references GRCh38 (hg38) or T2T-CHM13 (Chm13). Gaps in GRCh38 are identified by red bars.

### Figure 4. Supplemental

Novel variants and evolutionary signatures within newly accessible regions of the genome

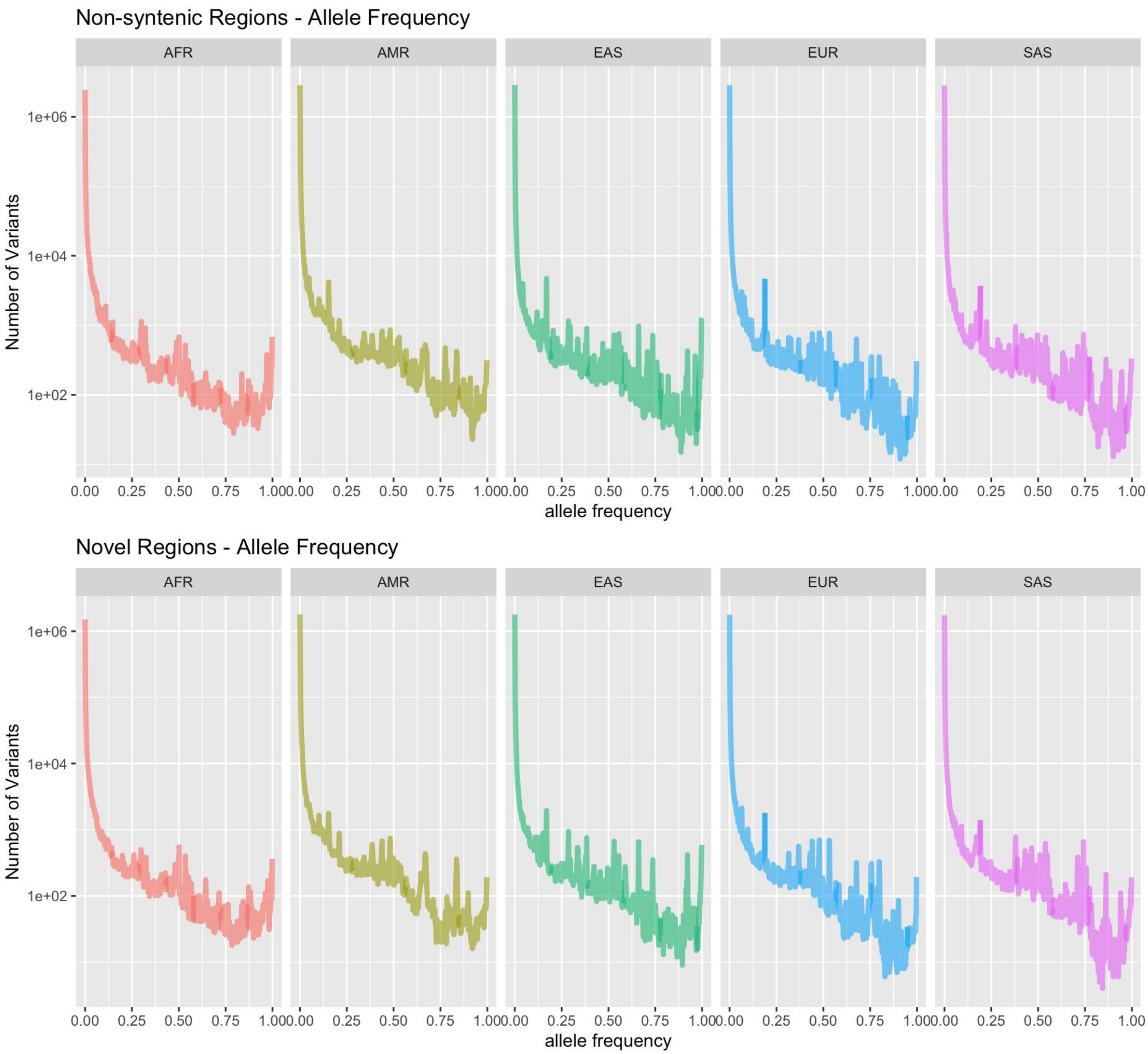


**Fig. S4.1. Variant allele frequencies within non-syntenic and novel regions**

SNV allele frequencies within non-syntenic (top) and novel (bottom) regions across the 1KGP samples for each of the 5 Superpopulations. Values are computed using unrelated (founder) samples as in Figure 2e although only variants within the autosomes are counted here.

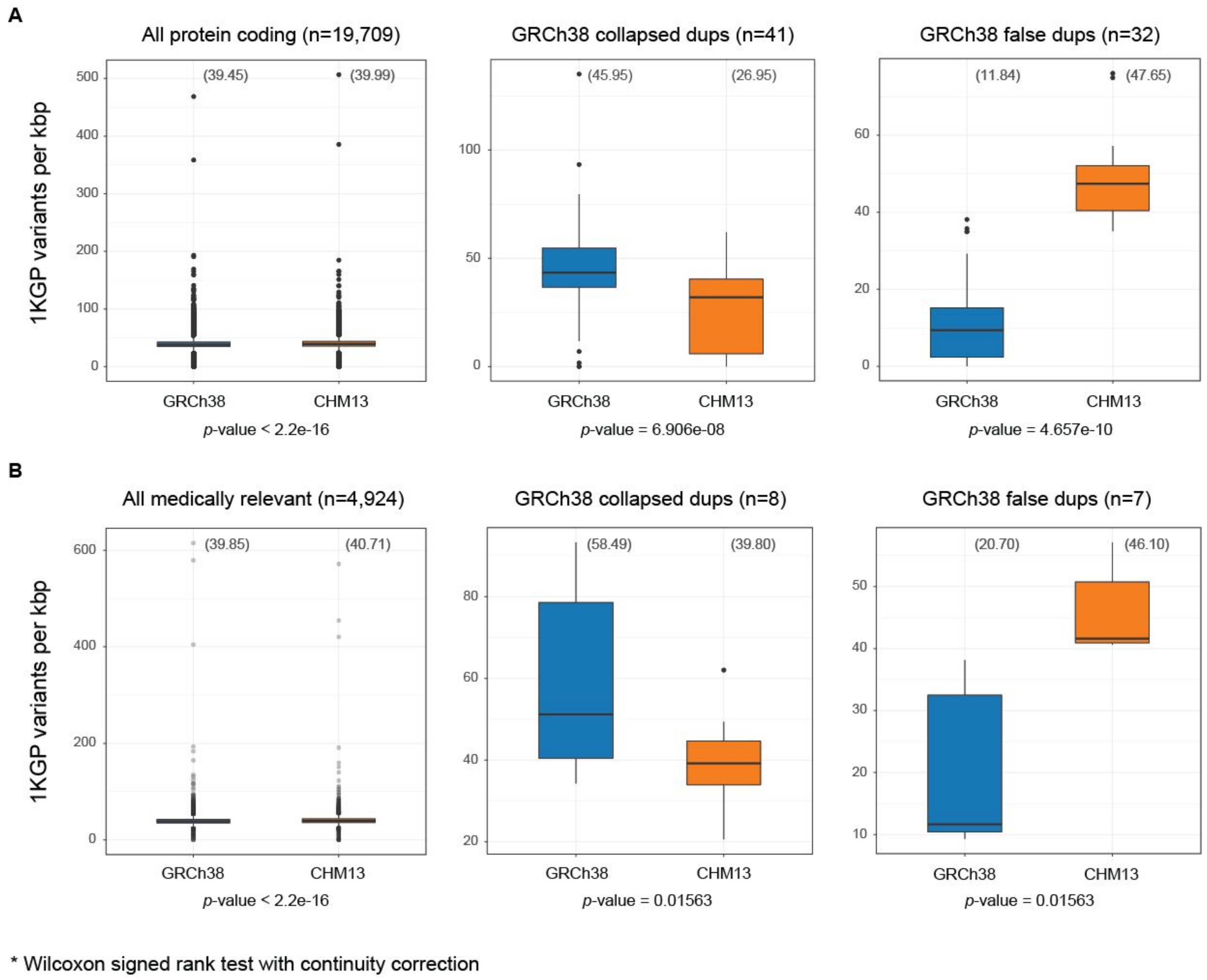


**Fig. S4.2. Comparison of 1 KGP variant densities within protein-coding genes between GRCh38 and T2T-CHM13.**

Variant densities are depicted as box plots across protein-coding genes in GRCh38 (blue) with successful lift over to T2T-CHM13 (orange) considering **(A)** all genes and **(B)** medically-relevant genes. Within these gene sets, variant densities were also determined for genes falling within GRCh38 collapsed duplications (dups) and false dups. Mean variant densities per reference genome is displayed in parentheses. *P-values* were calculated using a Wilcoxon signed-rank test with continuity correction.

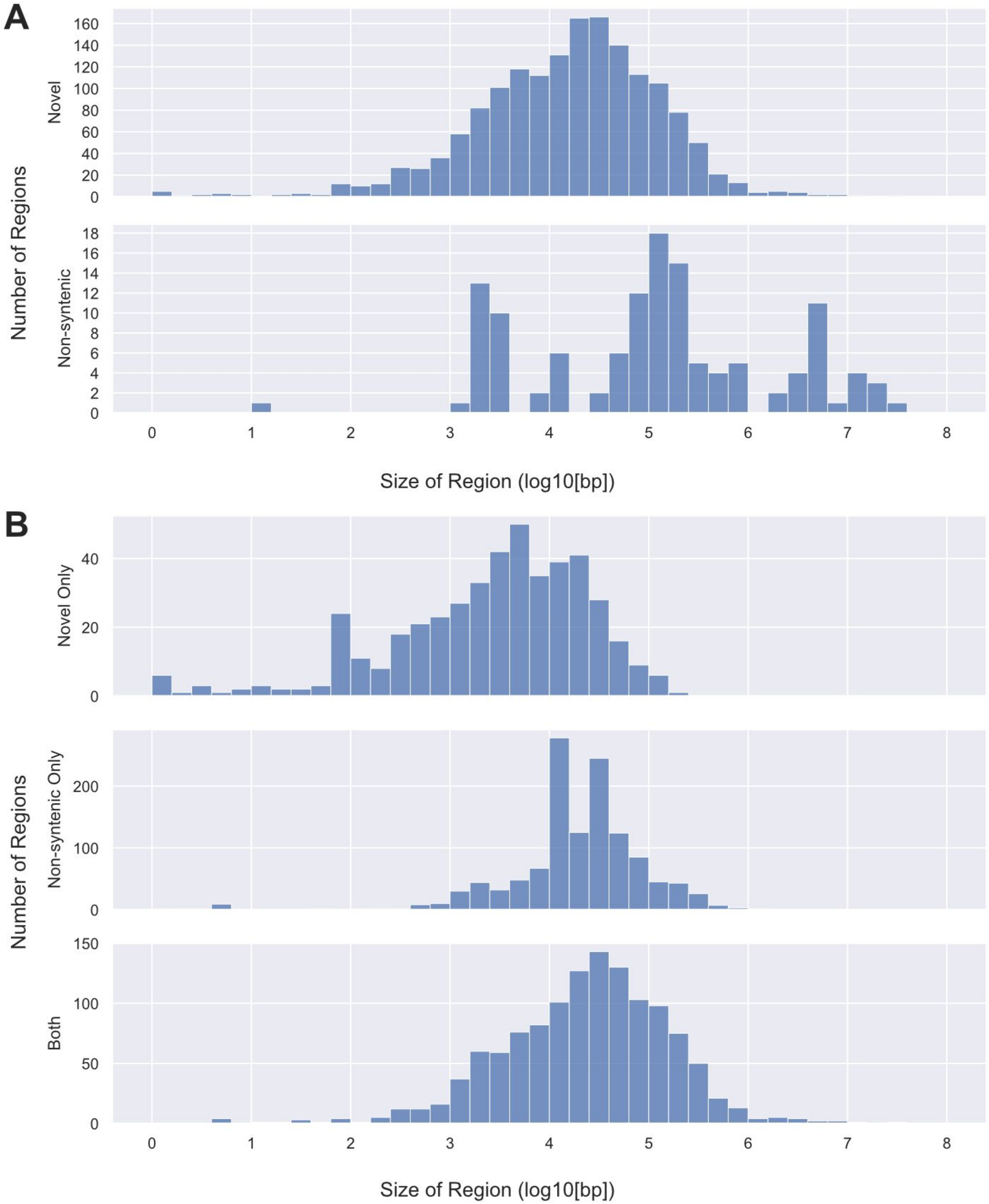


**Fig. S4.3. Distribution of region size for novel and non-syntenic regions**

**(A)** Distribution of region sizes for novel (top) and non-syntenic (bottom) regions as explored in Figure 4A-C. Sizes are shown in units of $\log_{10}$(bp). **(B)** Distribution of region sizes for non-overlapping novel (top) and non-syntenic (middle) regions, and regions annotated as both novel and non-syntenic (bottom) as explored in Figure 4D. Sizes are shown in units of $\log_{10}$(bp).

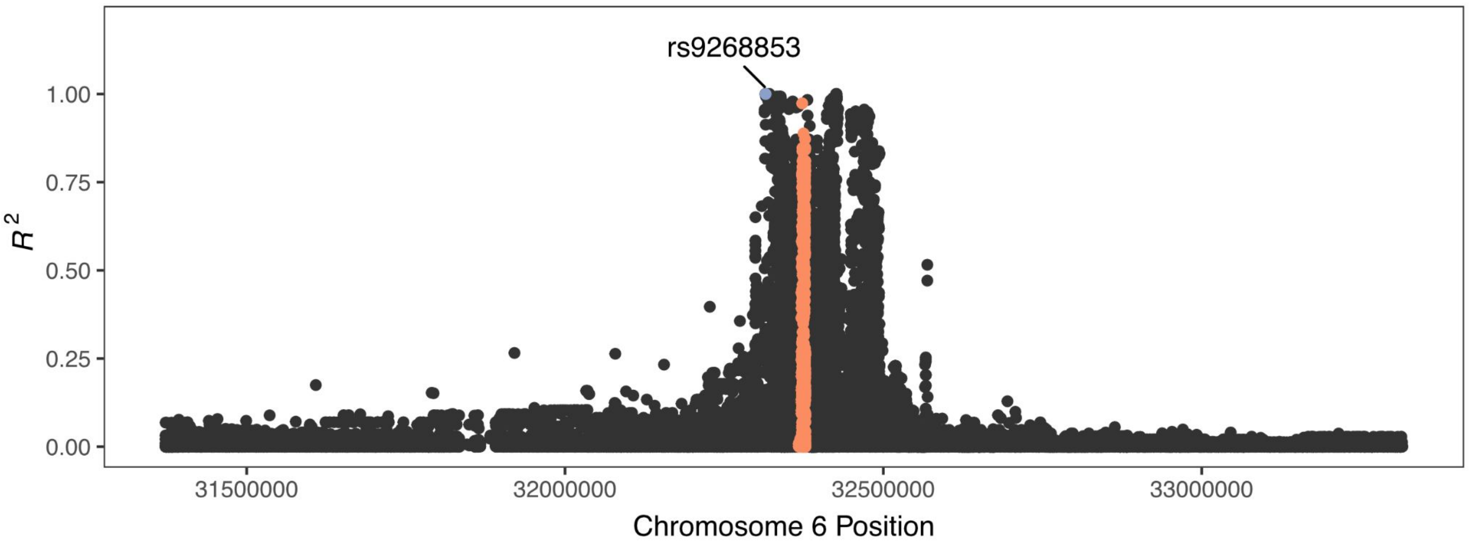


**Fig. S4.4. An example of pairwise LD between a known GWAS hit and novel variants in a non-syntenic region of T2T-CHM13.**

SNP rs9268853 (blue) is associated with fulminant type 1 diabetes in East Asian populations and segregates in strong LD ($R^2 > 0.8$) with 20 novel variants (orange) that were hidden to previous studies due to an insertion-deletion polymorphism that distinguishes GRCh38 from CHM13. While this example does not reflect an error or omission in GRCh38, it highlights the potential phenotypic and clinical relevance of such novel SNPs.

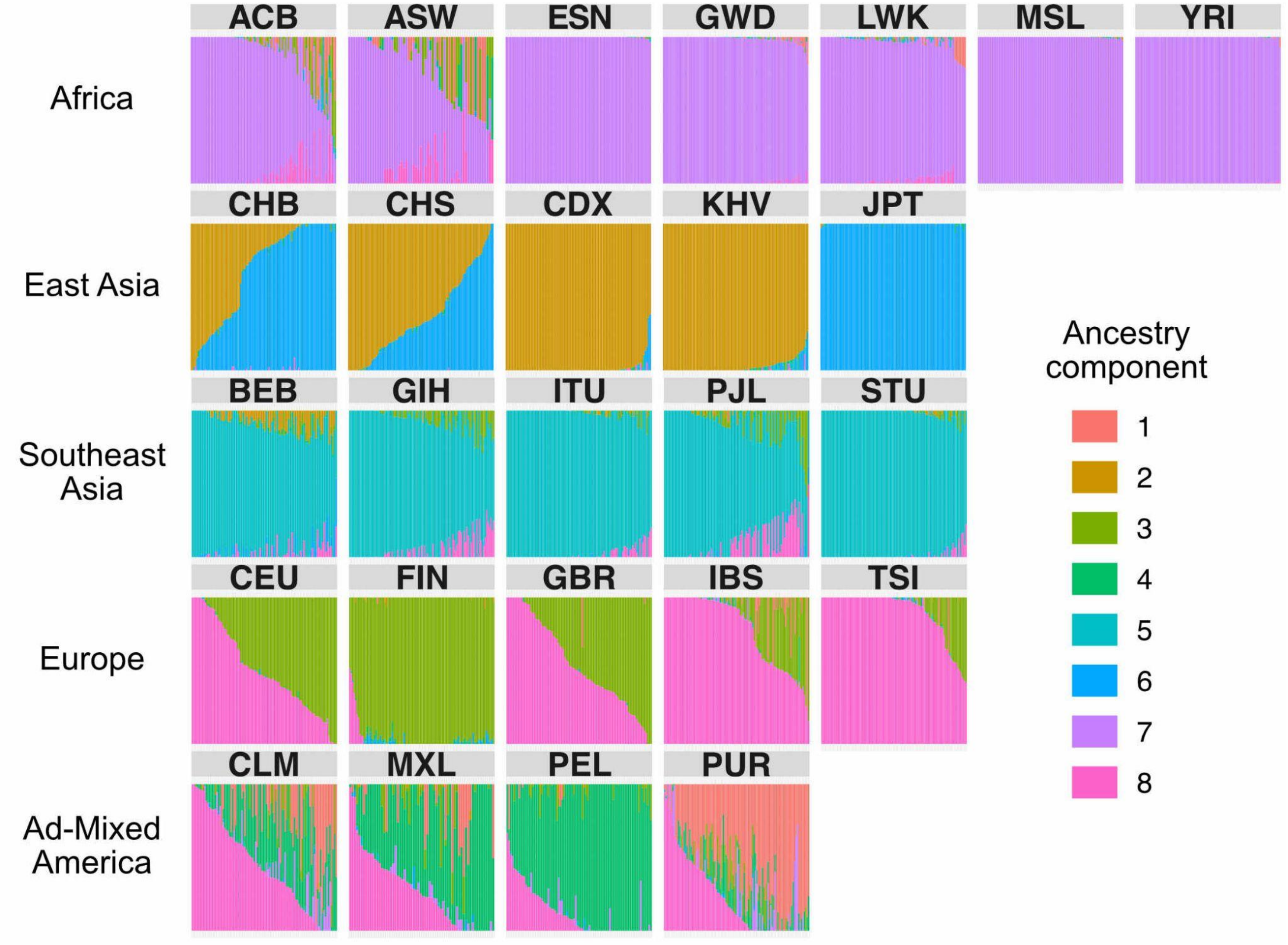


**Fig. S4.5. Inferred genome-wide ancestry of 1000 Genomes individuals, used to identify frequency-differentiated SNPs**

Admixture proportions ($k = 8$) for all samples in the 1000 Genomes dataset, inferred by Ohana. Vertical bars represent individual genomes and are grouped by population. Ohana models each individual as a combination of $k$ ancestry components and then searches for SNPs with evidence of frequency differentiation on these component lineages.

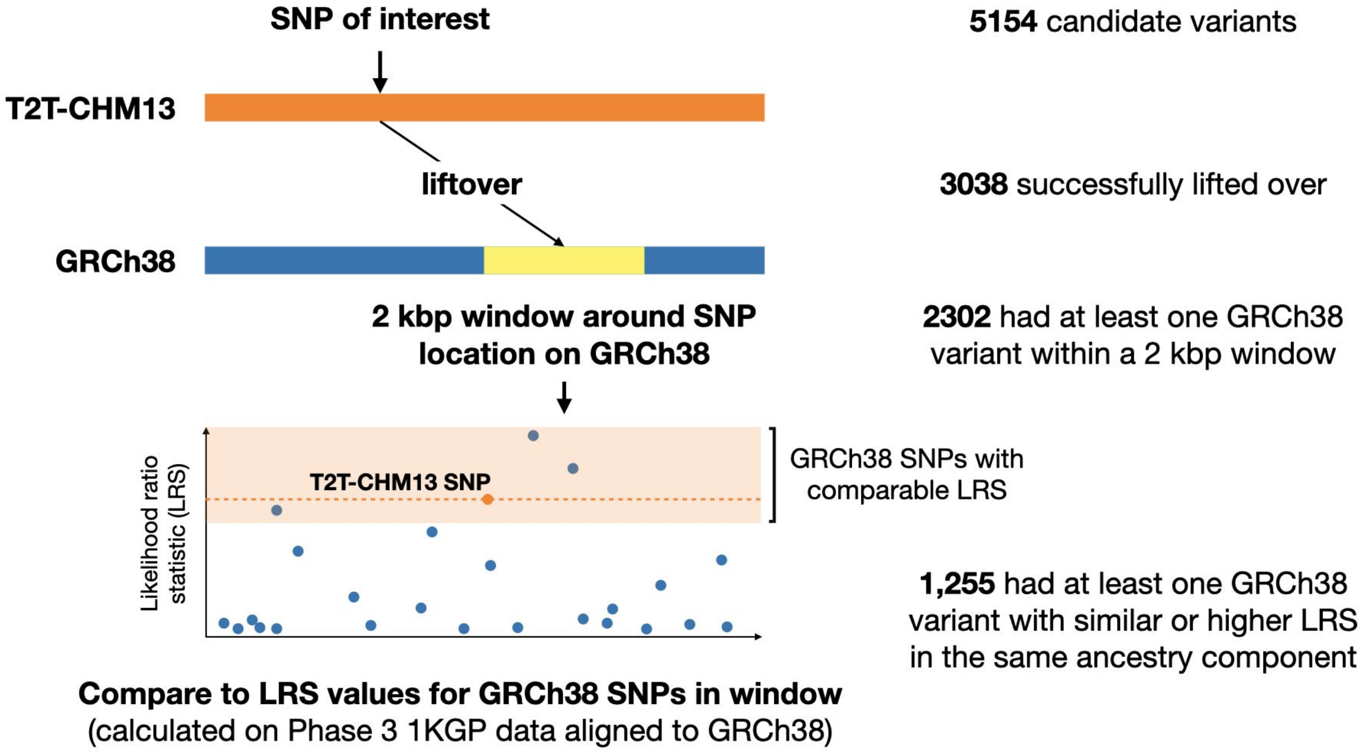


**Fig. S4.6. Comparing frequency-differentiated variants between T2T-CHM13 and GRCh38**

Schematic for comparing likelihood ratio statistic (LRS) values between variants called on T2T-CHM13 and variants called from the 1KGP Phase 3 data aligned to GRCh38. The 5,154 most highly frequency-differentiated variants across ancestry components were lifted over from T2T-CHM13 to GRCh38. We selected all GRCh38 variants within a 2 kbp window of the lifted over position. We considered the LRS score of a GRCh38 variant comparable to that of the T2T-CHM13 SNP if it was within 10 of the T2T-CHM13 LRS value or greater.

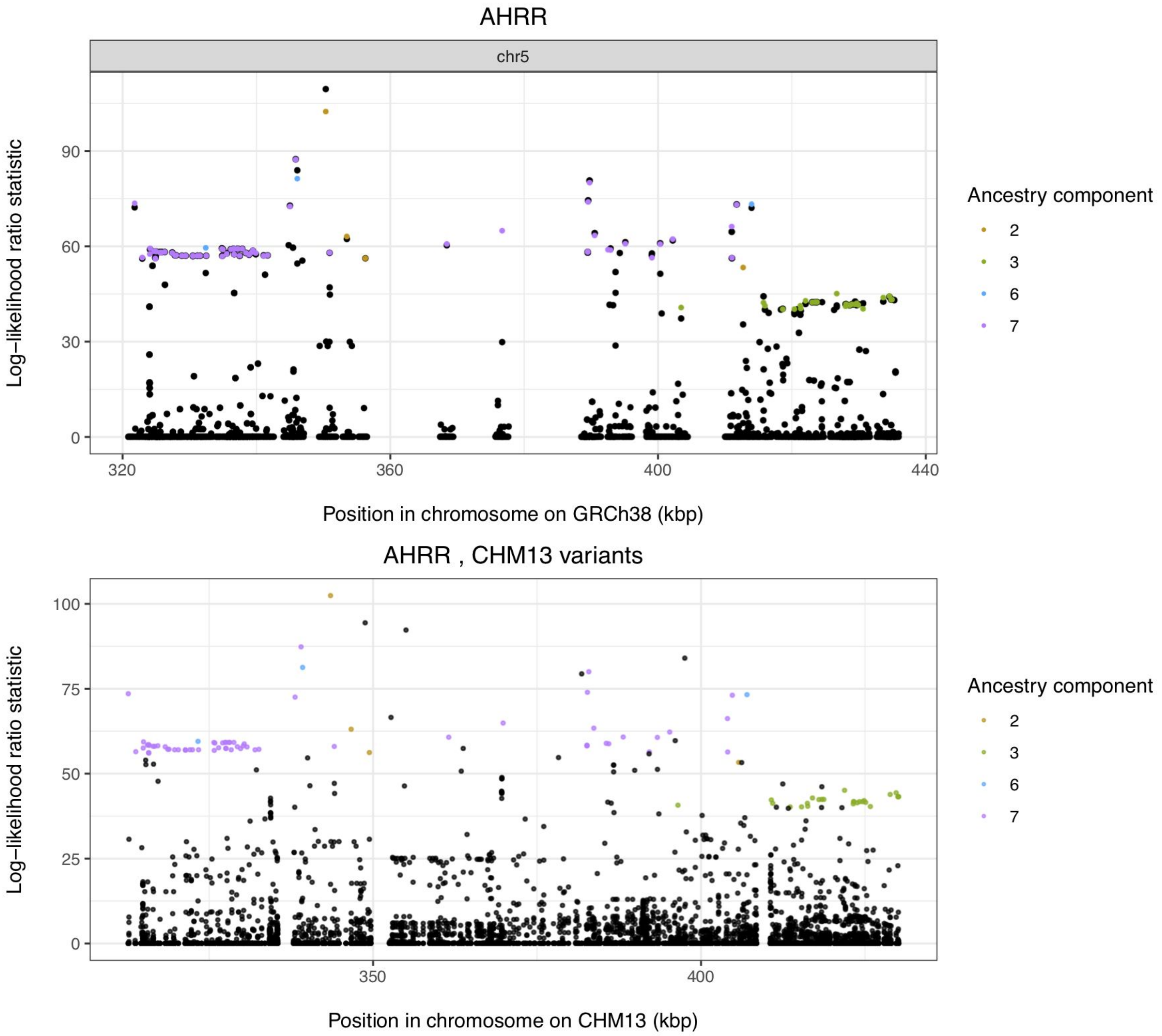


**Fig. S4.7. Example of a locus, overlapping the *AHRR* gene, with similar frequency differentiation results in T2T-CHM13 and GRCh38**

Likelihood ratio statistics (LRS) for variants overlapping the *AHRR* gene on GRCh38 (top plot) and T2T-CHM13 (bottom plot). T2T-CHM13 variants with outlier LRS values, indicating strong allele frequency differences between ancestry components, are colored by ancestry. In the top plot, colored points indicate T2T-CHM13 variants lifted over to GRCh38 and black points indicate variants called on GRCh38. In the bottom plot, black points indicate non-outlier T2T-CHM13 variants.

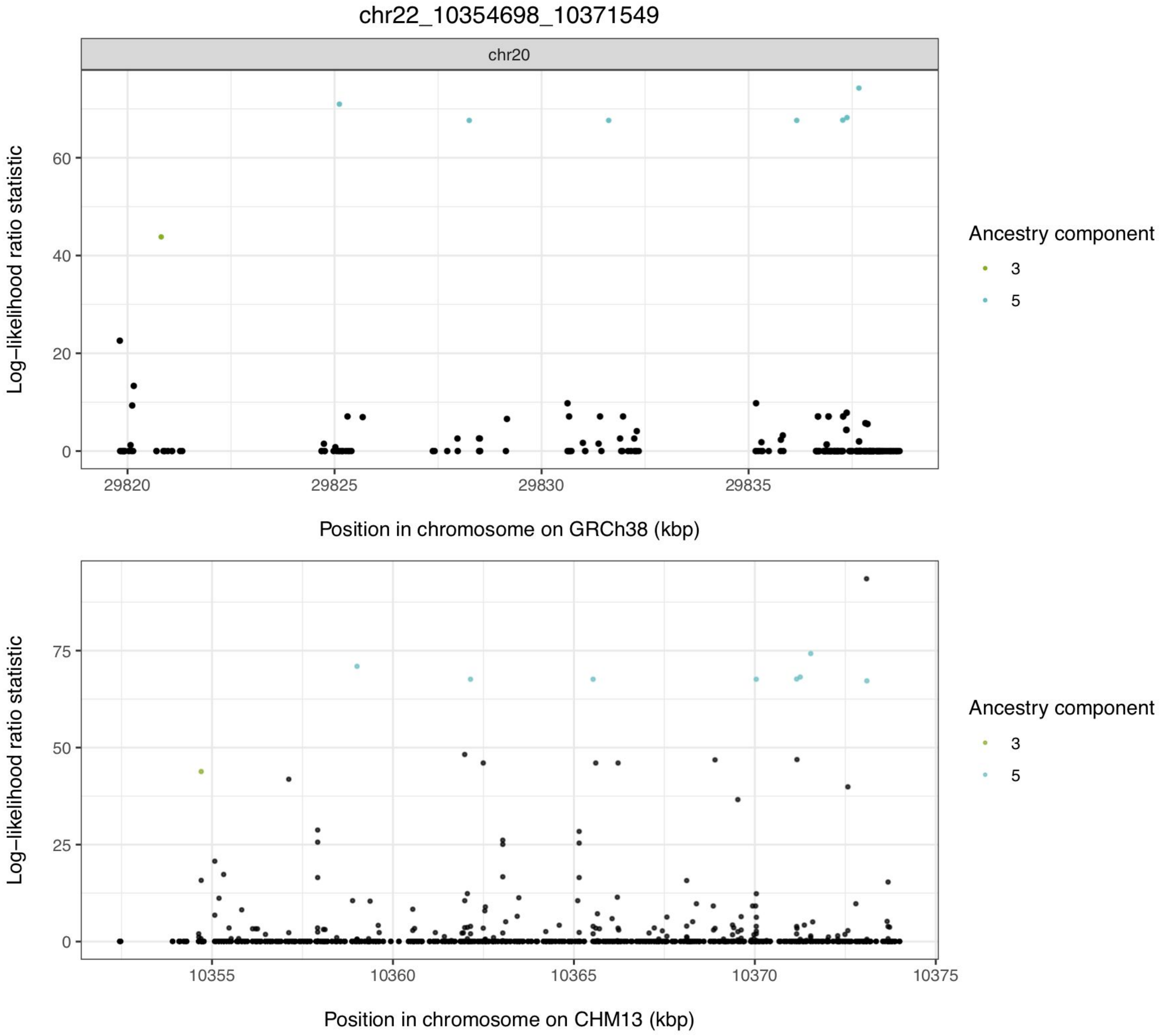


**Fig. S4.8. Example of a locus on chr22 with improved resolution in the T2T-CHM13 assembly**

Likelihood ratio statistics (LRS) for variants in a region on chr22 of T2T-CHM13 (bottom plot) and GRCh38 (top plot), where it lifts over to chr20. T2T-CHM13 variants with outlier LRS values, indicating strong allele frequency differences between ancestry components, are colored by ancestry. In the top plot, colored points indicate T2T-CHM13 variants lifted over to GRCh38 and black points indicate variants called on GRCh38. In the bottom plot, black points indicate non-outlier T2T-CHM13 variants.

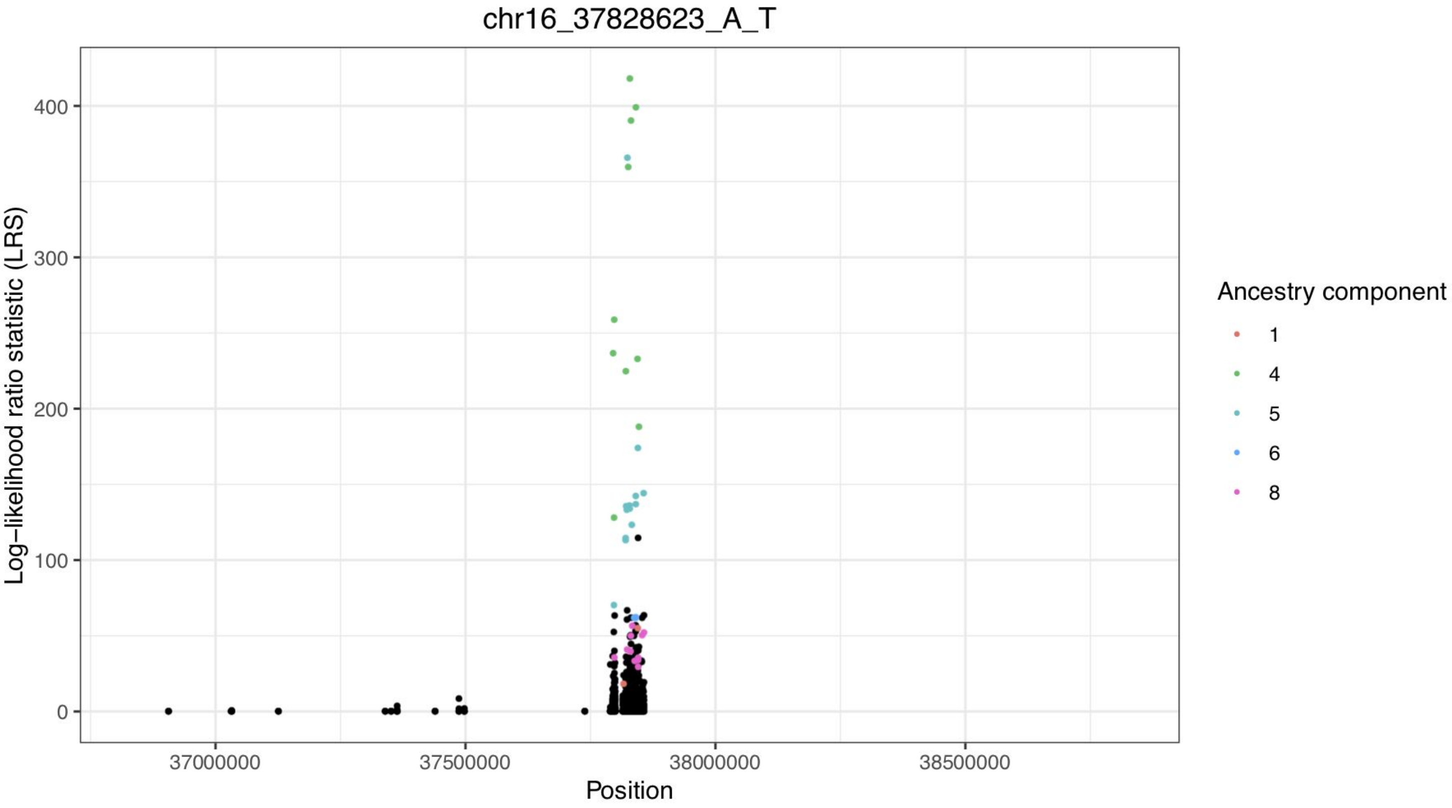


**Fig. S4.9. A frequency-differentiated novel locus at chr16:37828623**

Likelihood ratio statistics (LRS) for variants in a 2 Mbp window around chr16:37828623, a variant that reaches high allele frequencies in the Peruvian in Lima, Peru (PEL) population (ancestry component 4). T2T-CHM13 variants with outlier LRS values, indicating strong allele frequency differences between ancestry components, are colored by ancestry. Black points indicate non-outlier T2T-CHM13 variants.

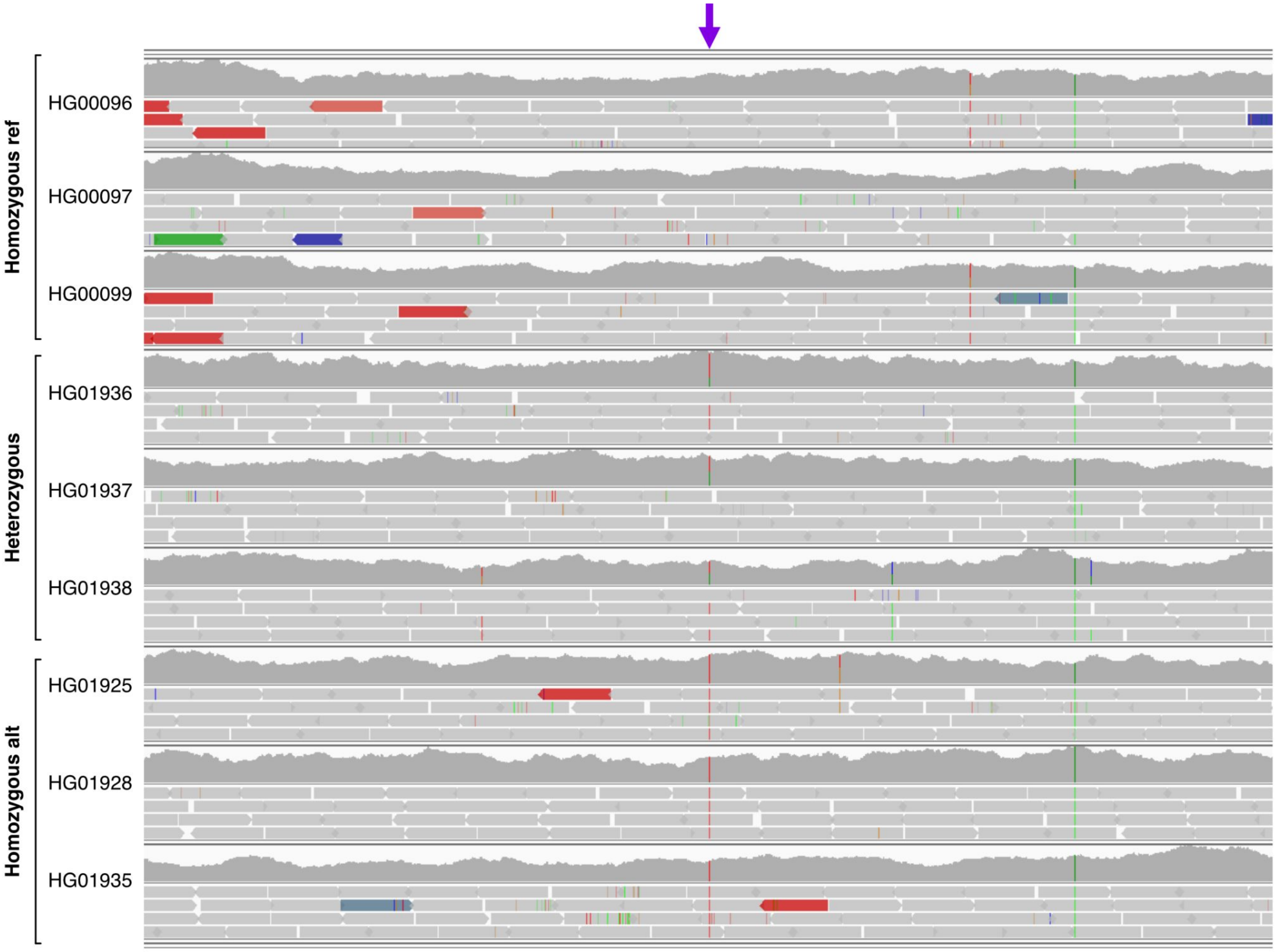


#### Fig. S4.10. Alignments to the region around chr16:37828623

Alignments to the region around an A -> T SNV (arrow) at chr16:37828623 on T2T-CHM13, for three homozygous reference (HG00096, HG00097, HG00099; European ancestry), heterozygous (HG01936, HG01937, HG01938; Peruvian in Lima, Peru [PEL] ancestry), and homozygous alternate (HG01925, HG01928, HG01935; PEL ancestry) individuals.

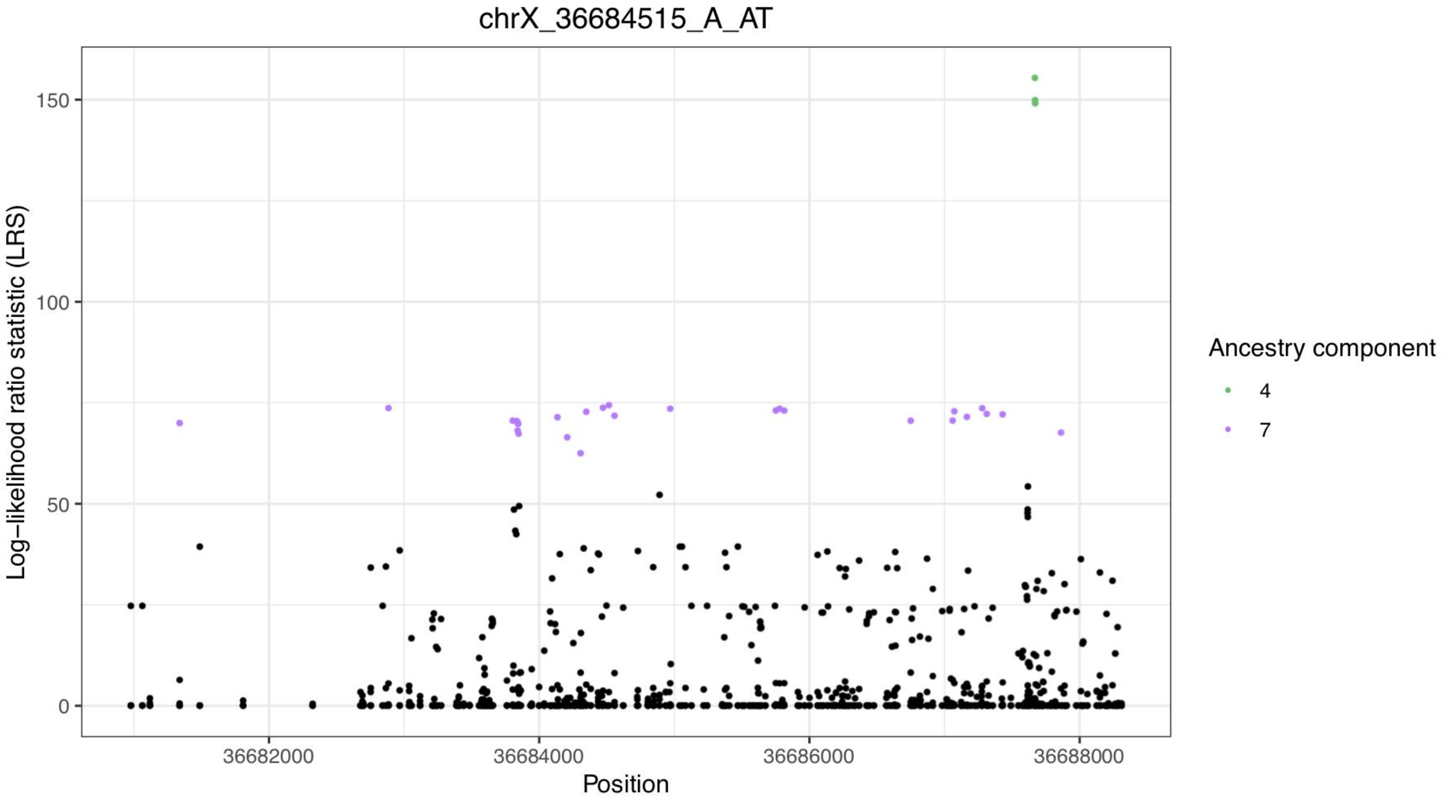


**Fig. S4.11. A frequency-differentiated novel locus at chrX:36684515**

Likelihood ratio statistics (LRS) for variants in a 2 Mbp window around chrX:36684515, a variant that reaches high allele frequencies in African populations (ancestry component 7). T2T-CHM13 variants with outlier LRS values, indicating strong allele frequency differences between ancestry components, are colored by ancestry. Black points indicate non-outlier T2T-CHM13 variants.

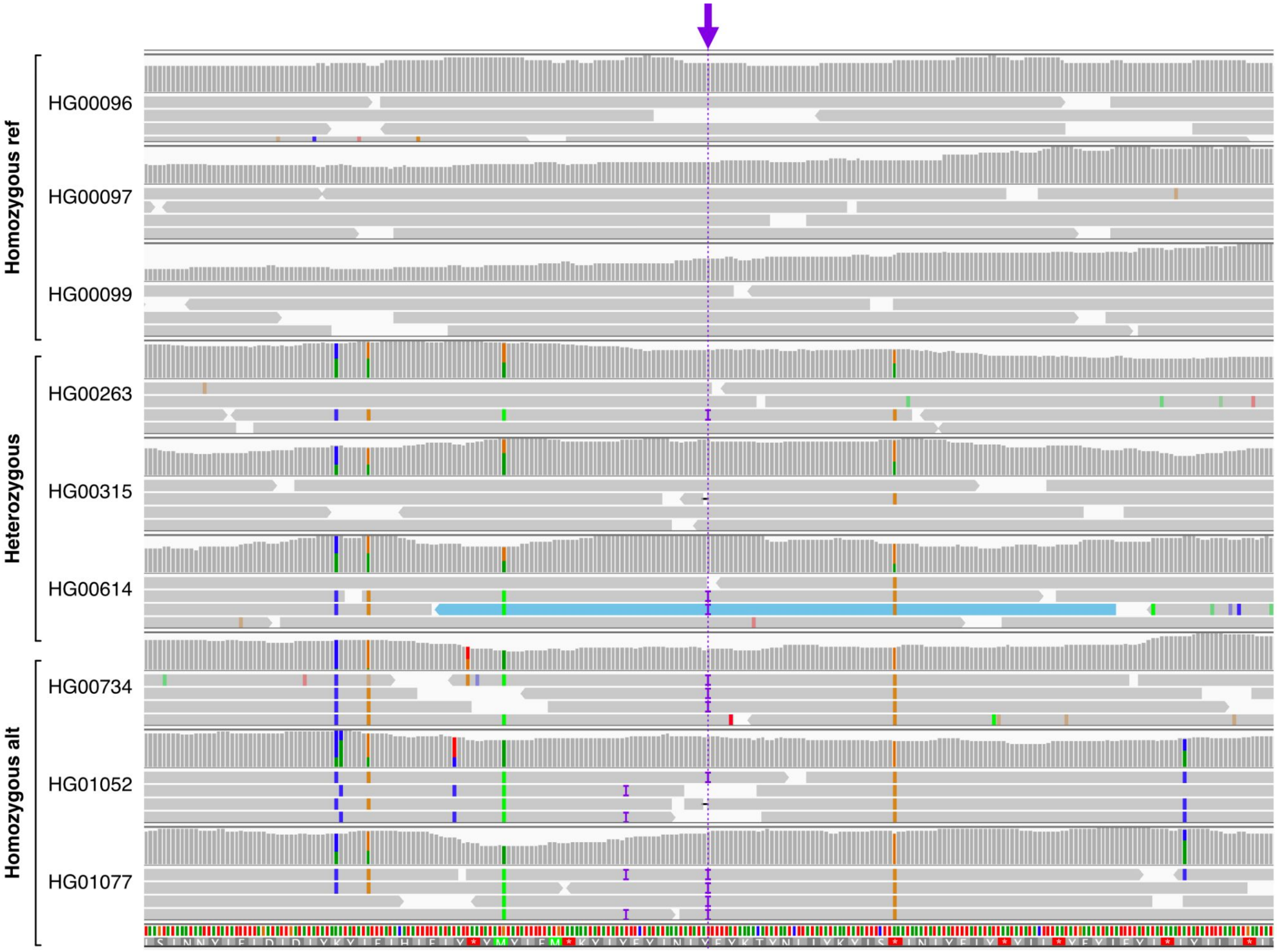


**Fig. S4.12. Alignments to the region around chrX:36684515**

Alignments to the region around a 1bp insertion (arrow, dashed line) at chrX:36684515 on T2T-CHM13, for three homozygous reference (HG00096, HG00097, HG00099; European ancestry), heterozygous (HG00263, HG00315, HG00614; European and East Asian ancestry), and homozygous alternate (HG00734, HG01052, HG01077; African ancestry) individuals.

### Figure 5. Supplemental

Impact of T2T-CHM13 on Clinical Genomics

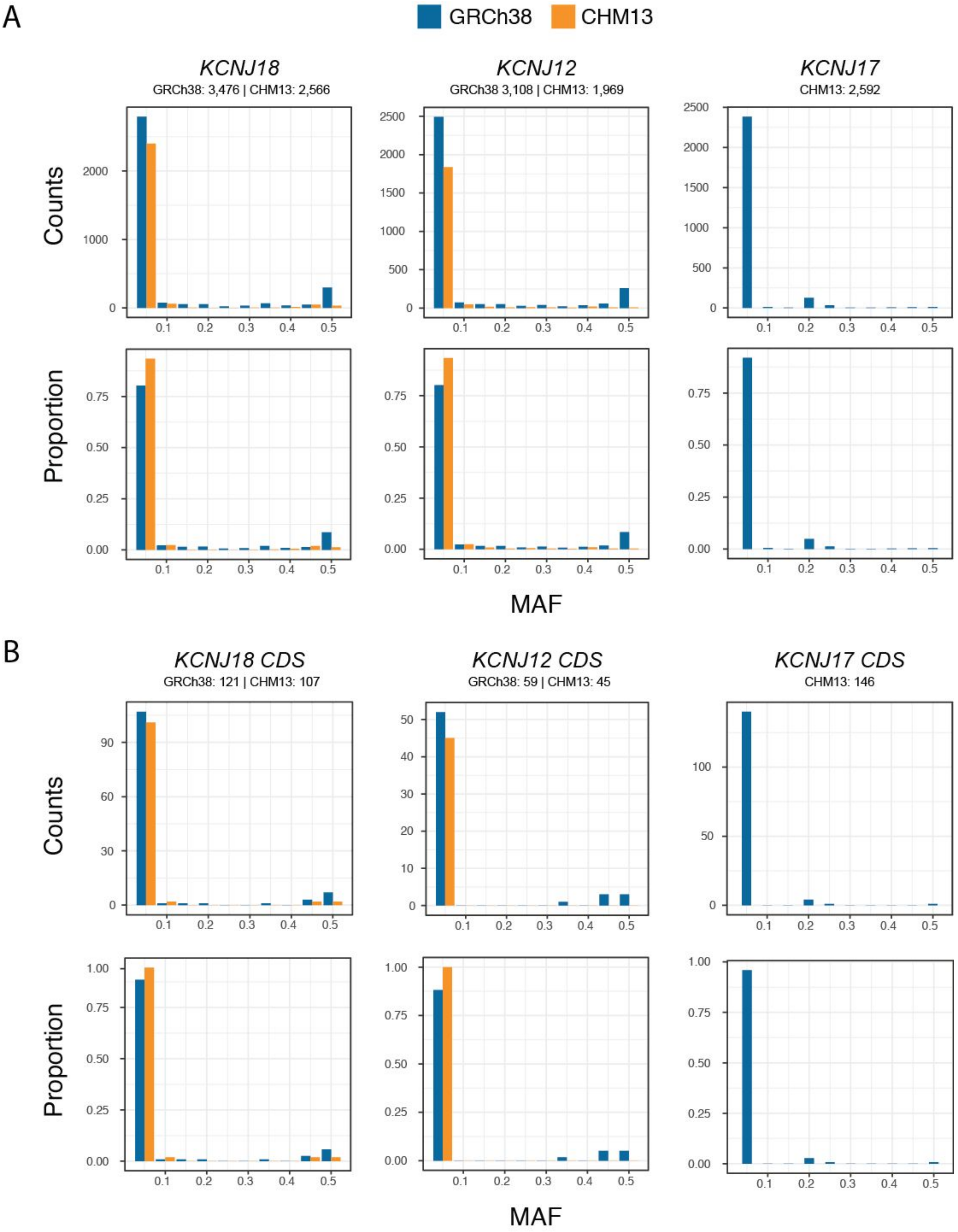


**Fig. S5.1. Folded site-frequency spectrum for *KCNJ18* paralogs in GRCh38 versus CHM13**

Total counts and proportions are depicted for varied minor-allele frequencies (MAF) of variants discovered within entire SDs (A) and CDS (B) in GRCh38 (blue) and CHM13 (orange) from 1KGP datasets. Total numbers of bi-allelic SNVs detected at each locus is indicated beneath each gene header.
